# Important Role of Hematopoietic Proteoglycan Serglycin in Liver Hepatocellular Carcinoma Associated with Tumor Microenvironment

**DOI:** 10.1101/2022.06.15.495916

**Authors:** Zengcheng Zou, Heping Xie, Wenhai Guo, Yue Li, Jiongshan Zhang, Yongwei Li

## Abstract

**Background:** Serglycin (SRGN) is a prominent hematopoietic proteoglycan that regulates tumorigenesis; however, its role in tumor immunity is unclear.

**Materials and methods:** We investigated the expression and prognostic potential of SRGN in liver hepatocellular cancer (LIHC) in the context of pan-cancer (for showing the similarity and heterogeneity) using the PrognoScan, GEPIA, Kaplan–Meier Plotter, and TIMER bioinformatics databases. HepG2 cells were transfected with an SRGN over-expression vector, and their proliferation, invasion, sorafenib resistance, and vasculature were examined *in vitro*. A subcutaneous xenograft tumor model was created in nude mice.

**Results:** SRGN expression was prominent in M2 macrophages in LIHC. The Kaplan–Meier Plotter indicated that SRGN RNA was a favorable prognostic factor after correcting for clinical factors. TIMER 2.0 showed that the immune infiltrates of CD8+ T cells, M1 and M2 macrophages, and endothelial cells were strongly correlated with SRGN RNA expression (r=0.552, P=5.79e^-29^; r=0.517, P=5.84e^-25^; r=0.696, P=3.26e^51^; and r=0.522, P=1.67e^-25^, respectively), and had prognostic potential in LIHC in patients with low or high levels of SRGN, in addition to resting memory CD4+ T cells, hematopoietic stem cells (HSCs), and myeloid-derived suppressor cells (MDSCs). SRGN promoted the proliferation of HepG2 cells *in vitro* and *in vivo*, and was associated with weak sorafenib resistance, invasion, and vasculature. CD206 and CD80 were up-regulated and down-regulated, respectively, in subcutaneous tumor tissues.

**Conclusions:** The results comprehensively revealed relationships between SRGN and tumor microenvironment(TME)-infiltrating cells, especially monocyte/macrophage subsets. These may constitute an important TME because the pro-tumorigenicity of SRGN in liver cancer.

## Introduction

It is estimated that 296 million people are infected with hepatitis B virus (HBV) worldwide [1]. HBV has a broad spectrum of clinical manifestations ranging from a healthy carrier state to acute and chronic hepatitis (chronic hepatitis B), liver cirrhosis, primary liver cancer (PLC, mainly liver hepatocellular carcinoma, LIHC), and end-stage liver disease. Most patients recover from acute HBV infection, but 1%–5% of patients develop LIHC [2]. HBV-specific B-cell antibodies and HBV-specific T cells combat HBV infection [3]. The HBV infection-based microenvironment consists of immunosuppressive and inflammatory cytokines and immune cells, including macrophages, neutrophils, and lymphocytes [4–6], which can complicate the LIHC tumor microenvironment (TME). Immunotherapy with PD-1/PD-L1 blockade has been regarded as a hot topic and promising therapy in LIHC [4]. However, there is a need to clarify the mechanisms linking immunity and malignancy to investigate new treatments for LIHC. Serglycin (SRGN, gene *SRGN*) is a prominent hematopoietic proteoglycan with inflammatory and malignant properties, which interacts with proteases, chemokines, and cytokines and is required for the formation of secretory granules [7–9].

The HepG2.2.15 cell line is derived from HepG2 human hepatoma cells chronically infected with HBV [10]. We previously analyzed the Human Genome U133 Plus 2.0 Array (Platform GPL570; Affymetrix, Santa Clara, CA) and revealed extensive changes in global gene transcription in HepG2.215 compared with HepG2 cells, with SRGN being the most significantly differentially expressed gene [11]. Levels of SRGN detected by flow cytometry were increased in peripheral monocytes, neutrophils, and lymphocytes in HBV-infected LIHC patients compared with healthy adults, and hematopoietic SRGN was associated with an unfavorable prognosis in terms of overall survival (OS) [12]. Based on these results, we comprehensively evaluated the prognostic potential of SRGN in pan-cancer using bioinformatics databases and further investigated its role in LIHC. Other PLC such as cholangio carcinoma (CHOL) is rarely mentioned in these databases.

## Materials and methods

### SRGN expression, prognosis analysis and correlation analysis

We examined the differential expression and prognosis of *SRGN* RNA in pan-cancer using Gene Expression Profiling Interactive Analysis (GEPIA, http://gepia.cancer-pku.cn/index.html, http://gepia2021.cancer-pku.cn/)[13], Kaplan–Meier plotter databases (http://kmplot.com/analysis/ (registration-freeKM-plotter))[14], the Tumor Immune Estimation Resource (TIMER, https://cistrome.shinyapps.io/timer/, http://timer.comp-genomics.org/)[15], and PrognoScan database (http://www.abren.net/PrognoScan/) [16], respectively. SRGN protein expression was analyzed by Clinical Proteomic Tumor Analysis Consortium (CPTAC) analysis. The expression of SRGN in immune cells was analyzed using GEPIA, TIMER and The Human Protein Atlas (HPA).

The correlation module was used to investigate correlations between SRGN and TME cells or gene markers using Spearman’s correlation analysis via TIMER. Spearman’s correlation was determined using the following criteria: 0.1–0.3: weak correlation, 0.3–0.5: moderate correlation, and 0.5–1.0: strong correlation [17]. The results were represented as a heatmap for pan-cancer and scatter plots for LIHC, respectively. SRGN expression in liver cell types was analyzed using HPA, SRGN–protein interaction was analyzed by STRING, and SRGN-associated genes in LIHC were analyzed using the University of Alabama at Birmingham Cancer data (UALCAN).

### Over-expression of SRGN in HepG2 cells

Details of the methods, including construction of an over-expression vector, lentiviral transduction, quantitative polymerase chain reaction, western blotting, Cell Counting Kit-8 cell viability assay, Transwell invasion assay, angiogenesis, vasculogenic mimicry, immunofluorescence, and hematoxylin and eosin staining are provided in the Supplementary Methods.

### Animal experiments

Male BALB/c nude mice (5-weeks-old, weight 18-20g) were purchased from Guangzhou University of Chinese Medicine Laboratory Animal Center (Guangzhou, China). Tumor cells (5×10^6^/0.1 mL phosphate-buffered saline) were injected randomly into the right axillary region and the mice were monitored twice a week for 4 weeks for tumor formation, then, sacrificed by carbon dioxide asphyxiation. After animal experiments, the carcasses were returned to the Laboratory Animal Center for harmless treatment. The study is reported in accordance with ARRIVE guidelines (https://arriveguidelines.org). The animal experiments were approved by Institutional Animal Care and Use Committee of Rulge Biotechnology (ethical approval number, 20230201002).

### Statistical analysis

Measured data were compared by *t*-tests or ANOVA using SPSS 22.0 (IBM Corp. Armonk, NY, USA) and counted data were compared with χ^2^ tests. Differences between means for data with a skewed distribution and variance were analyzed by rank sum tests. P < 0.05 was considered statistically significant.

## Results

### Expression of SRGN in pan-cancer and cell types

The differential expression of SRGN RNA between pan-cancer and adjacent normal tissues was analyzed using TIMER (Figure 1). SRGN levels were significantly higher in head and neck cancer (HNSC) and kidney renal clear cell carcinoma compared with adjacent normal tissues. SRGN expression was also higher in human papilloma virus (HPV)-positive compared with HPV-negative HNSC. SRGN levels were significantly lower in bladder urothelial carcinoma, breast invasive carcinoma (BRCA), colon adenocarcinoma, kidney chromophobe, kidney renal papillary cell carcinoma, LIHC, lung squamous cell carcinoma, lung adenocarcinoma (LUAD), prostate adenocarcinoma, rectum adenocarcinoma, and uterine corpus endometrial carcinoma than in adjacent normal tissues, and was lower in skin cutaneous melanoma (SKCM) tumor than in SKCM metastasis. (Figure 1A). KM-plotter analysis showed that SRGN mRNA levels differed significantly between tumor and adjacent normal tissues in 19 cancers, except for adrenal, liver, and renal_PA cancers (Supplementary Figure 1). However, KM-plotter analysis of GeneChip data showed that SRGN levels were significantly higher in adjacent normal tissues than in LIHC tumor tissues (non-paired tumor vs. adjacent normal tissues, P = 5.85e^-11^; paired tumor *vs.* adjacent normal tissues, P = 9.29e^-10^), which was consistent with the comparison among LIHC tumor, adjacent normal, and metastasis tissues (P = 7.92e^-11^; Figure 2A). There were no differences in SRGN levels based on RNA-Seq data between LIHC tissues and adjacent normal tissues as in GEPIA (Figure 2B, Supplementary Figure 2). The results for SRGN protein levels in tumor and normal samples were consistent with the RNA levels for breast cancer, lung cancer, and HNSC by CPTAC (Figure 1B), but there were no data for SRGN protein in liver cancer. Among all these cancers, SRGN levels were highest in patients with acute myeloid leukemia (AML) according to the TIMER, GEPIA, and KM-plotter databases.

**Figure 1.**
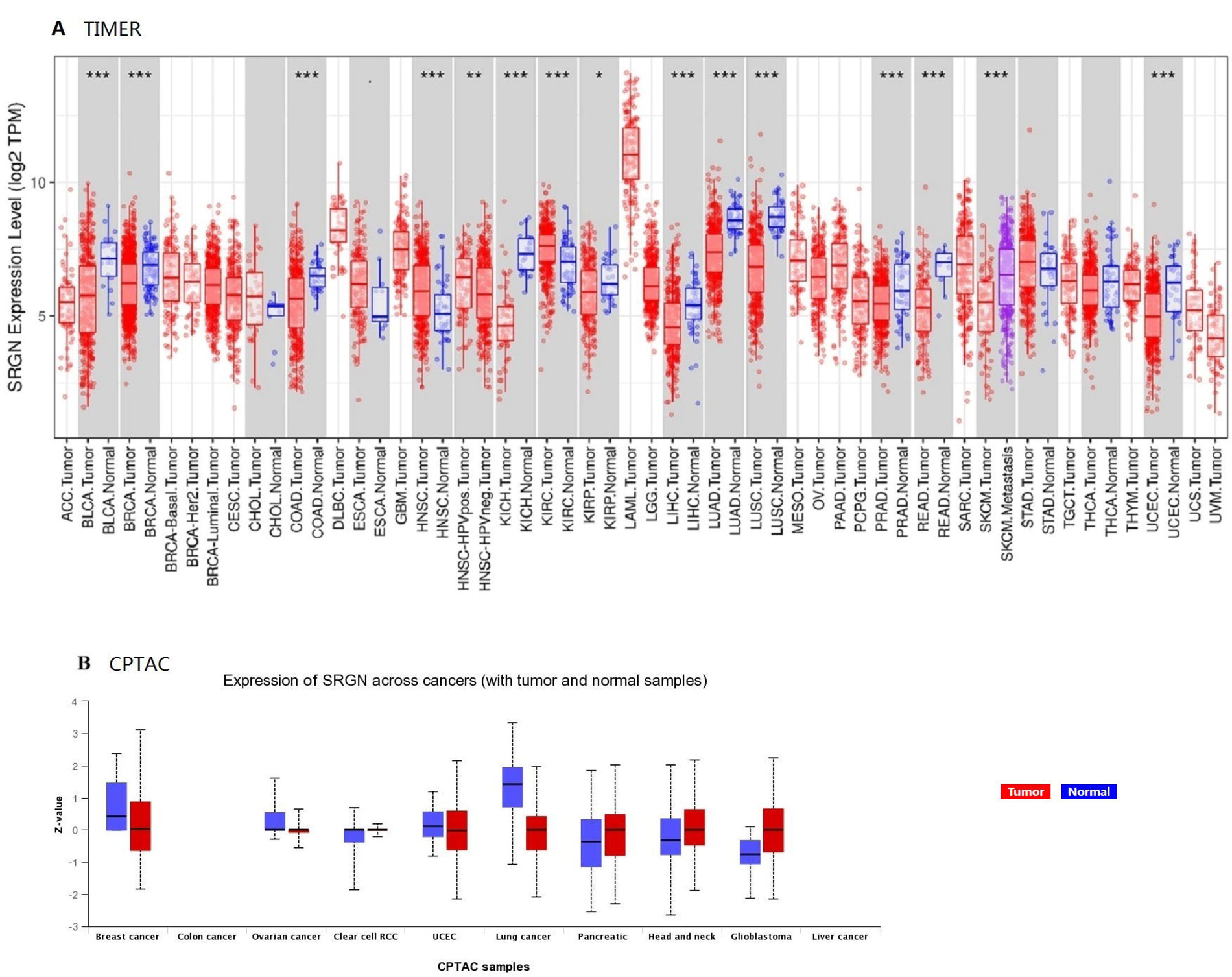
(A) Differential expression of SRGN in tumor (red column) compared with adjacent normal tissues (blue column) across all TCGA tumors by Wilcoxon’s test using TIMER (Diff Exp module) (*P < 0.05, **P < 0.01, ***P < 0.001). (B) The University of ALabama at Birmingham CANcer data analysis (UALCAN) portal provides protein expression analysis using data from the Clinical Proteomic Tumor Analysis Consortium (CPTAC). Protein expression data for liver cancer are not available.

**Figure 2.**
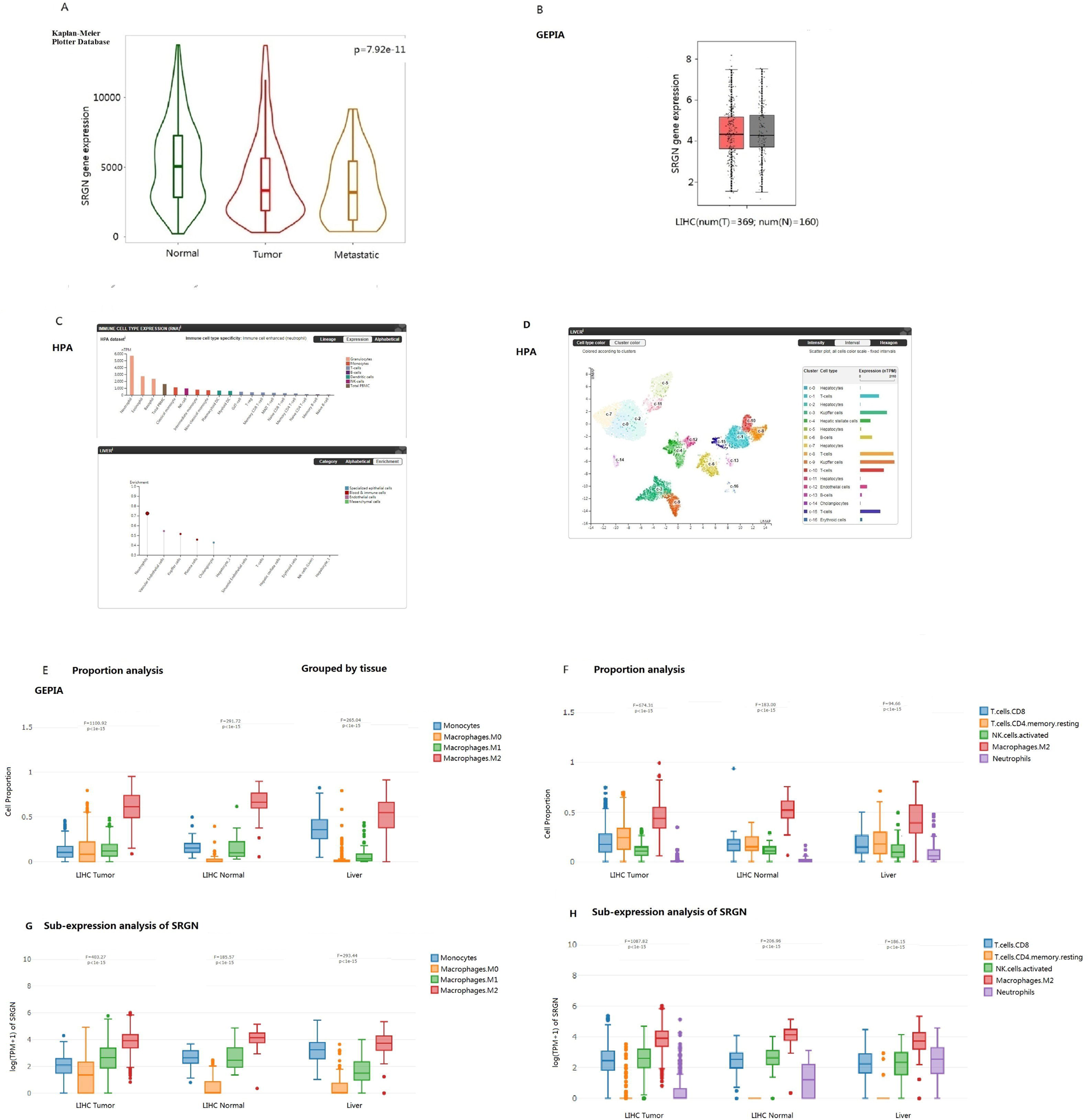
(A) SRGN gene expression in tumor, adjacent normal, and metastatic liver cancer tissues via the Kaplan–Meier plotter database. (B) Expression of SRGN RNA in LIHC and adjacent normal tissues via GEPIA. (C) SRGN RNA was expressed in immune cell types in the order neutrophils > eosinophils >basophils > total peripheral blood mononuclear cells > classical monocytes, and in liver cell types in the order neutrophils >endothelial cells > Kupffer cells > plasma cells > cholangiocytes via The Human Protein Atlas (HPA). (D) SRGN RNA in liver cells via HPA. SRGN was prominently expressed in T cells, Kupffer cells, B cells, hepatic stellate cells, and endothelial cells, and barely expressed in hepatocytes. (E, F) GEPIA proportion analysis. The proportion of each cell type shown with interactive boxplots by ANOVA test among cell types or TCGA/GTEx sub-datasets. (G, H) GEPIA sub-expression analysis. SRGN expression in each cell type shown with interactive boxplot by ANOVA test. All cell types were analyzed, but only cells with strong correlations with SRGN or prognostic potential are shown.

HPA analysis showed that SRGN RNA was expressed in immune cell types in the order neutrophils > eosinophils > basophils > total peripheral blood mononuclear cells > classical monocytes, and in liver cell types in the order neutrophils > vascular endothelial cells (ECs) > Kupffer cells > plasma cells > cholangiocytes. SRGN was prominently expressed in T cells, Kupffer cells, B cells, hepatic stellate cells, and ECs, but was rarely expressed in hepatocytes (Figure 2C, D). However, the proportion and sub-expression analysis via GEPIA showed that the proportion of M2 macrophages and sub-expression of SRGN in M2 were highest in LIHC, adjacent normal, and liver tissues, respectively (Figure 2E-H).

### Prognostic potential of SRGN in pan-cancer according to PrognoScan, GEPIA, Kaplan–Meier Plotter Databases, and TIMER

We analyzed the relationship between SRGN expression and prognosis in pan-cancer (Supplementary Figures 3–6, Table 1). PrognoScan analysis revealed that SRGN expression had a significant impact on prognosis in five cancer types: AML, diffuse large B-cell lymphoma (DLBCL), BRCA, lung cancer, and ovarian cancer (OV). High SRGN expression was associated with a poor prognosis in AML and DLBCL, but showed favorable prognostic potential in a small sample size of three solid tumors, BRCA, lung cancer and OV. According to GEPIA, SRGN only had prognostic value in testicular germ cell tumors, glioblastoma multiforme, and SKCM. According to the TIMER gene_outcome module, SRGN was a significant prognostic factor for OS in patients with sarcoma, SKCM, and SKCM metastasis after correcting for age, race, sex, and tumor purity (Supplementary Figure 5, Table 1).

**Table 1.**
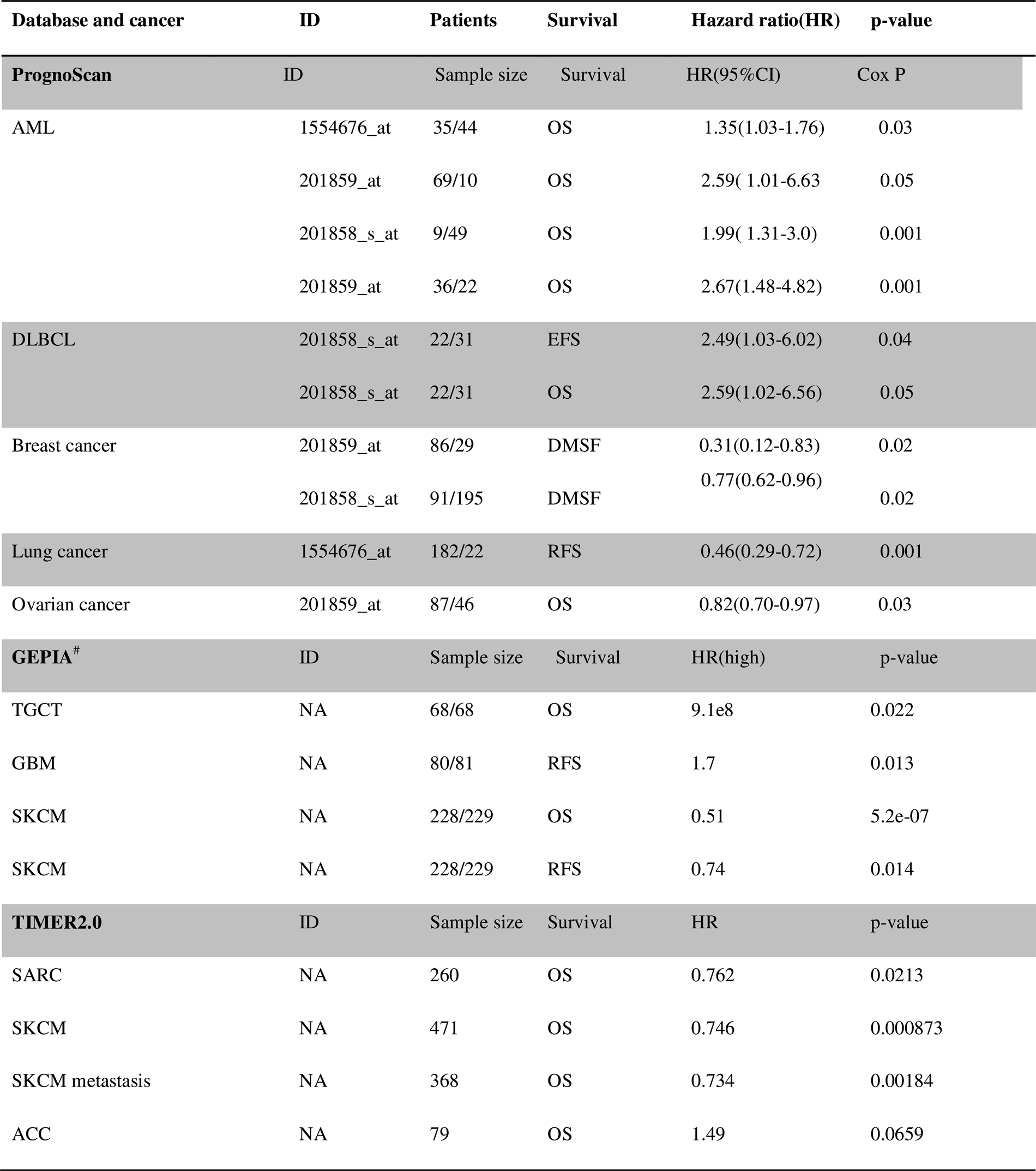
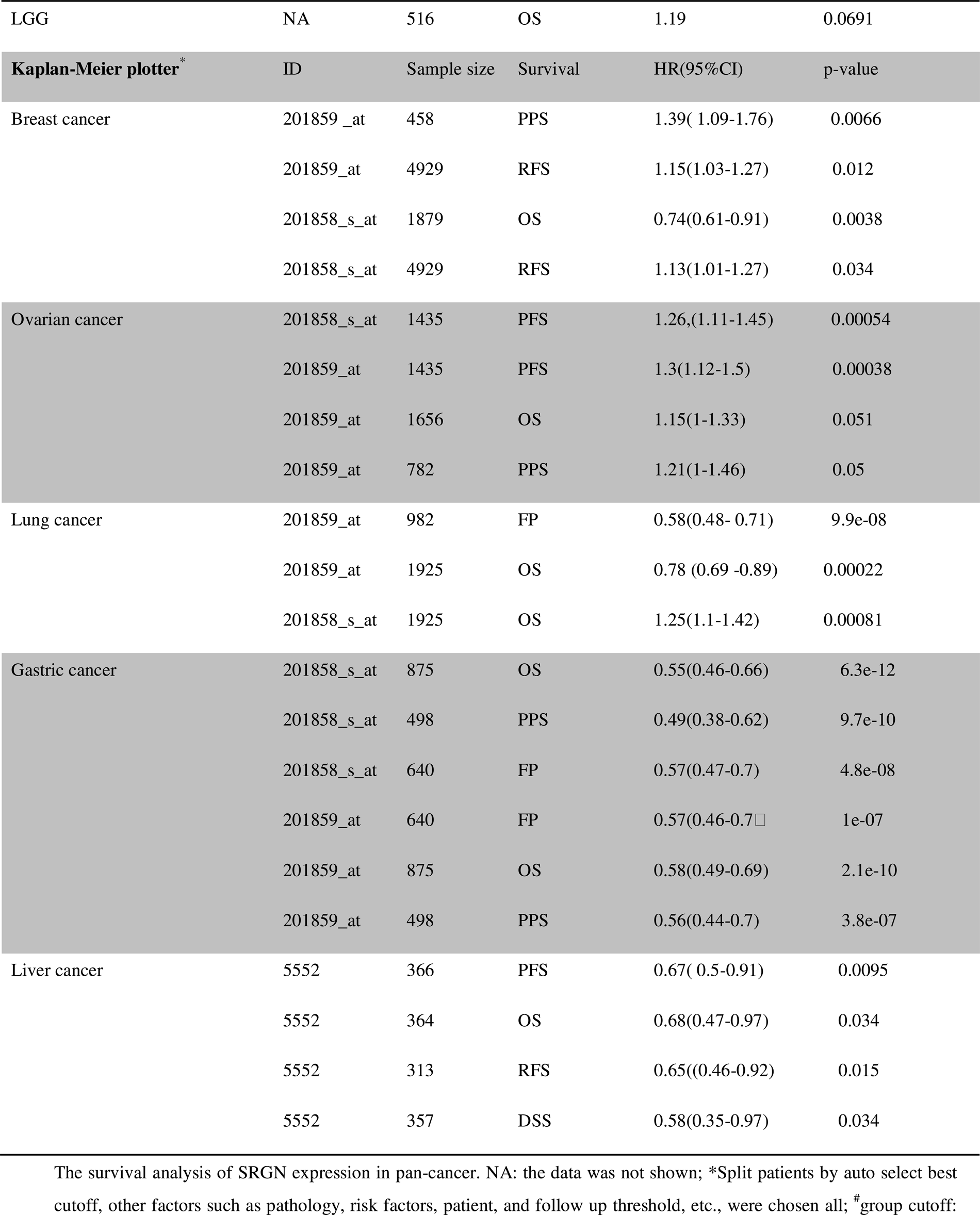
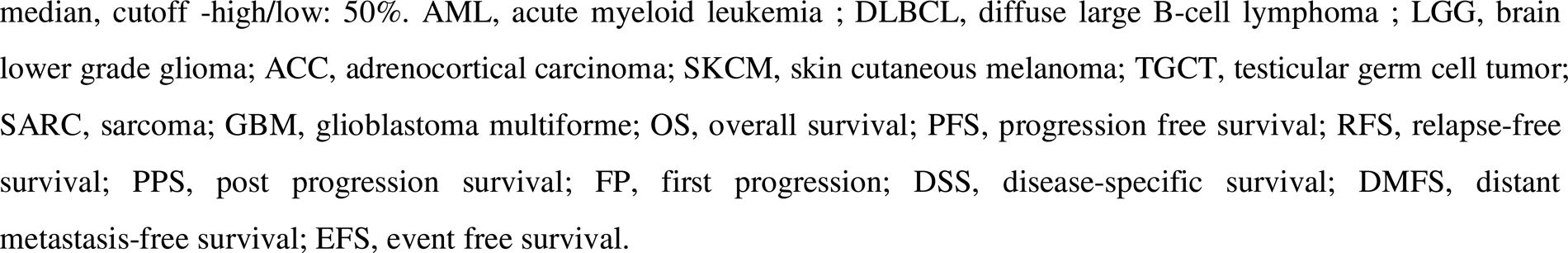
The correlation of serglycin expression and prognosis in cancers via four bioinformatics databases.

| Database and cancer | ID | Patients | Survival | Hazard ratio(HR) | p-value |
| --- | --- | --- | --- | --- | --- |
| PrognScan | ID | Sample size | Survival | HR(95%CI) | Cox P |
| AML | 1554676_at | 35/44 | OS | 1.35(1.03-1.76) | 0.03 |
|  | 201859_at | 69/10 | OS | 2.59( 1.01-6.63) | 0.05 |
|  | 201858_s_at | 9/49 | OS | 1.99( 1.31-3.0) | 0.001 |
|  | 201859_at | 36/22 | OS | 2.67(1.48-4.82) | 0.001 |
| DLBCL | 201858_s_at | 22/31 | EFS | 2.49(1.03-6.02) | 0.04 |
|  | 201858_s_at | 22/31 | OS | 2.59(1.02-6.56) | 0.05 |
| Breast cancer | 201859_at | 86/29 | DMSF | 0.31(0.12-0.83) | 0.02 |
|  | 201858_s_at | 91/195 | DMSF | 0.77(0.62-0.96) | 0.02 |
| Lung cancer | 1554676_at | 182/22 | RFS | 0.46(0.29-0.72) | 0.001 |
| Ovarian cancer | 201859_at | 87/46 | OS | 0.82(0.70-0.97) | 0.03 |
| GEPIA<sup>#</sup> | ID | Sample size | Survival | HR(high) | p-value |
| TGCT | NA | 68/68 | OS | 9.1e8 | 0.022 |
| GBM | NA | 80/81 | RFS | 1.7 | 0.013 |
| SKCM | NA | 228/229 | OS | 0.51 | 5.2e-07 |
| SKCM | NA | 228/229 | RFS | 0.74 | 0.014 |
| TIMER2.0 | ID | Sample size | Survival | HR | p-value |
| SARC | NA | 260 | OS | 0.762 | 0.0213 |
| SKCM | NA | 471 | OS | 0.746 | 0.000873 |
| SKCM metastasis | NA | 368 | OS | 0.734 | 0.00184 |
| ACC | NA | 79 | OS | 1.49 | 0.0659 |
| LGG | NA | 516 | OS | 1.19 | 0.0691 |
| **Kaplan-Meier plotter<sup>\*</sup>** | **ID** | **Sample size** | **Survival** | **HR(95%CI)** | **p-value** |
| Breast cancer | 201859_at | 458 | PPS | 1.39( 1.09-1.76) | 0.0066 |
|  | 201859_at | 4929 | RFS | 1.15(1.03-1.27) | 0.012 |
|  | 201858_s_at | 1879 | OS | 0.74(0.61-0.91) | 0.0038 |
|  | 201858_s_at | 4929 | RFS | 1.13(1.01-1.27) | 0.034 |
| Ovarian cancer | 201858_s_at | 1435 | PFS | 1.26,(1.11-1.45) | 0.00054 |
|  | 201859_at | 1435 | PFS | 1.3(1.12-1.5) | 0.00038 |
|  | 201859_at | 1656 | OS | 1.15(1-1.33) | 0.051 |
|  | 201859_at | 782 | PPS | 1.21(1-1.46) | 0.05 |
| Lung cancer | 201859_at | 982 | FP | 0.58(0.48- 0.71) | 9.9e-08 |
|  | 201859_at | 1925 | OS | 0.78 (0.69 -0.89) | 0.00022 |
|  | 201858_s_at | 1925 | OS | 1.25(1.1-1.42) | 0.00081 |
| Gastric cancer | 201858_s_at | 875 | OS | 0.55(0.46-0.66) | 6.3e-12 |
|  | 201858_s_at | 498 | PPS | 0.49(0.38-0.62) | 9.7e-10 |
|  | 201858_s_at | 640 | FP | 0.57(0.47-0.7) | 4.8e-08 |
|  | 201859_at | 640 | FP | 0.57(0.46-0.7) | 1e-07 |
|  | 201859_at | 875 | OS | 0.58(0.49-0.69) | 2.1e-10 |
|  | 201859_at | 498 | PPS | 0.56(0.44-0.7) | 3.8e-07 |
| Liver cancer | 5552 | 366 | PFS | 0.67( 0.5-0.91) | 0.0095 |
|  | 5552 | 364 | OS | 0.68(0.47-0.97) | 0.034 |
|  | 5552 | 313 | RFS | 0.65((0.46-0.92) | 0.015 |
|  | 5552 | 357 | DSS | 0.58(0.35-0.97) | 0.034 |

The survival analysis of SRGN expression in pan-cancer. NA: the data was not shown; <sup>\*</sup>Split patients by auto select best cutoff, other factors such as pathology, risk factors, patient, and follow up threshold, etc., were chosen all; <sup>#</sup>group cutoff:
median, cutoff -high/low: 50%. AML, acute myeloid leukemia ; DLBCL, diffuse large B-cell lymphoma ; LGG, brain lower grade glioma; ACC, adrenocortical carcinoma; SKCM, skin cutaneous melanoma; TGCT, testicular germ cell tumor; SARC, sarcoma; GBM, glioblastoma multiforme; OS, overall survival; PFS, progression free survival; RFS, relapse-free survival; PPS, post progression survival; FP, first progression; DSS, disease-specific survival; DMFS, distant metastasis-free survival; EFS, event free survival.

We examined the prognostic potential of SRGN mRNA in pan-cancer based on GeneChip and RNA-seq using KM-plotter. High SRGN levels were associated with poor OS, PFS, and post-progression survival (PPS) in OV (Supplementary Figure 6A). In BRCA, SRGN was a risk factor for PPS and RFS, and a protective factor for OS (Supplementary Figure 6B). In gastric cancer, high SRGN expression was associated with a favorable prognosis in terms of OS, first progression, and PPS (Supplementary Figure 6C). In lung cancer, high SRGN levels were significantly associated with a favorable or poor prognosis for OS (201859_at and 201858_s_at, respectively), and a favorable prognosis in terms of first progression (Supplementary Figure 6D).

### SRGN was a favorable prognostic factor in liver cancer according to the Kaplan–Meier Plotter Database

We investigated the relationship between SRGN expression and survival in patients with liver cancer using KM-plotter. High SRGN expression was associated with better OS, RFS, PFS, and DSS after correcting for stage, grade, AJCC_T stage, vascular invasion, sex, race, sorafenib treatment, hepatitis virus, and alcohol consumption (Supplementary Figure 6E; RNA_SeqID 5552; PFS HR = 0.67, 95% CI = 0.5–0.91, P = 0.0095, median survival 16.73 *vs.* 30.4 months; OS HR = 0.68, 95% CI = 0.47–0.97, P = 0.034, median survival 52 *vs.* 70.5 months; RFS HR = 0.65, 95% CI = 0.46–0.92, P = 0.015, median survival 21.87 *vs.* 40.97 months; and DSS HR = 0.58, 95% CI = 0.35–0.97, P = 0.034). However, the median DSS times were 104.17 *vs.* 84.4 months in the low *vs.* high expression cohorts. The prognostic potential of SRGN for survival may change over time. This was reflected by UALCAN, with SRGN indicating a poor prognosis in high-expression cohorts (P = 0.5). We further analyzed SRGN in relation to clinicopathological factors in liver cancer (Table 2). High SRGN mRNA expression was correlated with favorable OS and PFS in patients with stage 2–3+4, RFS in stages 2–3, and DSS in stages 2+3–3+4, with better OS, PFS, and DSS in AJCC_T 2 and 3, and with favorable RFS in AJCC_T 2. High SRGN expression was associated with better OS and DSS in patients with grade 2 cancer, better RFS in grades 2–3, and better PFS in grades 1–3. High SRGN expression was also associated with favorable OS, RFS, PFS, and DSS in men, Asians, and in patients with a history of alcohol consumption, and with favorable OS in women and Caucasian patients. Over-expression of SRGN in patients with hepatitis virus infection was associated with favorable prognoses in terms of RFS and PFS, and with favorable OS and DSS in patients without virus infection.

**Table 2.** Correlation of SRGN mRNA expression and prognosis in liver cancer with different clinical factors by Kaplan-Meier plotter.

| Clinical characteristics* | OS (n= 364) |  |  | RFS(n=313) |  |  |
| --- | --- | --- | --- | --- | --- | --- |
|  | N | Hazard ratio | p-value | N | Hazard ratio | p-value |
| **Male** | 246 | 0.56(0.36-0.88) | **0.01** | 208 | 0.56(0.36-0.88) | **0.0051** |
| **Female** | 118 | 0.49(0.24-0.99) | **0.044** | 105 | 0.56(0.28-1.1) | 0.085 |
| **White** | 181 | 0.58(0.36-0.93) | **0.022** | 147 | 0.75(0.48-1.18) | 0.21 |
| **Asian** | 155 | 0.51(0.28-0.94) | **0.028** | 143 | 0.55(0.33-0.92) | **0.022** |
| **Hepatitis virus(yes)** | 150 | 0.72(0.38-1.37) | 0.31 | 138 | 0.53(0.31-0.88) | **0.014** |
| **(no)** | 167 | 0.39(0.23-0.64) | **0.00014** | 142 | 0.65(0.38-1.12) | 0.12 |
| **Alcohol consumption(yes)** | 115 | 0.35(0.18-0.7) | **0.0017** | 98 | 0.52(0.28-0.97) | **0.037** |
| **(no)** | 202 | 0.65(0.38-1.1) | 0.1 | 182 | 0.63(0.38-1.02) | 0.06 |
| **Sorafenib treatment** | 29 | 0.51(0.17-1.54) | 0.23 | 22 | 0.41(0.16-1.07) | 0.06 |
| **Vascular invasion(none)** | 203 | 0.65(0.38-1.12) | 0.12 | 175 | 0.74(0.45-1.21) | 0.23 |
| **(micro)** | 90 | 2.14(0.74-6.22) | 0.15 | 81 | 0.59(0.27-1.3) | 0.18 |
| **AJCC_T(1)** | 180 | 0.76(0.42-1.39) | 0.37 | 160 | 1.27(0.72-2.24) | 0.4 |
| **(2)** | 90 | 0.44(0.21-0.95) | **0.031** | 79 | 0.46(0.21-1.01) | **0.046** |
| **(3)** | 78 | 0.46(0.23-0.9) | **0.021** | 65 | 0.56(0.27-1.15) | 0.11 |
| **Grade (1)** | 55 | 0.44(0.17-1.14) | 0.084 | 45 | 1.74(0.64-4.71) | 0.27 |
| **2** | 174 | 0.55(0.33-0.94) | **0.026** | 147 | 0.53(0.31-0.93) | **0.024** |
| **3** | 118 | 1.3(0.69-2.46) | 0.42 | 106 | 0.54(0.31-0.93) | **0.025** |
| **Stage 1** | 170 | 0.73(0.39-1.38) | 0.34 | 153 | 1.55(0.87-2.76) | 0.14 |
| **1+2** | 253 | 0.67(0.42-1.09) | 0.11 | 227 | 0.74(0.48-1.13) | 0.16 |
| **2** | 83 | 0.45(0.2-1.01) | **0.046** | 74 | 0.51(0.25-1.05) | 0.063 |
| **2+3** | 166 | 0.49(0.3-0.79) | **0.0028** | 142 | 0.52(0.33-0.81) | **0.0033** |
| **3** | 83 | 0.41(0.21-0.8) | **0.0071** | 68 | 0.54(0.29-1.01) | 0.051 |
| **3+4** | 87 | 0.45(0.24-0.85) | **0.011** | 68 | 0.54(0.29-1.01) | 0.051 |
| **Clinical characteristics** | PFS (n=366) |  |  | DSS(n=357) |  |  |
|  | N | Hazard ratio | p-value | N | Hazard ratio | p-value |
| **Male** | 246 | 0.54□0.37-0.77□ | **0.00073** | 241 | 0.53□0.29-0.97□ | **0.037** |
| **Female** | 120 | 0.56□0.31-1.01□ | 0.051 | 116 | 0.41□0.16-1.08□ | 0.062 |
| **White** | 183 | 0.73□0.49-1.07□ | 0.11 | 177 | 0.62□0.35-1.11□ | 0.1 |
| **Asian** | 155 | 0.58□0.36-0.94□ | **0.025** | 152 | 0.31□0.14-0.68□ | **0.002** |
| **Hepatitis virus(yes)** | 152 | 0.55□0.34-0.88□ | **0.011** | 149 | 1.54□0.67-3.55□ | 0.31 |
| **(no)** | 167 | 0.64□0.39-1.03□ | 0.062 | 163 | 0.37□0.2-0.69□ | **0.0013** |
| **Alcohol consumption(yes)** | 115 | 0.43□0.25-0.74□ | **0.002** | 115 | 0.29□0.13-0.66□ | **0.0017** |
| **(no)** | 204 | 0.68□0.43-1.08□ | 0.098 | 197 | 0.63□0.31-1.29□ | 0.2 |
| **Sorafenib treatment** | 30 | 0.37□0.15-0.9□ | **0.023** | 29 | 0.51□0.17-1.54□ | 0.23 |
| **Vascular invasion(none)** | 204 | 0.84□0.54-1.3□ | 0.43 | 200 | 1.46□0.7-3.04□ | 0.31 |
| **(micro)** | 91 | 0.61□0.34-1.08□ | 0.084 | 88 | 4.62(0.6-35.68) | 0.11 |
| **AJCC_T(1)** | 180 | 1.58□0.95-2.63□ | 0.073 | 177 | 3.82(0.9-16.26) | 0.051 |
| (2) | 92 | 0.45□0.25-0.8□ | **0.0049** | 89 | 0.36(0.13-1.04) | **0.049** |
| (3) | 78 | 0.46□0.24-0.88□ | **0.016** | 75 | 0.4(0.18-0.89) | **0.021** |
| **Grade (1)** | 55 | 0.4□0.18-0.9□ | **0.023** | 55 | 0.32(0.09-1.17) | 0.072 |
| **2** | 175 | 0.6□0.37-0.96□ | **0.031** | 169 | 0.48(0.24-0.95) | **0.032** |
| **3** | 119 | 0.54□0.32-0.93□ | **0.024** | 116 | 1.31(0.61-2.83) | 0.49 |
| **Stage 1** | 170 | 1.62□0.96-2.74□ | 0.071 | 167 | 3.18(0.74-13.74) | 0.1 |
| **1+2** | 254 | 0.69□0.45-1.06□ | 0.088 | 249 | 0.67(0.34-1.34) | 0.26 |
| **2** | 84 | 0.44□0.24-0.82□ | **0.0083** | 82 | 0.33(0.1-1.08) | 0.054 |
| **2+3** | 167 | 0.49□0.33-0.74□ | **0.00043** | 163 | 0.38(0.21-0.7) | **0.0013** |
| **3** | 83 | 0.45□0.24-0.82□ | **0.0074** | 81 | 0.36(0.17-0.78) | **0.007** |
| **3+4** | 88 | 0.45□0.25-0.81□ | **0.0066** | 84 | 0.37(0.18-0.79) | **0.0075** |

\* The sample numbers with some clinical characteristics were too low for meaningful analysis, such as Black or Africa American, Grade 4, macro invasion, etc. N: sample size.

### Prognostic potential of SRGN expression associated with immune cells in pan-cancer

We analyzed SRGN expression in relation to six immune infiltrates using TIMER 2.0. Correcting for six immune infiltrates and clinical factors (age, sex, stage, and tumor purity), high SRGN expression was a risk factor for OS in cholangiocarcinoma (CHOL), LUAD, and thymoma (THYM), but a favorable prognostic factor in LIHC and SKCM metastasis (Supplementary Table 7). Only SKCM metastasis was significantly different between the groups with low and high SRGN levels (log-rank test, P < 0.001; Supplementary Figure 8).

### Correlation and prognostic potential of SRGN with immune cells in LIHC

SRGN expression was significantly positively correlated with immune infiltrations of CD4+ T cells, B cells, macrophages, CD8+ T cells, neutrophils, and dendritic cells (Supplementary Figure 9). The correlations were moderate to strong (r = 0.34, 0.483, 0.579, 0.616, 0.631, and 0.714, respectively; P = 0.40e^-11^, 1.73e^-21^, 7.02e^-32^, 4.16e^-37^, 9.47e^-40^, and 2.94e^54^, respectively). Cancer type (LIHC), clinical characteristics (age, stage, tumor purity, sex, and race), six immune infiltrates, and gene symbols (*SRGN*) were analyzed using a multivariable Cox proportional hazard model. Of 305 patients enrolled, 101 died. Race (black), stage 4, macrophages, and dendritic cells were risk factors for OS, whereas stage 3, CD4+ T cells, CD8+ T cells, and SRGN were favorable factors (Supplementary Table 7). However, there were no significant survival differences for the six types of immune cells and SRGN in the low-*vs.* high-expression cohorts according to log-rank test (Supplementary Figure 8).

### Correlations between SRGN and TME cells and gene markers in LIHC

We analyzed the correlations and outcomes for each immune cell subset and other TME cell types using TIMER 2.0. SRGN expression was significantly negatively correlated with tumor purity (Rho = −0.53, P = 1.67e^-26^). Eleven proportions of cell subsets were negatively and 32 were positively correlated with SRGN expression (Table 3, Supplementary Figure 10). Immune cells included 10 CD4+ T cell subsets (CD4+ and CD4+ naïve T cells (by two algorithms CIBERSORT and XCELL), central memory, effector memory, activated memory, resting memory, and memory CD4+ T cells, CD4+ (non-regulatory), and Th1 and Th2 CD4+ T cells); two CD8+ T cell subsets (central memory and effector memory CD8+ t cells); three B cell subsets (naïve, plasma_(CIBERSORT-ABS and XCELL), and memory B cells); regulatory and follicular helper T cells; monocytes, four macrophage subsets (macrophages (EPIC and TIMER), and M0, M1, and M2 macrophages (CIBERSORT-ABS and TIDE); common lymphoid progenitors; neutrophils; three natural killer (NK) cell subsets (NK cells and resting and activated NK cells); three myeloid dendritic cell subsets (myeloid dendritic cells and activated and resting myeloid dendritic cells); and NK T cells. Naïve CD4+ T cells, plasma B cells, macrophages, and M2 macrophages were negatively or positively correlated with SRGN expression according to different algorithms: *e.g*., M2 macrophages (CIBERSORT-ABS: r = 0.696, P = 3.26e^51^; TIDE: r = −0.516, P = 0.66e^-25^). CD8+ T cells, M1 macrophages, myeloid dendritic cells, activated NK cells, and ECs showed strong positive correlations with SRGN expression levels (r = 0.552, P = 5.79e^-29^; r = 0.517, P = 5.84e^-25^; r = 0.627, P = 4.97e^-39^; r = 0.524, P = 1.11e^-25^; r = 0.522, P = 1.67e^-25^, respectively). The remaining six cell types, including hematopoietic stem cells (HSCs) and myeloid-derived suppressor cells (MDSCs), showed no correlation with SRGN expression.

**Table 3.** The correlation between SRGN and infiltrating cells in LIHC by TIMER2.0.

| TME Cells | Rho | p-value |
| --- | --- | --- |
| T cell CD4+ | 0.368 | 1.76e-12 |
| T cell CD4+ naïve_CIBERSORT) | -0.152 | 4.69e-03 |
| T cell CD4+ naïve_XCELL) | 0.171 | 1.44e-03 |
| T cell CD4+ central memory | -0.262 | 7.79e-07 |
| T cell CD4+ effector memory | 0.107 | 4.8e-02 |
| T cell CD4+ memory activated | 0.12 | 2.64e-02 |
| T cell CD4+ memory resting | 0.412 | 1.55e-15 |
| T cell CD4+ memory | 0.305 | 7.2e-09 |
| T cell CD4+(non-regulatory) | 0.175 | 1.1e-03 |
| T cell CD4+ Th1 | -0.234 | 1.09e05 |
| T cell CD4+ Th2 | 0.206 | 1.19e-04 |
| **T cell CD8+** | **0.552** | **5.79e-29** |
| T cell CD8+ central memory | 0.336 | 1.49e-10 |
| T cell CD8+ effector memory | 0.119 | 2.65e-02 |
| B cell | 0.308 | 5.19e-09 |
| B cell naïve | 0.145 | 7.06e-03 |
| B cell plasma_CIBERSORT-ABS | 0.121 | 2.4e-02 |
| B cell plasma_XCELL | -0.11 | 4.12e-02 |
| B cell memory | 0.108 | 4.49e-02 |
| T cell regulatory | 0.399 | 1.34e-14 |
| T cell follicular helper | 0.252 | 2.22e-06 |
| Monocyte | 0.481 | 2.21e-21 |
| Macrophage/monocyte | 0.481 | 2.21e-21 |
| Macrophage_EPIC | -0.382 | 1.9e-13 |
| Macrophage_TIMER | 0.481 | 2.31e-21 |
| Macrophage M0 | -0.123 | 2.23e-02 |
| **Macrophage M1** | **0.517** | **5.84e-25** |
| **Macrophage M2_CIBERSORT-ABS** | **0.696** | **3.26e-51** |
| **Macrophage M2_TIDE** | **-0.516** | **6.66e-25** |
| **Myloid dendritic cell** | **0.627** | **4.97e-39** |
| Myloid dendritic cell activated | 0.475 | 7.62e-21 |
| Myloid dendritic cell resting | 0.142 | 8.13e-03 |
| Common lymphoid progenitor | 0.211 | 8.13e-05 |
| Neutrophil | 0.426 | 1.11e-16 |
| NK cell | -0.115 | 3.26e-02 |
| NK cell resting | -0.164 | 2.18e-03 |
| **NK cell activated** | **0.524** | **1.11e-25** |
| T cell NK | -0.147 | 6.11e-03 |
| Mast cell | 0.109 | 4.27e-02 |
| Mast cell resting | 0.192 | 3.23e-04 |
| Mast cell activated | -0.166 | 2.03e03 |
| **Endothelial cell** | **0.522** | **1.67e-25** |
| Cancer associated fibroblast | 0.456 | 4.09e-19 |

We further analyzed the correlations between SRGN expression and biomarkers of CD8+ T cells, M1 and M2 macrophages, tumor-associated macrophages, and monocytes in pan-cancer, especially LIHC (Supplementary Figures 11 and 12). The gene markers have been reported previously [18]. All the markers, as well as CTLA4, PDCD1, CD274, CD80, CD44, vascular endothelial growth factor C, and transforming growth factor-β2, were positively correlated with SRGN in LIHC. UALCAN analysis revealed SRGN-associated genes in LIHC. The monocyte marker CD86 was the top positive correlated gene (Figure 3A and B). According to HPA analysis, markers of vascular ECs, neutrophils, and Kupffer cells showed strong correlations with SRGN (r > 0.5, Figure 3C). STRING analysis showed SRGN–protein interactions (Figure 3D). The top three proteins were granzyme B, perforin-1, and CD44 antigen.

**Figure 3.**
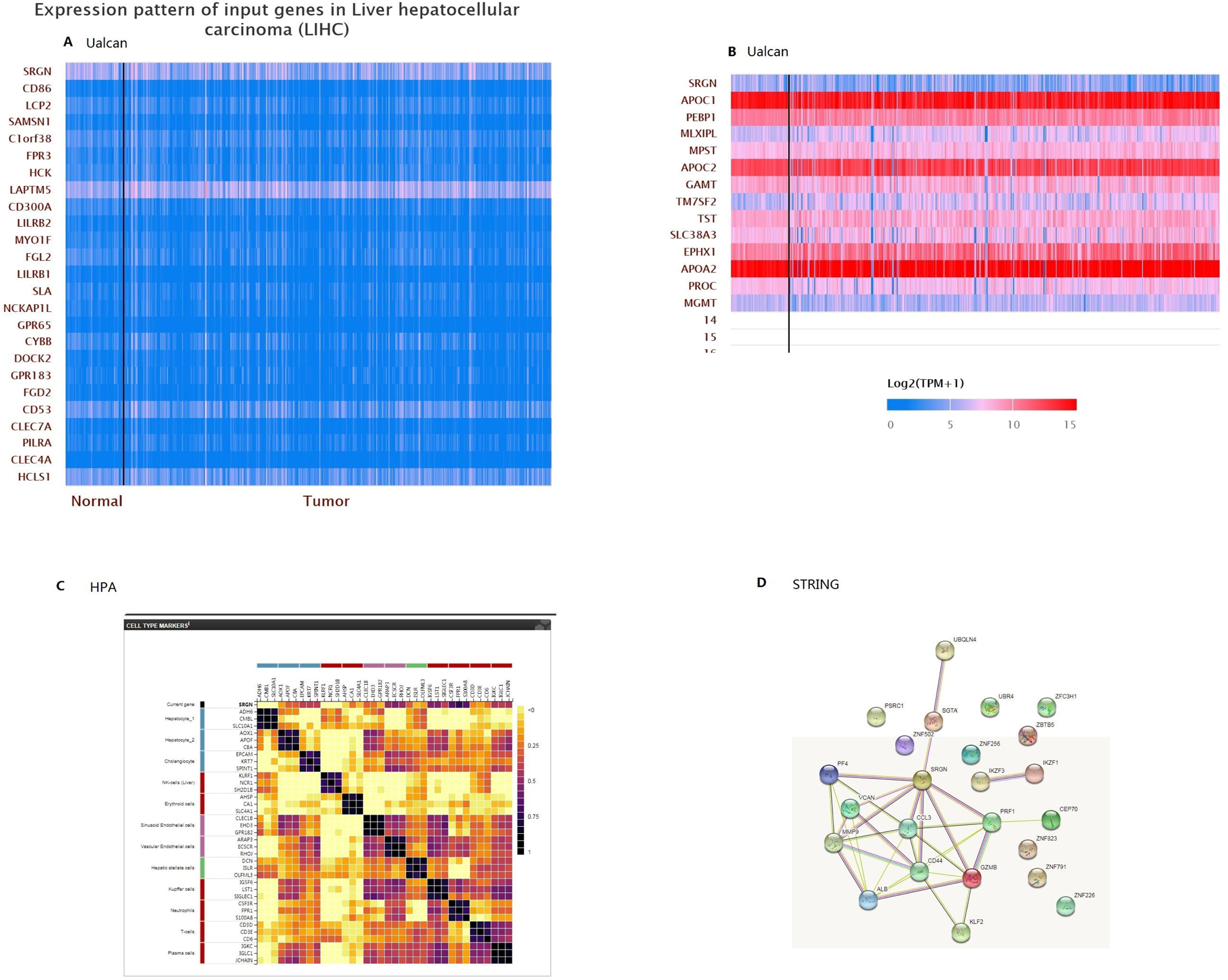
Genes positively (A) and negatively (B) correlated with SRGN in LIHC by UALCAN analysis (data from TCGA). (C) Correlations between SRGN and liver cell biomarkers via HPA. (D) Predicted functional proteins with SRGN (*Homo sapiens*) by STRING analysis (https://cn.string-db.org/cgi/network?taskId=bHNlp6Ws7IcB&session_Id=bSY6tL5m6eXy).

### Prognostic potential of SRGN with TME cells in LIHC

We analyzed TME infiltrates (21 cell types), clinical characteristics (age, stage, purity, sex, and race), and SRGN in relation to OS in patients with LIHC using the Cox proportional hazards model. The cell types and subsets are shown in Table 4 and Figure 4, with SRGN as a significant prognostic factor in the Cox model. According to log-rank test, high infiltration of MDSCs, M1 macrophages, or monocyte_MCPCOUNTER was associated with unfavorable OS in the cohort with high SRGN expression (HR = 2.78, P = 0.00668; HR = 3.09, P = 0.00633; HR = 2.75, P = 0.025, respectively), whereas high infiltration of CD8+ T cells, resting memory CD4+ T cells, ECs, or HSCs was associated with a favorable prognosis for OS (HR = 0.53, P = 0.048; HR = 0.501, P = 0.0383; HR = 0.364, P = 0.00638; HR = 0.379, P = 0.00868, respectively). In the cohort with low SRGN expression, high infiltration of macrophage_TIMER, macrophage M0_CIBERSORT, and macrophage M2_CIBERSORT-ABS were unfavorable factors for OS (HR = 1.82, P = 0.0432; HR = 1.86, P = 0.0356; HR = 1.99, P = 0.0366, respectively).The three cellsubsets, together with CD8+ T cell, showed time-dependent survival curves. The log-rank test of SRGN expression with the other six cell types (central memory and Th2 CD4+ T cells, Tregs, monocytes (XCELL), myeloid dendritic cells, and resting myeloid dendritic cells) showed no significant difference in LIHC.

**Figure 4.**
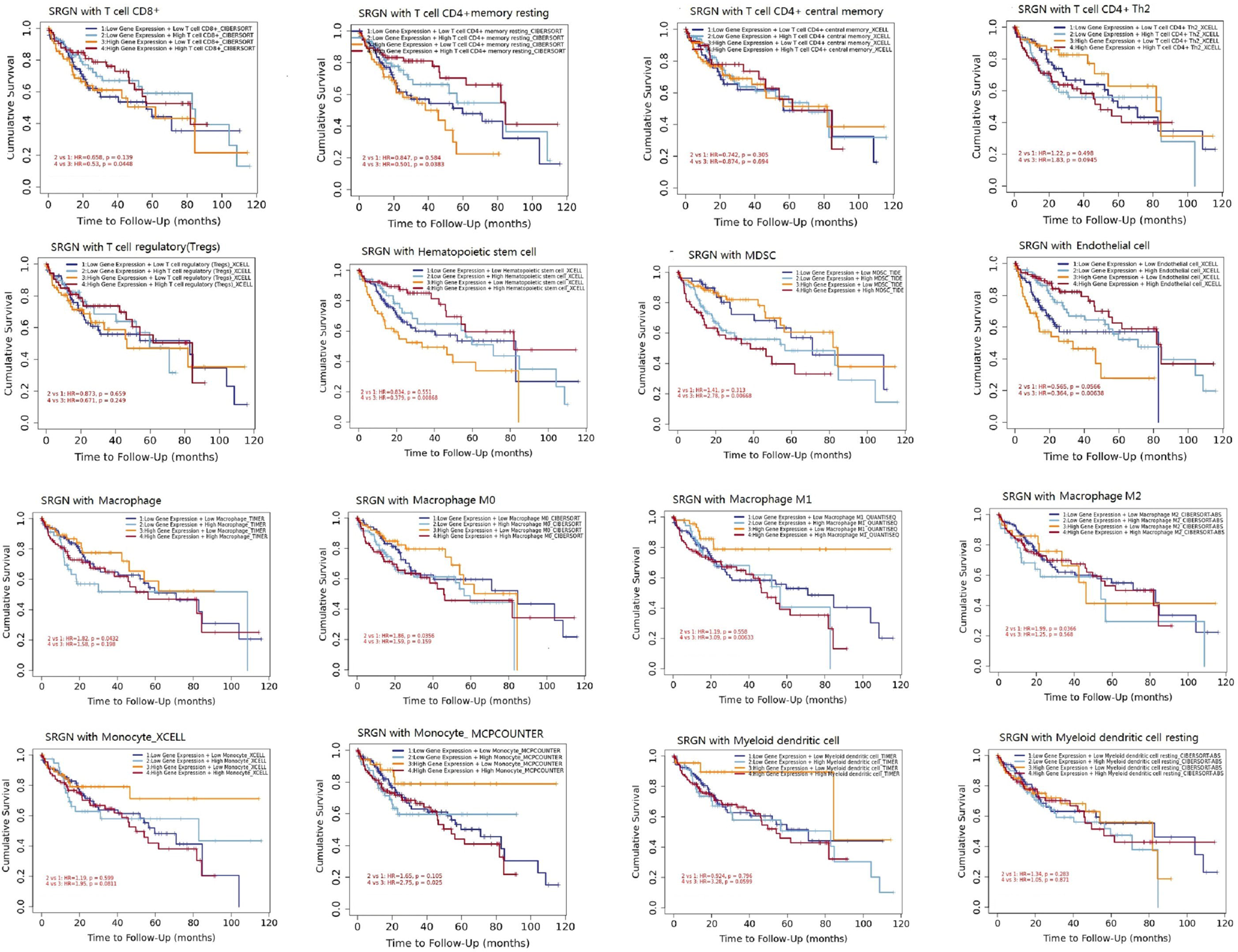
Comparisons of Kaplan–Meier survival curves between cohorts with high and low levels of SRGN expression associated with high and low levels of infiltrating cells in LIHC via TIMER2.0.

**Table 4.** The survival analysis of SRGN expression associated with infiltrating cells in LIHC by TIMER2.0.

| Cells in TME | Low SRGN |  | High SRGN |  |
| --- | --- | --- | --- | --- |
|  | expression | p-value | expression | p-value |
|  | Hazard ratio |  | Hazard ratio |  |
| T cell CD8+ | 0.658 | 0.139 | 0.53 | **0.0448** |
| T cell CD4+ memory resting | 0.847 | 0.584 | 0.501 | **0.0383** |
| Hematopoietic stem cell | 0.834 | 0.551 | 0.379 | **0.00868** |
| MDSC | 1.41 | 0.313 | 2.78 | **0.00668** |
| Endothelial cell | 0.565 | 0.0566 | 0.364 | **0.00638** |
| Monocyte(MCPcounter) | 1.65 | 0.105 | 2.75 | **0.025** |
| Macrophage | 1.82 | **0.0432** | 1.58 | 0.198 |
| Macrophage M1 | 1.19 | 0.558 | 3.09 | **0.00633** |
| **Macrophage 0** | 1.86 | **0.0356** | 1.59 | 0.159 |
| **Macrophage 2** | 1.99 | **0.0366** | 1.25 | 0.568 |
| T cell CD4+ central memory | 0.742 | 0.305 | 0.874 | 0.694 |
| T cell CD4+ Th2 | 1.22 | 0.498 | 1.83 | 0.0945 |
| Tregs | 0.873 | 0.659 | 0.671 | 0.249 |
| Monocyte(XCELL) | 1.19 | 0.599 | 1.95 | 0.0811 |
| Myloid dendritic cell | 0.924 | 0.796 | 3.28 | 0.0599 |
| Myloid dendritic cell resting | 1.34 | 0.283 | 1.05 | 0.871 |

TME: tumor microenvironment

### *In vitro* and *in vivo* pro-tumorigenicity of SRGN

SRGN promoted the proliferation of SRGN-over-expressing HepG2SG cells compared with HepG2SG-NC cells (transfected with blank control vector) and HepG2, respectively (P < 0.05, P < 0.01; Figure 5A-C). HepG2SG cells showed weak sorafenib resistance (resistance index 1–2; Figure 5D). HepG2SG cells also showed increased cell invasion compared with HepG2-NC cells by Transwell assay (P ≤ 0.001, Figure 5E). The number of tubular structures was higher in human umbilical vein ECs (HUVECs) cultured with HepG2SG supernatant compared with HepG2SG-NC supernatant (P ≤ 0.001; Figure 5F). SRGN promoted vascular mimicry tube formation in HepG2SG cells compared with HepG2SG-NC cells (P < 0.01; Figure 5G). Subcutaneous xenograft tumors derived from HepG2SG cells were heavier than those from HepG2SG-NC cells (P < 0.05; Figure 5H). Hematoxylin and eosin staining showed that cells were distributed in clumps or strands in subcutaneous tumor tissues (Figure 5I). SRGN and CD206 were up-regulated while CD80 was down-regulated in subcutaneous tumor tissues in HepG2SG xenograft mice compared with HepG2 SG-NC mice (P < 0.05, P < 0.01; Figure 5J).

**Figure 5.**
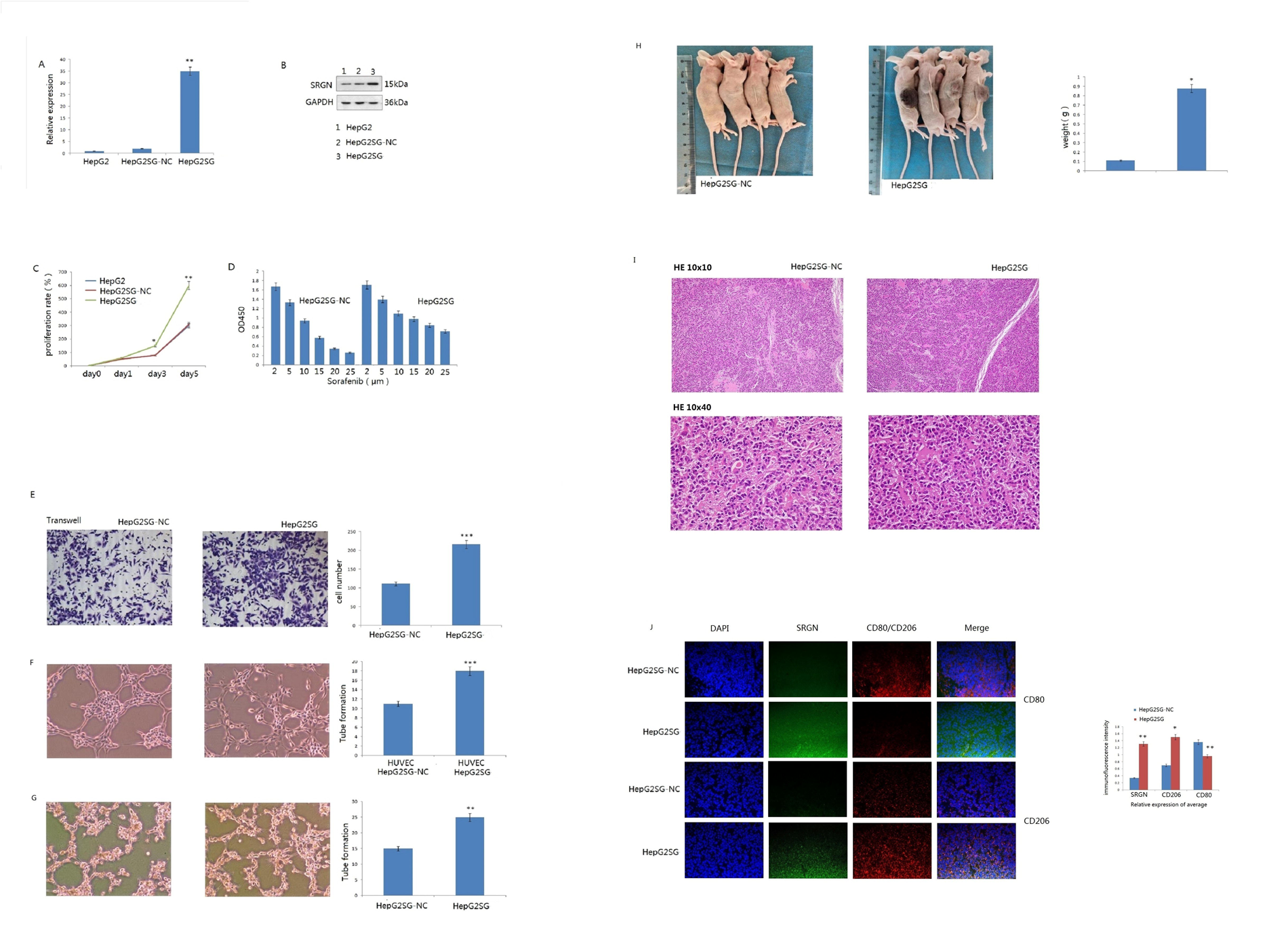
*In vitro* and *in vivo* experiments of pro-tumorigenicity of SRGN (A) Relative expression of SRGN in HepG2SG (over-expressing SRGN) and HepG2SG-NC (transfected with blank control vector) examined by quantitative reverse transcription-polymerase chain reaction. (B) Expression of SRGN protein in HepG2SG and HepG2SG-NC cells examined by western blotting. (C) Proliferation rates (%) of HepG2SG and HepG2SG-NC cells by CCK8 assay (*P < 0.05, **P < 0.01). (D) Proliferation rates (%) of HepG2SG and HepG2SG-NC cells treated with sorafenib, 2> resistance index>1. (E) Invasion of HepG2SG and HepG2SG-NC cells examined by Transwell assay (***P < 0.001). (F) Angiogenesis of HUVECs cultured with supernatant from HepG2SG and HepG2SG-NC cells, respectively (***P ≤ 0.001). (G) Vasculogenic mimicry of HepG2SG and HepG2SG-NC cells (**P < 0.01). (H) Weight of HepG2SG and HepG2SG-NC subcutaneous xenograft tumors in nude mice (*P < 0.05). (I) Hematoxylin and eosin and (J) immunofluorescence staining of subcutaneous tumor tissues. Both SRGN and CD206 were up-regulated while CD80 was down-regulated in HepG2SG mice compared with HepG2SG-NC mice (*P < 0.05, **P < 0.01). Light blue: DAPI; green: fluorescein isothiocyanate-stained SRGN; red: CY3-stained CD80 or CD206.

## Discussion

SRGN is a hematopoietic intracellular proteoglycan expressed in secretory granules and vesicles, and secreted into the extracellular medium [19]. The core SRGN protein consists of eight serine/glycine repeats attached by variable glycosaminoglycan bonds, depending on the cell type and status [20, 21]. SRGN exerts pro-tumorigenic effects in multiple ways [22], and several malignant cell types depend on SRGN to promote aggregation [23]. However, its role remains controversial.

TIMER, GEPIA, and KM-Plotter database analyses showed that SRGN levels were higher in AML than in other cancers, and SRGN expression was a prognostic risk factor for survival in four AML and two DLBCL groups according to PrognoScan analysis. The prognostic potential of SRGN RNA expression was consistent with that for SRGN protein, as determined by immunohistochemistry (IHC) and enzyme-linked immunosorbent assay [24]. The pro-tumorigenic effect of SRGN was recently confirmed in AML [25]. However, no consistent role has been demonstrated in solid tumors.

IHC analyses in lung, breast, prostate, and colon cancer cells showed that SRGN expression was up-regulated in advanced tumors and in the activated TME [26]. In the present study, SRGN RNA levels were lower in BRCA and lung cancer compared with normal adjacent tissues according to TIMER, KM-plotter, and GEPIA analyses, as also reported by Oncomine [27]. SRGN protein levels were decreased in the above two cancers compared with normal tissues according to CPTAC. PrognoScan analysis demonstrated that high levels of SRGN mRNA were associated with a favorable prognosis in small samples of patients with OV, lung cancer, and BRCA, but SRGN RNA expression was also a prognostic risk factor in a large sample size in three cancers analyzed via KM-plotter, and experimental and clinical studies have identified pro-tumorigenic effects of SRGN in lung cancer and BRCA [27–33]. SRGN was correlated with drug resistance and stemness in BRCA, promoting invasion and metastasis and inducing BRCA cell aggregation [27–31]. High levels of SRGN predicted a poor prognosis in LUAD, promoting NSCLC cell migration [32, 33]. The current results for SKCM were consistent with a previous report that identified SRGN as a favorable factor in SKCM and SKCM-metastasis tissues via GEPIA and TIMER 2.0 [27], indicating that the result was reproducible by bioinformatics analysis. However, the prognostic potential and tumorigenicity of SRGN in sarcoma, glioblastoma multiforme, testicular germ cell tumors, gastric cancer, CHOL, and THYM have rarely been studied.

The controversial role of SRGN is more evident in LIHC. SRGN mRNA levels were lower in LIHC than in adjacent normal tissues via TIMER, but were higher in liver cancer *vs.* normal tissues via Oncomine [27], showed no difference via GEPIA, and were inconsistent between GeneChip and RNA-seq data via KM-plotter. IHC staining showed that SRGN levels were higher in 56.7% of HCC tissues *vs.* 3.1% of adjacent non-tumor tissues [9]. We previously showed that levels of hematopoietic SRGN examined by flow cytometry were higher in patients with chronic hepatitis B, liver cirrhosis, and LIHC than in healthy adults, while levels of SRGN mRNA were lower in HepG2.215 compared with HepG2 cells [11, 12]. HepG2 is a hepatoblastoma-derived cell line. However, it expresses the lowest SRGN level among 24 liver cancer cell lines (https://www.proteinatlas.org/ENSG00000122862-SRGN/cell+line), and HepG2 cells show poor *in vivo* tumorigenicity. In the present study, HepG2 cells over-expressing SRGN had significant proliferation *in vitro* and tumorigenicity *in vivo*, which identified pro-tumorigenicity of SRGN in liver cancer. SRGN expression may be affected by inflammation and malignant transformation induced by HBV, thus increasing the complexity of survival analyses.

SRGN mRNA expression levels showed no prognostic potential in LIHC *via* PrognoScan, GEPIA, or TIMER 2.0 (analyzed without TME cells). According to the KM-plotter database, SRGN mRNA was a favorable prognostic factor in LIHC after adjusting for stage, grade, AJCC_T, sex, race, alcohol consumption, hepatitis virus infection, sorafenib treatment, and vascular invasion; however, SRGN protein expression was an unfavorable factor for OS and time to recurrence [9, 12]. The prognostic potential of the SRGN transcriptome and protein were thus contradictory, as in lung cancer and BRCA. As a prominent hematopoietic proteoglycan, SRGN should be analyzed together with TME cells in a multivariate Cox regression model.

The liver is the major hematopoietic organ during the fetal period. Hepatic hematopoiesis can generate lymphoid, myeloid, and erythroid lineages, and is involved in tumor surveillance and rejection [34, 35]. SRGN maintains homeostasis of the body’s immune cell populations, controlling the magnitude and durability of the immune response [36]. In the present study, six prominent immune cells showed strong correlations with SRGN expression, and SRGN was a favorable prognostic factor for OS associated with macrophages, dendritic cells, and CD4^+^ and CD8^+^ T cells. However, there are abundant immune cell subsets, and further analyses are required.

SRGN has been studied in neutrophils, cytotoxic T lymphocytes, macrophages, CD8+ T cells, platelets, and NK cells [8, 37–41]. The negative correlation between SRGN and tumor purity suggested that SRGN may be prominently expressed in TME-infiltrating cells, and the correlations between SRGN and TME cells was reliable. The results showed that both neutrophils and monocytes were more prevalent in normal liver tissues than in LIHC and normal tissues adjacent to LIHC. M2 macrophages acquire pro-tumorigenic properties by responding to CCL2 and interleukin-10, triggering the release of cytokines, chemokines, and growth factors to promote malignancy [42]. M2 macrophages with immunosuppressive properties were the most abundant of the five types of immune cells in healthy liver tissues, as in LIHC and adjacent normal tissues, which may reflect the fact that the liver demonstrates natural immune tolerance [4]. M1 macrophages and CD8+ T cells antagonize pro-tumorigenesis and both were second to M2 macrophages in LIHC and adjacent normal tissues. These results suggested that the differentiation and activation of monocytes/macrophages may vary in physiological and pathological liver tissues. SRGN can be induced in macrophages [37]. The glycosaminoglycan of SRGN varies according to the differentiation of monocytes to macrophages or the activation of macrophages [20]. SRGN level were higher in M2 macrophages than in neutrophils in LIHC, adjacent normal, and liver tissues by GEPIA, but SRGN in neutrophils was dominant in the liver by HPA. Spearman’s correlation analysis indicated significant correlations between SRGN and 43 TME cells in LIHC. Together with the above previous studies [8, 37–41], these results suggest that SRGN may have an extensive impact on the immune system. CTLA4, PDCD1, CD274, and granzyme B, as markers of T cell exhaustion [19], were correlated with SRGN. The *in vivo* results indicated that CD206 was increased while CD80 was down-regulated in HepG2SG mice, in contrast to the positive correlation between SRGN and CD80 *via* TIMER2.0. SRGN might promote monocyte/macrophage differentiation or migration to LIHC, as reported in myeloma [43]. The TME was immunosuppressive in HepG2SG mice, but further studies are needed to verify these bioinformatics results.

Survival analysis showed that macrophages (TIMER), M0, and pro-tumorigenic M2 macrophages were associated with low SRGN levels, while monocytes (MCPCOUNTER) and “antitumorigenic” M1 macrophages with high SRGN levels were unfavorable prognostic factors. The prior cells associated with low SRGN expression might maintain the immunosuppressive status, while the latter with high SRGN expression might promote excessive proinflammatory status into immune exhaustion or tolerance status, promoting the conversion of “antitumorigenic” M1 macrophages to pro-tumorigenic ones [44]. The prognostic potential of SRGN was closely related to monocyte / macrophage subsets. The positive correlations between SRGN and CTLA4, PD-1, and PD-L1 also suggested that SRGN was related to immune escape, and that immune therapy, e.g., PD-1 inhibitors, might have an effect in SRGN-positive LIHC. CD4+ T cell subsets had extensive, weak-to-moderate correlations with SRGN (seven positive and three negative); however, only high resting memory CD4+ and CD8+ T cells in the cohort with high SRGN expression showed protective effects against LIHC by log-rank test. It is possible that high levels of SRGN may play a role in the cytotoxicity of T cells in LIHC, as reported in virus infection [39, 41]. MDSCs have immunosuppressive activity against T-cell responses, and high SRGN expression and MDSCs indicated an unfavorable prognosis. Further studies are warranted to investigate the interaction between SRGN and these immune cells.

Vascular invasion had no effect on the prognostic potential of SRGN mRNA according to KM-plotter. However, high SRGN has been reported to up-regulate the proliferation of HUVECs [45], and SRGN protein expression was positively related to vascular invasion in HCC patients [9]. The present study showed that ECs and vascular endothelial growth factor C were strongly correlated with SRGN, and SRGN induced the formation of tubular structures by both HUVECs and HepG2 cells. High levels of circulating HSCs and endothelial progenitor cells were associated with worse OS in HCC patients [46]; however, high SRGN RNA levels with high HSCs or ECs were associated with favorable OS by TIMER 2.0 analysis. Both high levels of SRGN and ECs might promote normalization of tumor vasculature; however, further studies are required to investigate this controversy.

The prognostic capability of SRGN may vary according to the covariates in Cox regression. Regardless of the algorithms, SRGN mRNA and protein expression patterns, in addition to clinicopathological factors, observation endpoints, different populations, and sample sizes, may result in contradictory survival results for pan-cancers, except for blood cancers. One of two SRGN transcript variants is protein-coding (GEPIA, provided by RefSeq, Jul 2010). The present study constructed an SRGN-over-expressing vector (based on the same sequence of ID5552 analyzed by KM-plotter) and identified pro-tumorigenicity and metastasis in PLC, as other cancers [27–33]. Menyhárt et al. reviewed 318 genes related to HCC survival and showed that none had equivalent prognostic value to tumor stage. They also noted that survival results according to protein expression were inconsistent with those for the transcriptome [47]. SRGN may have an important effect on TME cells according to the multivariate Cox regression model.

In addition to the above limitations, the sequencing data and microarray analysis of tumor tissues might have systematic bias. Further studies combined with spatiotemporal single-cell RNA sequencing are needed to clarify the results. Second, variable glycosaminoglycan bonds might interfere with the function of SRGN. Although SRGN showed contradictory expression and prognostic potential in bioinformatics analysis across different databases, both *in vivo* and *in vitro* experiments were performed to identify its pro-tumorigenicity in many reports [9, 12, 27–33]. Future studies could clarify more mechanisms of SRGN in PLC.

Although SRGN was a limited prognostic factor in PLC, the results comprehensively revealed relationships between SRGN and monocyte/macrophage subsets, ECs, HSCs, MDSCs, CD8^+^ T cells, and resting memory CD4^+^ T cells, which may contribute to the development of new immunotherapy strategies.

## Acknowledgements

We thank International Science Editing (http://www.International_scienceediting.com) for editing a draft of this manuscript.

## Availability of data and materials

Please contact the corresponding author Yongwei Li upon reasonable requests.

## Ethics approval and consent to participate

Not Applicable for patient’s ethics approval.

