## Supplementary material for "Important Role of Hematopoietic Proteoglycan Serglycin in Liver Hepatocellular Carcinoma Associated with Tumor Microenvironment": supplemmental: SRGN生物信息学投稿last.docx

Zengcheng Zou^1^, Heping Xie^1^, Wenhai Guo^1^, Yue Li^1^, Jiongshan Zhang^1*^, Yongwei Li^1*^

^1^ Department of Traditional Chinese Medicine, The Third Affiliated Hospital of Sun Yat-sen University, Guangzhou, Guangdong, China

Zengcheng Zou, Heping Xie and Wenhai Guo are co-authors

Funding statement: Supported by Science and Technology Planning Project of Guangdong Province, China, no. 2017A030313738, 2022A1515011689, 2023A1515011937

the construction project of inheritance studio of national famous and old traditional Chinese Medicine experts, no.140000020132

**Abstract**

**Background** Serglycin (SRGN) is a prominent hematopoietic proteoglycan that regulates tumorigenesis; however, its role in tumor immunity is unclear.

**Materials and methods** We investigated the expression and prognostic potential of SRGN in liver hepatocellular cancer (LIHC) in the context of pan-cancer (for showing the similarity and heterogeneity) using the PrognoScan, GEPIA, Kaplan–Meier Plotter, and TIMER bioinformatics databases. HepG2 cells were transfected with an SRGN over-expression vector, and their proliferation, invasion, sorafenib resistance, and vasculature were examined in vitro. We also created a subcutaneous xenograft tumor model in nude mice.

**Results** SRGN expression was prominent in M2 macrophages in LIHC. The Kaplan–Meier Plotter indicated that SRGN RNA was a favorable prognostic factor after correcting for clinical factors. TIMER 2.0 showed that the immune infiltrates of CD8+ T cells, M1 and M2 macrophages, and endothelial cells were strongly correlated with SRGN RNA expression (r=0.552, P=5.79e^-29^; r=0.517, P=5.84e^-25^; r=0.696, P=3.26e^51^; and r=0.522, P=1.67e^-25^, respectively), and had prognostic potential in LIHC in patients with low or high levels of SRGN, in addition to resting memory CD4+ T cells, hematopoietic stem cells (HSCs), and myeloid-derived suppressor cells (MDSCs). SRGN promoted the proliferation of HepG2 cells in vitro and in vivo, and was associated with weak sorafenib resistance, invasion, and vasculature. CD206 and CD80 were up-regulated and down-regulated, respectively, in subcutaneous tumor tissues.

**Conclusions** The results comprehensively revealed relationships between SRGN and monocyte/macrophage subsets, endothelial cells, HSCs, MDSCs, CD8+ T cells, and resting memory CD4+ T cells. These may constitute an important tumor microenvironment because SRGN promoted tumorigenesis in liver cancer cell line.

**Lentiviral transduction**

SRGN sequences were designed following the gene (Homo sapiens (human) Gene, variant 3, NM, NCBI132 Reference 105 Sequence 3.2, mRNA, NCBI132 ID: 5552). SRGN XhoI F: 5'ccgctcgaggccacc ATGATGCAGAAGCTACTCAAATGCAGTC3',BamHI R: 5'cgcggatcc TTATAACATAAAATCCTCTTCTAATCCATG 3'. RNA was extracted by Trizol, amplificated by PCR. SRGN and pLVX- IRES-Neo vector 15μL each was digested with XhoI/BamHI, recovered and connected. The ligation product was transformed to 50 µL of DH5α competent cells, identified positive clones to be cloned into pLVX-IRES-Neo vector. The shuttle plasmid containing the target sequence and the packaging plasmid pGag/Pol, pRev, pVSV-G were constructed and prepared by Guangzhou Yeshan Biotechnology Co., Ltd., and 293T cell was co-transfected with the transfection reagent LipofectamineTM 2000. After 72 hours of culture, the lentiviral particles were collected and infected HepG2 cells with 400μg/mL G418 to screen for one month. HepG2 cells stably overexpressing SRGN or a blank pLVX- IRES-Neo vector were established, and named HepG2SG and HepG2SG-NC, respectively.

**Real-time quantitative polymerase chain reaction (qPCR)**

The primers were as follows: SRGN 150 bp, F: CTGCAAAC TGCCTTGAAGAA, R: GTGGGAAGATACGATTCAAGTC; β-actin 275 bp, F: TGGATCAGCAAGCAGGAGT A, R: TCGGCCACATTGT GAACTTT. qPCR assays were performed in HepG2SG-NC group and HepG2SG. Total RNA was extracted with Trizol solution (Invitrogen Corporation, USA) following the manufacturer’s instructions. cDNA was synthesized with a first-strand cDNA synthesis kit (Takara Inc., Dalian, P. R. China). SYBR Green qPCR SuperMix (Invitrogen Corporation, USA) was used for qPCR (ABI, PRISM® 7500 Sequence Detection System). The reaction conditions were 50°C 2 min, 95°C 2 min, 95°C 15 s, 60°C 32 s, 40 cycles; melting curve analysis was at 60°C–95°C. Each sample was assayed three times. Relative mRNA expression was normalized to the corresponding β-actin expression and analyzed by the 2−△△Ct method.

**Immunoprotein electrophoresis**

Cell lysates were prepared in RIPA buffer (Thermo Scientific™, USA) and the protein concentration was determined by the bicinchoninic acid assay. Proteins were separated by 12% SDS-PAGE, and transferred to 0.45 μm polyvinylidene difluoride (PVDF) membranes (Immobilon-P Transfer Membrane, Millipore, IPVH00010). After blocking with 5% nonfat dried milk in TBST buffer, membranes were incubated overnight at 4°C with Rabbit Anti-SRGN antibody (BIOSS bs-6789R, ratio 1:1000), and GAPDH antibody as control (Kangcheng Bio, Shanghai, China, Catalog No.KC-5G5, ratio 1:10,000). After incubation with peroxidase-labeled rabbit anti-rat IgG (Bode Biotech Co., Ltd., Wuhan, China, Cat. no BA1058) secondary antibody at 1:2000 and 37°C for 1 h, the bands were read with a Pro-light HRP Chemiluminescence Kit (TIANGEN, Beijing, China), and Image J software (Gel Image Analyzer, Tianneng Technology Co., Ltd, Shanghai, China, Tanon 1220).

**CCK8 assay of cell viability**

For the cell viability assay, the cell cultures were allocated to a HepG2SG-NC group and HepG2SG groups, and treated with 2, 5, 10, 15, 20, 25μm Sorafenib, respectively. Cell viability was evaluated by an CCK8 assay (CellTiter 96 AQueous One Solution Cell Proliferation Assay, Promega, Madison,USA, Cat. No. G3582). The cell density was 1×104 cells / 100μL per well into a 96-well plate. Adherent cells need to be adhered to the wall before collecting cells at each time point for detection. Collected cells at each time point (0, 1, 3, 5 days) and added CCK-8 solution (Biyuntian, Cat. No. C0037), the ratio was 1:10. That was, 100μL culture medium wasadded to10μL detection liquid. The results were read at an absorbance of 490 nm using multiscan MK3 plate reader (Thermo Fisher Scientific Ltd, Waltham, MA, USA). The results were obtained in three independent assays with five replicates each. The effect on cell viability at each assay time was reported as inhibition rate = (1 − mean ODHepG2SG ÷ mean ODHepG2) × 100%.

**Transwell invasion assay**

Invasiveness was assayed in HepG2SG-NC and HepG2SG cultures. Matrigel (BD Biosciences, Franklin Lakes, NJ, USA) was dissolved overnight at 4°C, diluted 1:3 in prechilled serum-free medium, and 40 μL was added to prechilled Transwell chambers (BD Biosciences, USA) and incubated at 37°C for 2 h. The next day, 1 × 105 cells in 100 μL serum-free DMEM medium were added to the upper chamber and 600 μL DMEM containing 10% FBS was added to the lower chamber. After 48 h incubation, the cells adhering to the lower surface of the upper chamber were fixed in 4% paraformaldehyde and stained with crystal violet. Absorbance was measured at 570 nm with a microplate reader (Thermo Fisher multiscan MK3).

**HE staining**

Dehydration of the paraffin sections, placed the sections into xylene I 20min→xylene II 20min→anhydrous ethanol I 10min→anhydrous ethanol II 10min→95% alcohol 5min→90% alcohol 5min→80% alcohol 5min→70% alcohol 5min→washed in distilled water. Hematoxylin staining of nuclei: Sections were stained with Harris hematoxylin for 3-8 min, washed, differentiated with 1% hydrochloric acid alcohol for several seconds, rinsed with tap water, returned to blue with 0.6% ammonia water, and rinsed with running water. Eosin-stained cytoplasm: Sections were stained in eosin staining solution for 1-3min. Dehydration and sealing: put the sections in 95% alcohol I for 5 min →95% alcohol II for 5 min→anhydrous ethanol I for 5 min→anhydrous ethanol II for 5 min→xylene I for 5 min→xylene II for 5 min. then sealed with neutral gum.

**Immunofluorescence**

Dehydration of the paraffin sections, placed the sections in xylene I for 15 min, xylene II for 15 min, anhydrous ethanol I for 5 min, anhydrous ethanol II for 5 min, 85% alcohol for 5 min, 75% alcohol for 5 min, and distilled water for washing. The tissue sections were placed in a retrieval box filled with citrate antigen retrieval buffer for antigen retrieval. After natural cooling, the slides were placed in PBS and washed three times. A histochemical pen was used to draw circles around the tissue, added autofluorescence quencher to the circle for 5 minutes, and rinsed with running water for 10 minutes, added BSA in the circle and incubated for 30min, dropped the primary antibody prepared with PBS in a certain proportion on the slice, and incubated the slice at 4°C overnight in a humid box, added the secondary antibody to the primary antibody in the circle to cover the tissue, and incubated for 50 min at room temperature in the dark. The slides were placed in PBS and washed three times. DAPI staining solution was added to the circle and incubated for 10 min at room temperature in the dark, the slides in PBS washed for 3 times. After drying, the sections were mounted with anti-fluorescence quenching mounting medium. The sections were observed under a fluorescence microscope and images were collected.

**Angiogenesis experiment and vasculargenic mimicry**

The Matrigel (BD Company, 356234, USA) was added 150-200uL to each well of 48-well plates; put it in a 37°℃ incubator for more than 2 hours to solidify the Matrigel. Human umbilical vein endothelial cells (HUVECs) incubated with serum-free medium of respective HepG2SG and HepG2SG-NC for 12 h, then added 2×104 cells /200uL medium to each well, observed the vessels after 4-6h, randomly selected 5 fields of the images, analyzed the results with Image J software, and counted the number of lumens. For vasculargenic mimicry, the procedures as above, while the HepG2SG-NC and HepG2SG cells were used, and observed the vessels after 3-5 days.

**Prognostic potential of SRGN in pan-cancer according to PrognoScan, GEPIA, Kaplan–Meier Plotter Databases, and TIMER**

We analyzed the relationship between SRGN expression and prognosis in pan-cancer (Supplementary Figures 3–6, Table 1). PrognoScan analysis revealed that SRGN expression had a significant impact on prognosis in five cancer types: AML, diffuse large B-cell lymphoma (DLBCL), BRCA, lung cancer, and ovarian cancer (OV). High SRGN expression was associated with a poor prognosis in AML and DLBCL, but showed favorable prognostic potential in the three solid tumors: BRCA for distant metastasis-free survival in 86/29 (patients with low/high expression of SRGN) and 91/195 patients, respectively; lung cancer for RFS in 182/22 patients; and OV for OS in 87/46 patients. According to GEPIA, SRGN only had prognostic value in testicular germ cell tumors, glioblastoma multiforme, and SKCM. According to the TIMER gene_outcome module, SRGN was a significant prognostic factor for OS in patients with sarcoma, SKCM, and SKCM metastasis after correcting for age, race, sex, and tumor purity (Supplementary Figure 5, Table 1).

**Prognostic potential of SRGN expression associated with immune cells in pan-cancer**

We analyzed SRGN expression in relation to six immune infiltrates using TIMER 2.0. SRGN expression was a prognostic factor in cholangiocarcinoma (CHOL), LIHC, LUAD, SKCM-metastasis, and thymoma (THYM), correcting for six immune infiltrates and clinical factors (age, sex, stage, and tumor purity). High SRGN expression was a risk factor for OS in CHOL, LUAD, and THYM, but a favorable prognostic factor in LIHC and SKCM metastasis (Supplementary Table 7). Only SKCM metastasis was significantly different between the groups with low and high SRGN levels (log-rank test, P < 0.001; Supplementary Figure 8).

**Acknowledgements:** We thank International Science Editing (http://www. International scienceediting.com) for editing a draft of this manuscript.

**Disclosure Statement:** The authors declare no potential conflicts of interest.

**Ethics approval and consent to participate:** Not Applicable for patient’s ethics approval.

**References**

### 1. https://www.who.int/news-room/fact-sheets/detail/hepatitis-b

### 2. Gurtsevitch VE. Human oncogenic viruses: Hepatitis B and hepatitis C viruses and their role in hepatocarcinogenesis. Biochemistry (Moscow), 2008; 73(5): 504-513.

3. [Bertoletti A](https://www.ncbi.nlm.nih.gov/pubmed/?term=Bertoletti%20A%5BAuthor%5D&cauthor=true&cauthor_uid=27084039), [Ferrari C](https://www.ncbi.nlm.nih.gov/pubmed/?term=Ferrari%20C%5BAuthor%5D&cauthor=true&cauthor_uid=27084039). Adaptive immunity in HBV infection. [J Hepatol.](https://www.ncbi.nlm.nih.gov/pubmed/27084039) 2016; 64(1 Suppl):S71-S83.

11. Yongwei Li, Mingfen Zhu, Gang Li. Expression of serglycin in HepG2 cells with the different HBV infection status. Journal of Gastroenterology and Hepatology. 2013; 28( Supplement 3): 913.

12. Li Y, Chen H, Lu H, Zou Z, Li Y. [Prognostic significance of hematopoietic-cell serglycin for the survival of hepatocellular carcinoma: A single-center retrospective study.](https://pubmed.ncbi.nlm.nih.gov/33081679/) Comb Chem High Throughput Screen. 2021; 24(7):986-995.

13. Tang Z, Li C, Kang B, Gao G, Li C, Zhang Z. GEPIA: a web server for cancer and normal gene expression profiling and interactive analyses. Nucleic Acids Res. 2017; 45(W1):W98-W102.

14. Nagy A, Munkárcsy G, Győrffy B. Pancancer survival analysis of cancer hallmark genes, Scientific Reports, 2021; 11(1): 6047.

15. Li T, Fan J, Wang B, Traugh N, Chen Q, Liu JS, et al. TIMER: a web server for comprehensive analysis of tumor- infiltrating immune cells. Cancer Res. 2017; 77: e108-e110.

16．Mizuno, H., Kitada, K., Nakai, K. et al. PrognoScan: a new database for meta-analysis of the prognostic value of genes. BMC Med Genomics. 2009; 2:18.

 31. Zhang Z, Deng Y, Zheng G, Jia X, Xiong Y, Luo K, Qiu Q, Qiu N, Yin J, Lu M, Liu H, Gu Y, He Z.SRGN- TGFbeta2 regulatory loop confers invasion and metastasis in triple-negative breast cancer. Oncogenesis. 2017; 6(7):e360.
