## Supplementary material for "Important Role of Hematopoietic Proteoglycan Serglycin in Liver Hepatocellular Carcinoma Associated with Tumor Microenvironment": supplemmental: Supplementary 1,3-5,7.pdf

**Supplementary Figure 1** Differential expression of SRGN in cancers via GEPIA, *p<0.05.

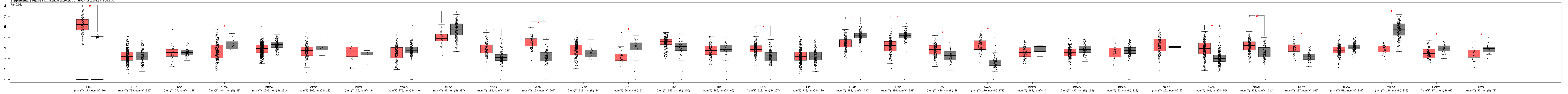

**Supplementary Figure 3** Kaplan-Meier survival curves of SRGN in pan-cancer via PrognoScan. red line, patients with high SRGN expression; blue line, patients with low SRGN expression. OS, overall survival; EFS, event-free survival; DMFS, distant metastasis-free survival; RFS, relapse-free survival; AML, acute myeloid leukemia; DLBCL, diffuse large B-cell lymphoma.

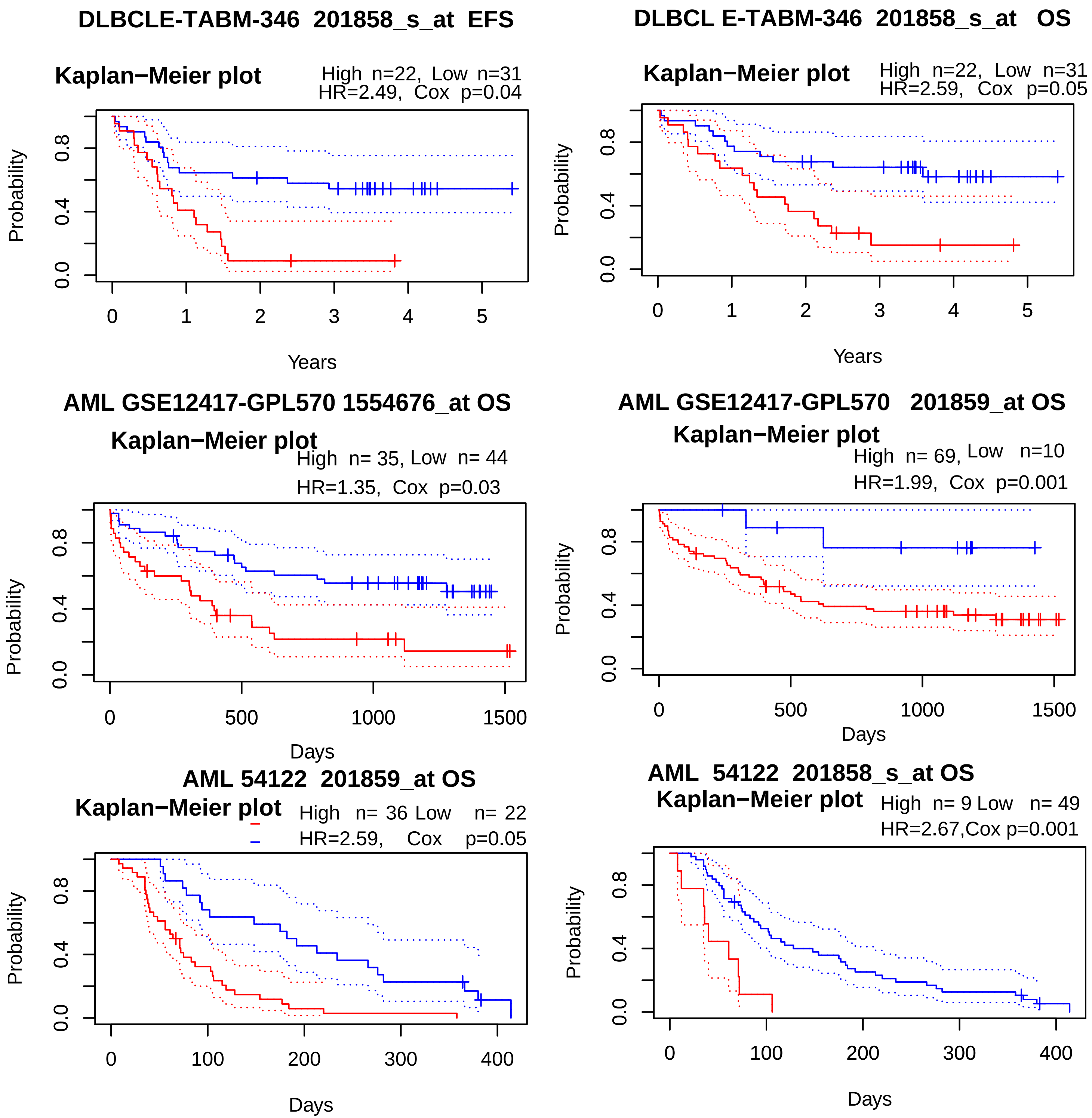
DLBCLE-TABM-346 201858_s_at EFS
Kaplan−Meier plot
High n=22, Low n=31
HR=2.49, Cox p=0.04
Probability
0.0 0.4 0.8
0 1 2 3 4 5
Years
DLBCL E-TABM-346 201858_s_at OS
Kaplan−Meier plot
High n=22, Low n=31
HR=2.59, Cox p=0.05
Probability
0.0 0.4 0.8
0 1 2 3 4 5
Years
AML GSE12417-GPL570 1554676_at OS
Kaplan−Meier plot
High n= 35, Low n= 44
HR=1.35, Cox p=0.03
Probability
0.0 0.4 0.8
0 500 1000 1500
Days
AML GSE12417-GPL570 201859_at OS
Kaplan−Meier plot
High n= 69, Low n=10
HR=1.99, Cox p=0.001
Probability
0.0 0.4 0.8
0 500 1000 1500
Days
AML 54122 201859_at OS
Kaplan−Meier plot
High n= 36 Low n= 22
HR=2.59, Cox p=0.05
Probability
0.0 0.4 0.8
0 100 200 300 400
Days
AML 54122 201858_s_at OS
Kaplan−Meier plot
High n= 9 Low n= 49
HR=2.67,Cox p=0.001
Probability
0.0 0.4 0.8
0 100 200 300 400
Days

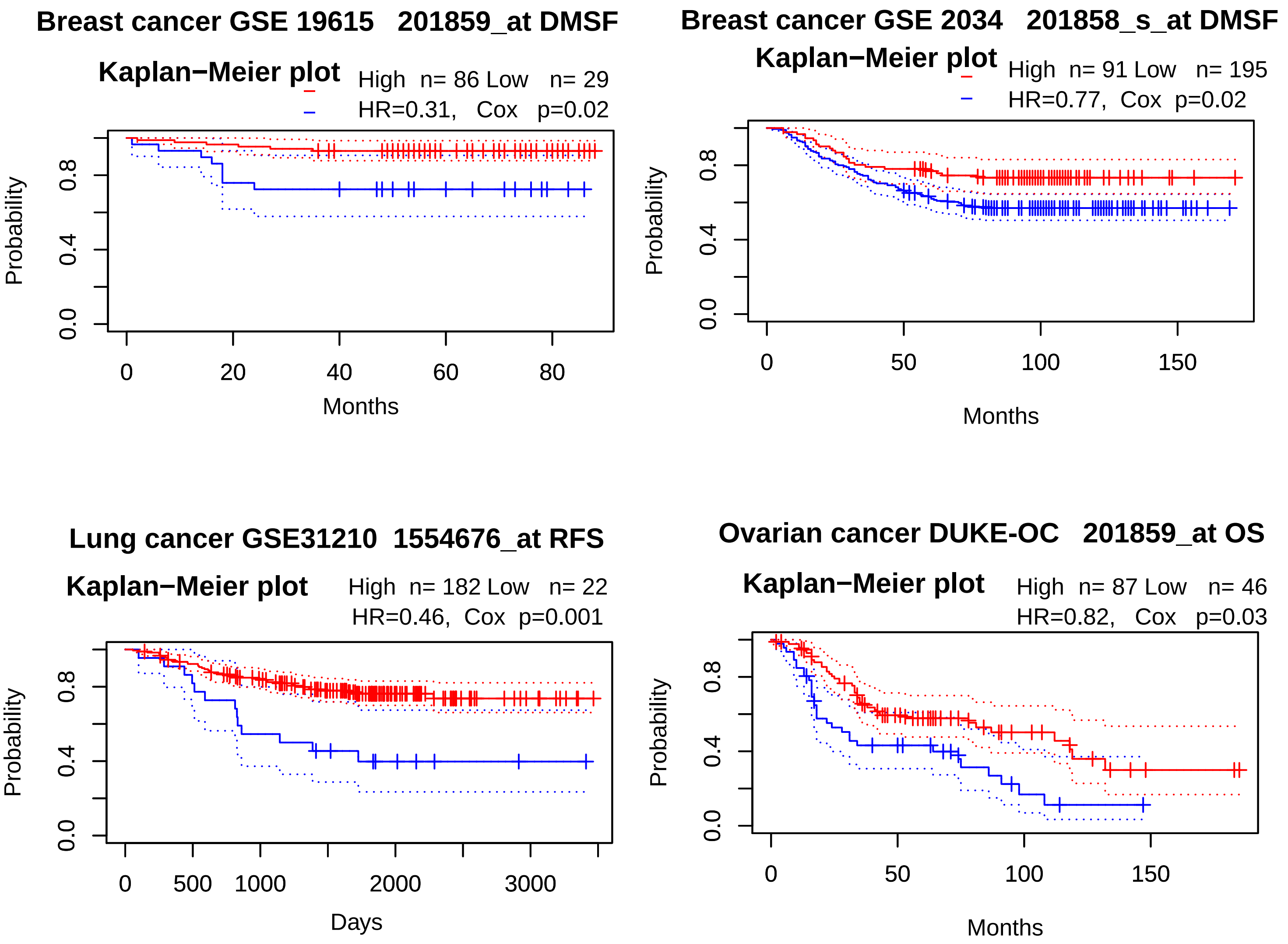
Breast cancer GSE 19615 201859_at DMSF
Kaplan−Meier plot
High n= 86 Low n= 29
HR=0.31, Cox p=0.02
Probability
0.0
0.4
0.8
0
20
40
60
80
Months
Breast cancer GSE 2034 201858_s_at DMSF
Kaplan−Meier plot
High n= 91 Low n= 195
HR=0.77, Cox p=0.02
Probability
0.0
0.4
0.8
0
50
100
150
Months
Lung cancer GSE31210 1554676_at RFS
Kaplan−Meier plot
High n= 182 Low n= 22
HR=0.46, Cox p=0.001
Probability
0.0
0.4
0.8
0
500
1000
2000
3000
Days
Ovarian cancer DUKE-OC 201859_at OS
Kaplan−Meier plot
High n= 87 Low n= 46
HR=0.82, Cox p=0.03
Probability
0.0
0.4
0.8
0
50
100
150
Months

**Supplementary Figure 4** Kaplan-Meier survival curves of SRGN in pan-cancer via GEPIA. High SRGN levels was associated with poor prognosis for OS in TGCT (The 95% CI as dotted line, HR(high) = 9.1e8, p(HR)=1, p=0.022), and poor RFS (Disease Free Survival) in GBM (glioblastoma multiforme, HR(high) =1.7, p(HR)=0.014, p=0.013), while with favorable prognosis for OS and RFS in SKCM (skin cutaneous melanoma, HR(high)=0.51, p(HR)=8.2e-07, p=5.2e-07; HR(high) =0.74, p(HR)0.015, p=0.014, respectively). n (high or low): the number of patients with high or low expression of SRGN.

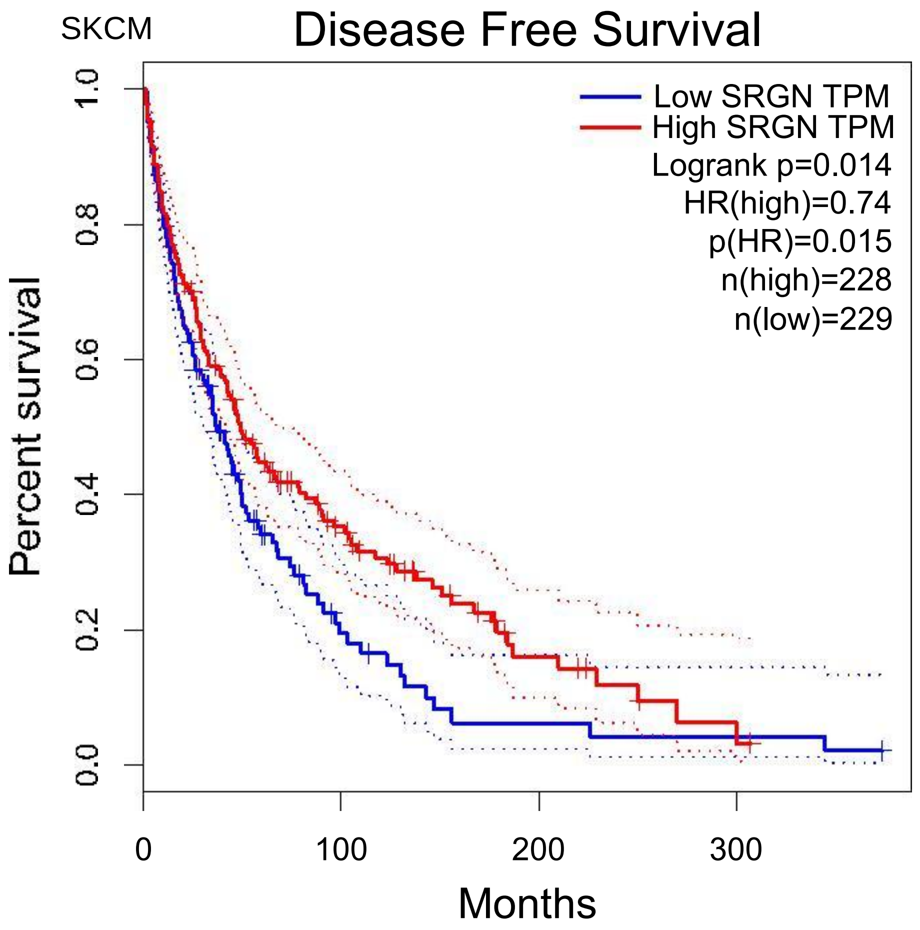
SKCM
Disease Free Survival
Low SRGN TPM
High SRGN TPM
Logrank p=0.014
HR(high)=0.74
p(HR)=0.015
n(high)=228
n(low)=229
Percent survival
0.0
0.2
0.4
0.6
0.8
1.0
0
100
200
300
Months

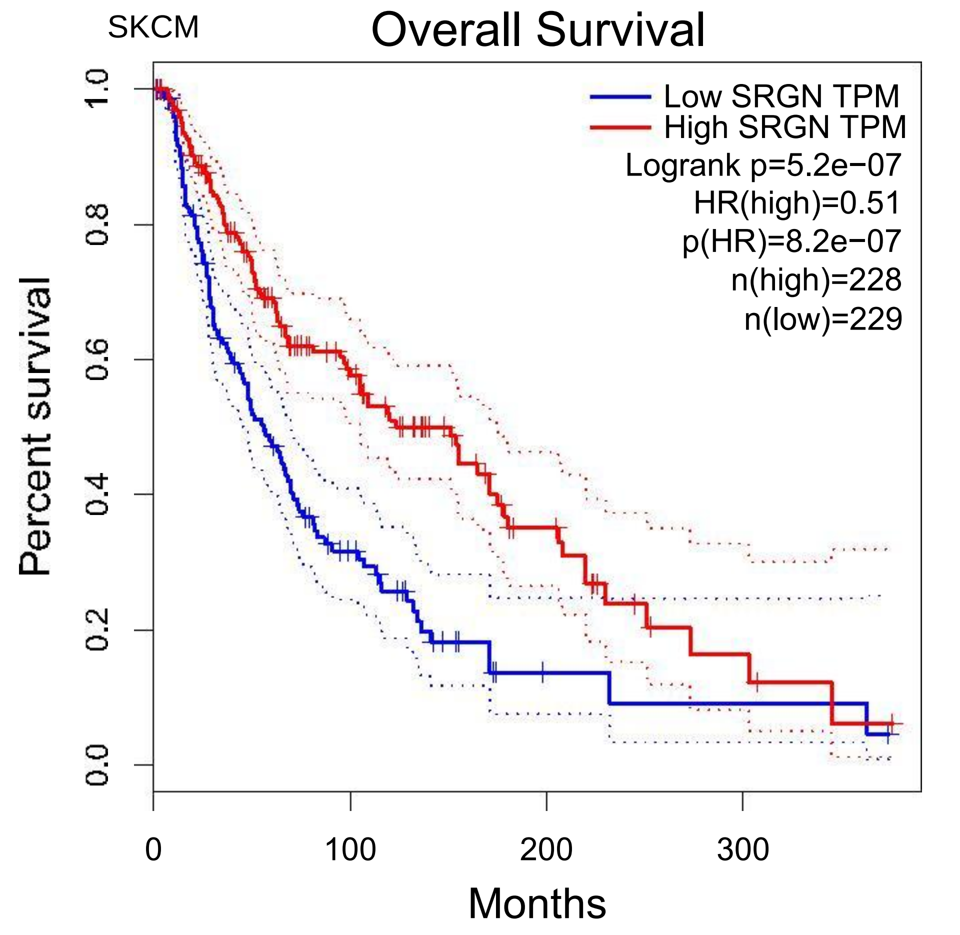
SKCM
Overall Survival
Low SRGN TPM
High SRGN TPM
Logrank p=5.2e−07
HR(high)=0.51
p(HR)=8.2e−07
n(high)=228
n(low)=229
Percent survival
1.0
0.8
0.6
0.4
0.2
0.0
0
100
200
300
Months

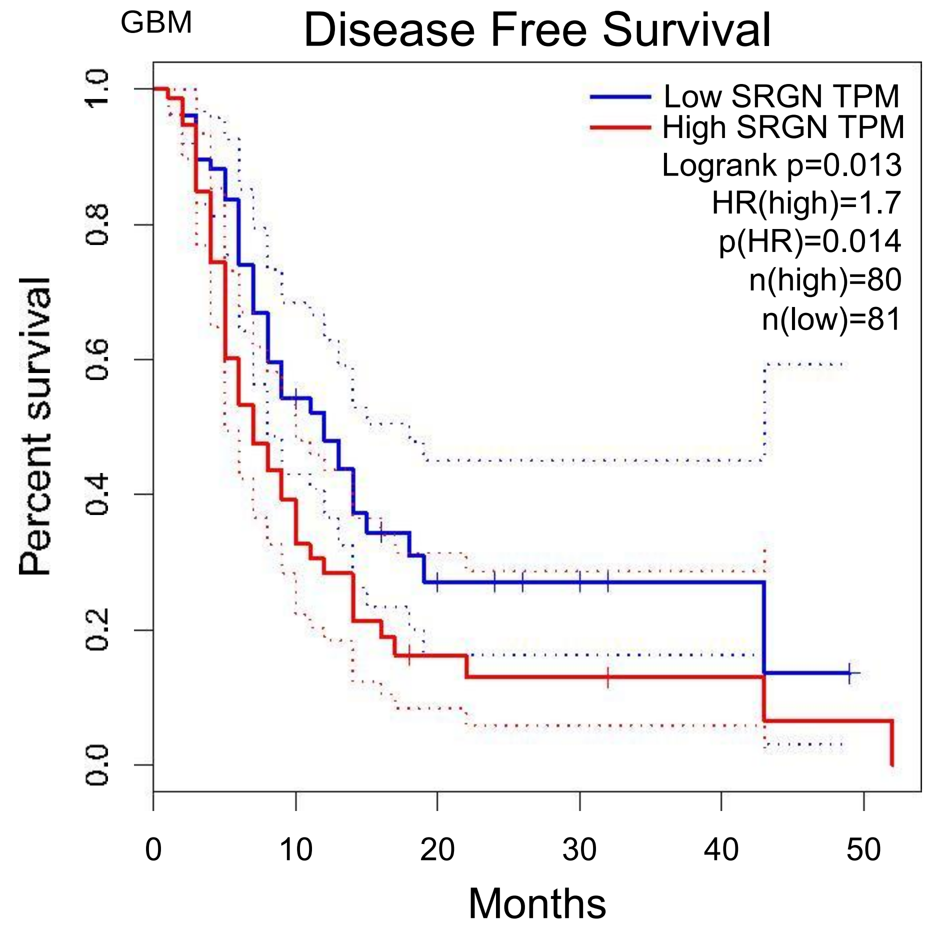
GBM
Disease Free Survival
Low SRGN TPM
High SRGN TPM
Logrank p=0.013
HR(high)=1.7
p(HR)=0.014
n(high)=80
n(low)=81
Percent survival
0.0
0.2
0.4
0.6
0.8
1.0
0
10
20
30
40
50
Months

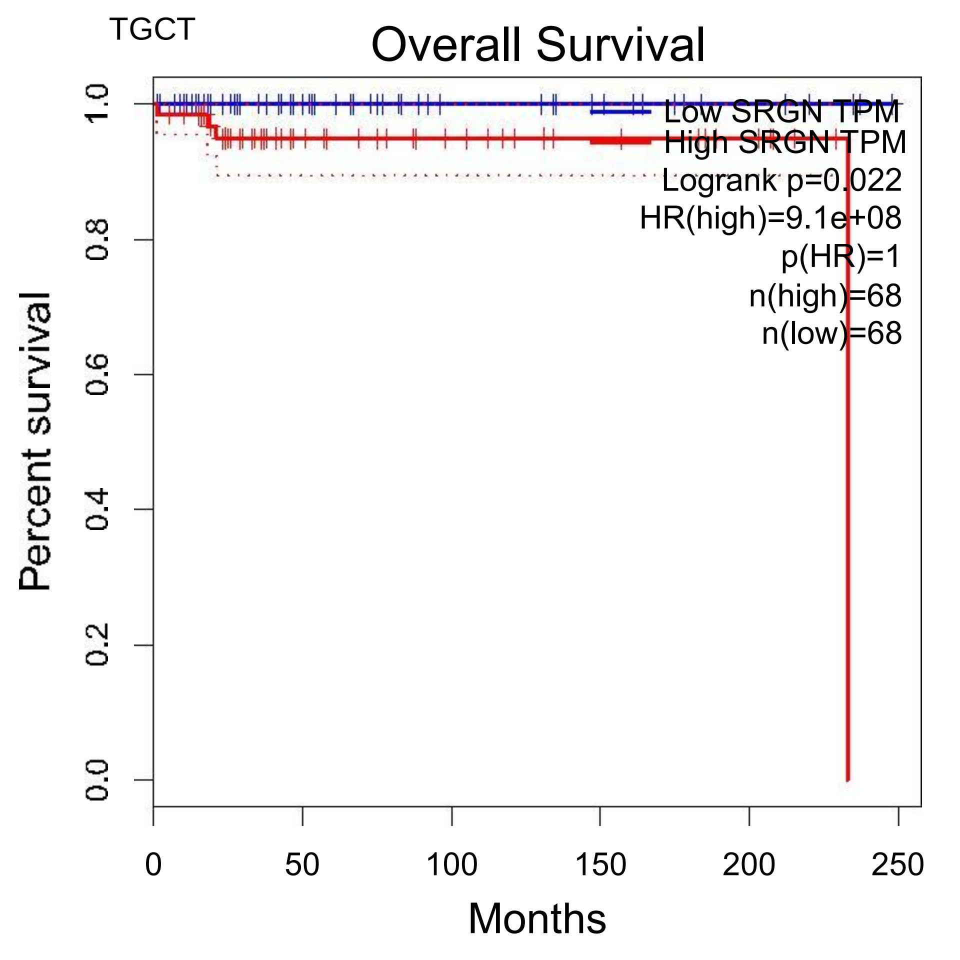
TGCT
Overall Survival
Low SRGN TPM
High SRGN TPM
Logrank p=0.022
HR(high)=9.1e+08
p(HR)=1
n(high)=68
n(low)=68
Percent survival
0.0
0.2
0.4
0.6
0.8
1.0
0
50
100
150
200
250
Months

**Supplementary Figure 5** Kaplan-Meier survival curves of SRGN in pan-cancer viagene_outcome module of TIMER. The Coxph model showed the OS with SRGN, age, race, gender, and purity. SRGN was a protective factor in SARC, SKCM and SKCM metastasis (HR=0.762, P=0.0213; HR=0.746, P=0.000873; HR=0.734, P=0.00184, respectively). ACC (Adrenocortical carcinoma) and LGG (Lower Grade Glima) had marginally worse prognosis with higher SRGN levels (HR=1.49, p=0.0659 and HR=1.19, p=0.0691, respectively).

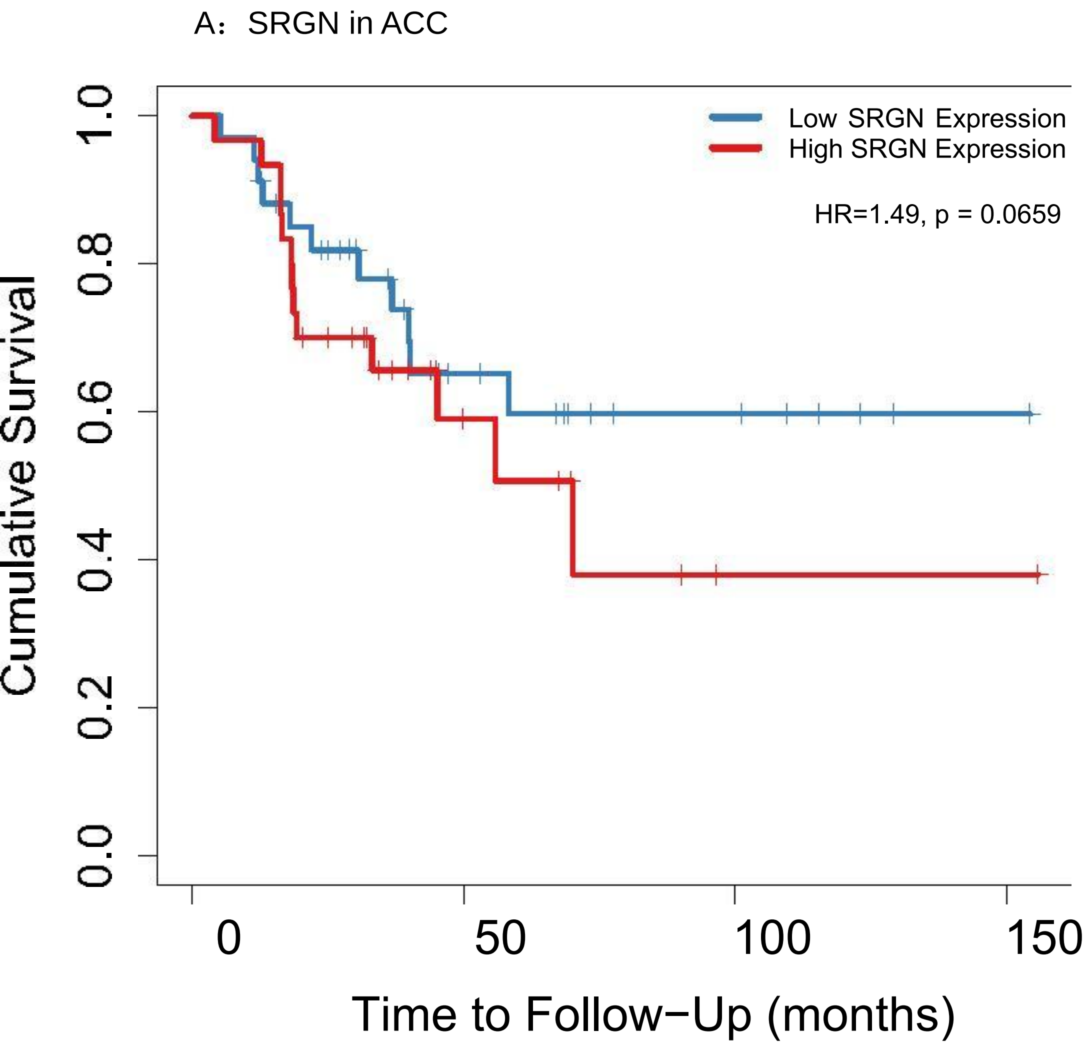
A：SRGN in ACC
Low SRGN Expression
High SRGN Expression
HR=1.49, p = 0.0659
Cumulative Survival
1.0
0.8
0.6
0.4
0.2
0.0
0
50
100
150
Time to Follow−Up (months)

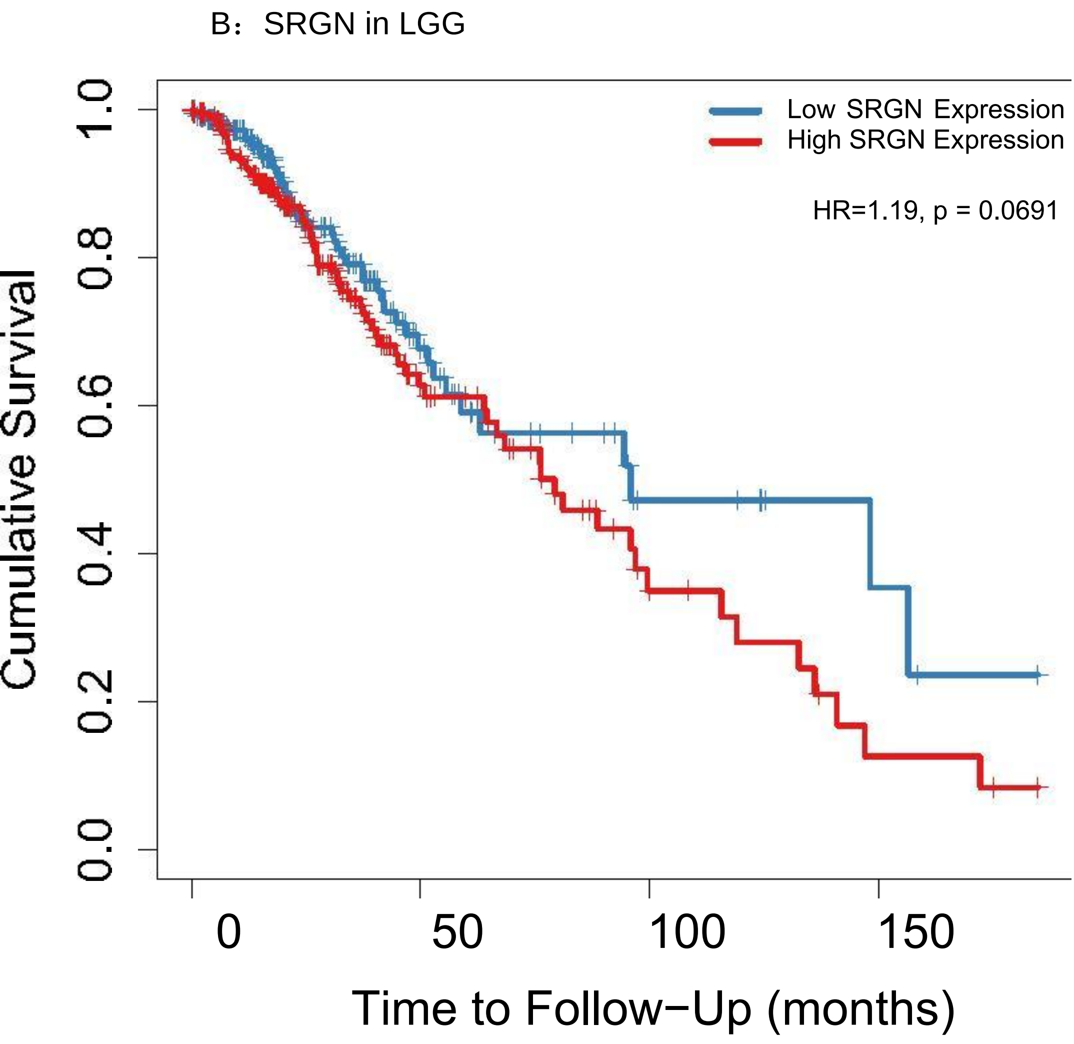

B：SRGN in LGG
Low SRGN Expression
High SRGN Expression
HR=1.19, p = 0.0691
Cumulative Survival
1.0
0.8
0.6
0.4
0.2
0.0
0
50
100
150
Time to Follow−Up (months)

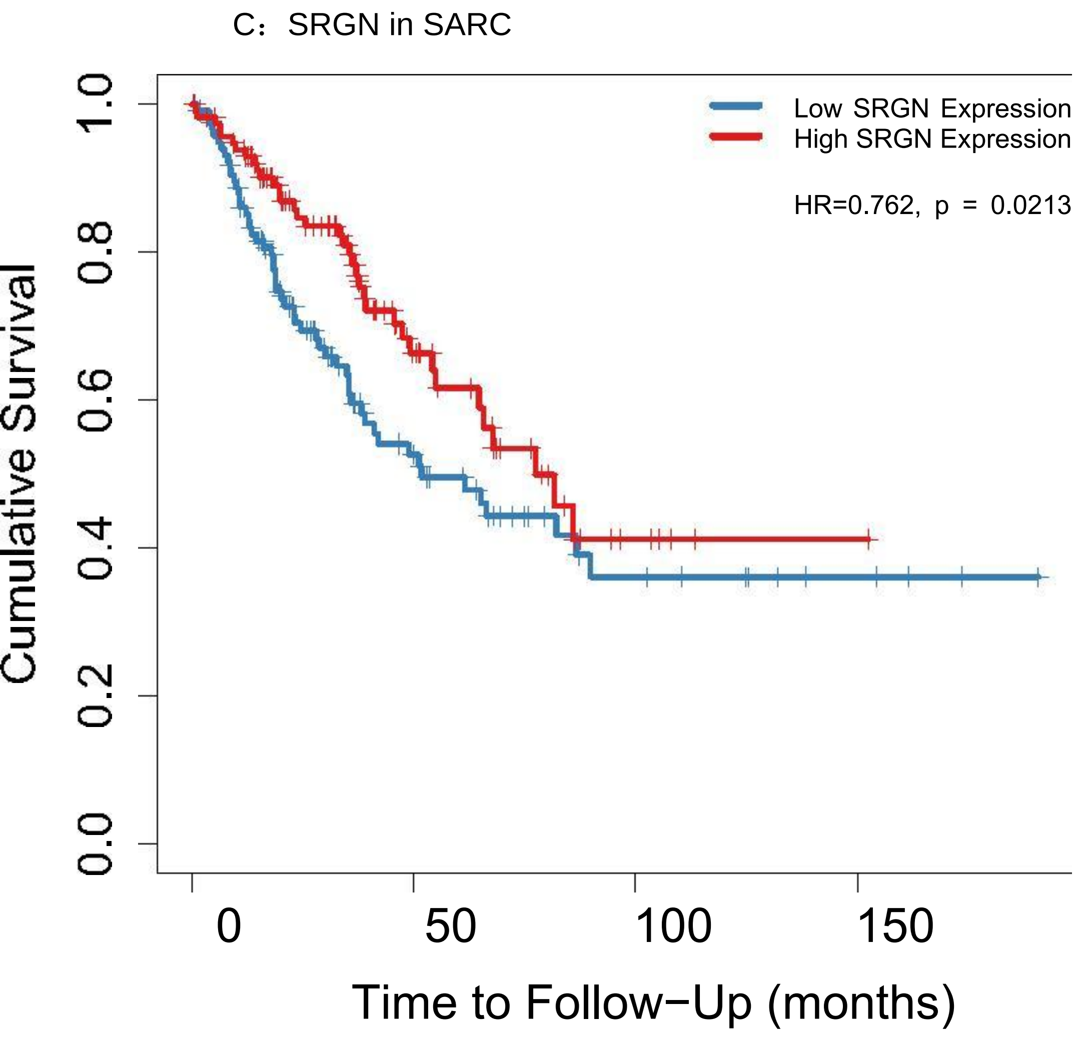

C：SRGN in SARC
Low SRGN Expression
High SRGN Expression
HR=0.762, p = 0.0213
Cumulative Survival
0.0
0.2
0.4
0.6
0.8
1.0
0
50
100
150
Time to Follow−Up (months)

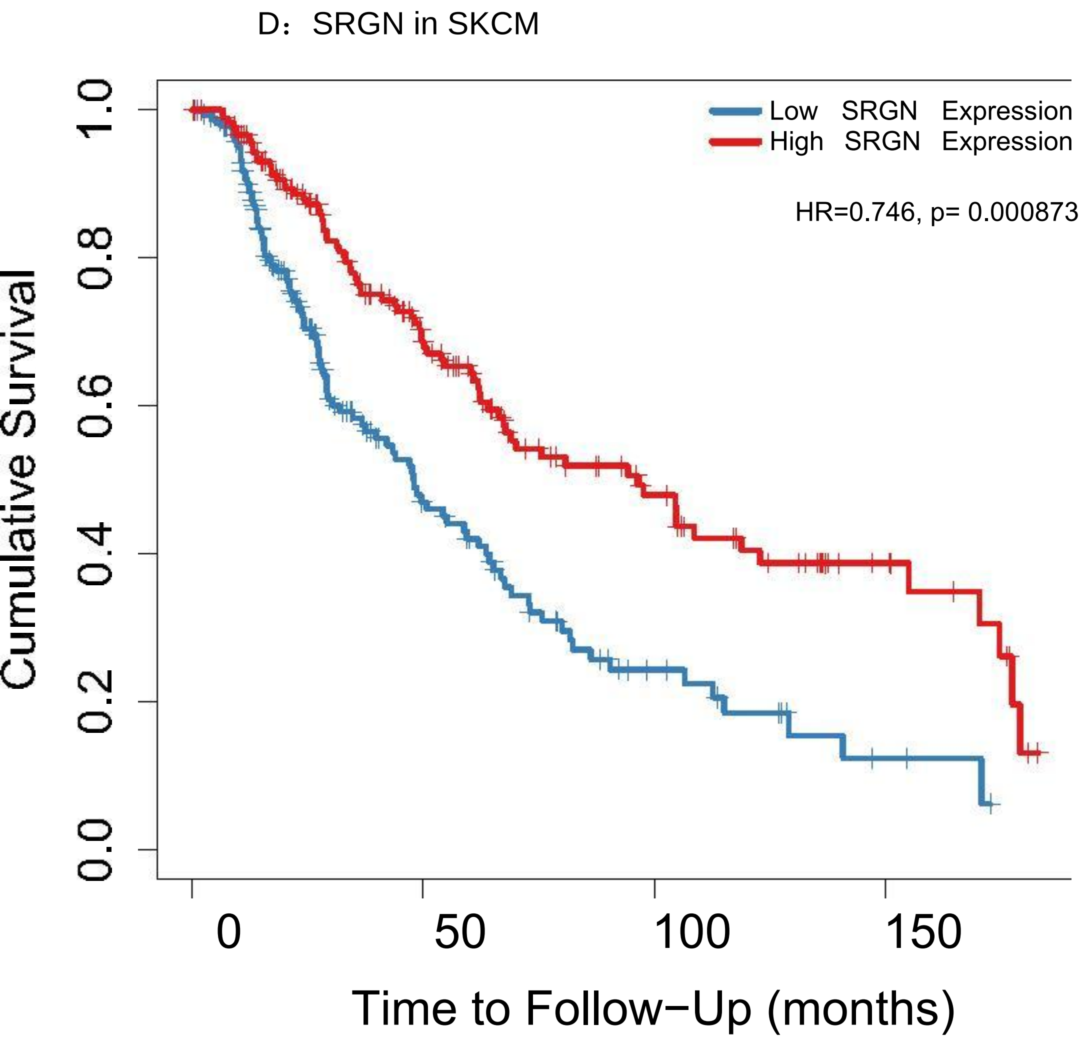
D：SRGN in SKCM
Low SRGN Expression
High SRGN Expression
HR=0.746, p= 0.000873
Cumulative Survival
0.0
0.2
0.4
0.6
0.8
1.0
0
50
100
150
Time to Follow−Up (months)

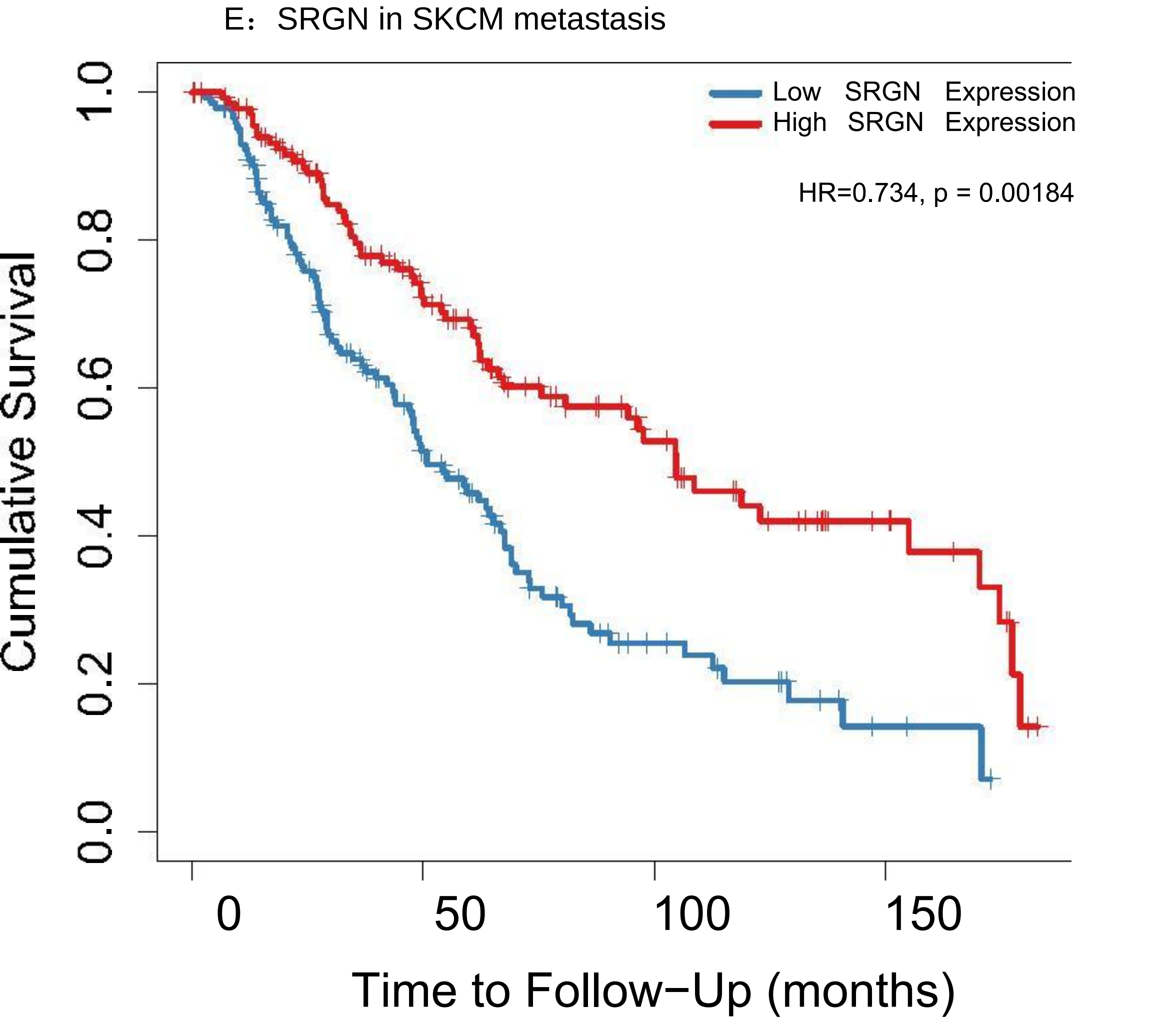

E：SRGN in SKCM metastasis
Low SRGN Expression
High SRGN Expression
HR=0.734, p = 0.00184
Cumulative Survival
0.0
0.2
0.4
0.6
0.8
1.0
0
50
100
150
Time to Follow−Up (months)

**Supplementary Table 7** The Coxph model of CHOL, LIHC, LUAD, SKCM metastasis and THYM via TIMER, Survival module. “Survival” module builds a multivariable Cox proportional hazard model with tumor immune subsets（B Cell CD8+ T Cell CD4+ T Cell Macrophage Neutrophil Dendritic Cell）, correcting for multiple covariates including clinical factors (age, gender, ethnicity, tumor stages) and SRGN expression by function coxph() from R package 'survival'. Abbreviation: CHOL (Cholangiocarcinoma) LIHC (Liver Hepatocellular Carcinoma) LUAD (Lung Adenocarcinoma) SKCM-Metastasis THYM (Thymoma).

```
Model: Surv(CHOL) ~ Age + Gender + Race + Stage + Purity + B_cell + CD8_Tcell + CD4_Tcell + Mac
36 patients with 18 dying
                coef            HR 95%CI_l       95%CI_u p.value sig
Age            0.042  1.042000e+00   0.986  1.102000e+00   0.144
gendermale     1.114  3.045000e+00   0.577  1.606400e+01   0.189
raceBlack      1.558  4.751000e+00   0.124  1.814160e+02   0.402
raceWhite     -3.264  3.800000e-02   0.003  5.750000e-01   0.018   *
stage2         1.130  3.095000e+00   0.553  1.733300e+01   0.199
stage3        25.621  1.339568e+11   0.000           Inf   0.998
stage4         1.647  5.193000e+00   1.010  2.668900e+01   0.049   *
Purity         1.475  4.372000e+00   0.011  1.786918e+03   0.631
B_cell      -520.581  0.000000e+00   0.000  0.000000e+00   0.034   *
CD8_Tcell    218.442  7.382572e+94   0.000 8.075749e+226   0.159
CD4_Tcell     72.948  4.798699e+31   0.000 1.267674e+167   0.647
Macrophage   -66.088  0.000000e+00   0.000 1.682044e+154   0.758
Neutrophil -2279.927  0.000000e+00   0.000  0.000000e+00   0.012   *
Dendritic    359.741 1.712938e+156   0.000           Inf   0.142
SRGN           1.243  3.466000e+00   1.066  1.126800e+01   0.039   *
Rsquare= 0.516 (max possible= 9.46e-01 )
Likelihood ratio test p= 3.64e-02
Wald test p= 3.51e-01
Score (logrank) test p= 1.06e-01

Model: Surv(LIHC) ~ Age + Gender + Race + Stage + Purity + B_cell + CD8_Tcell + CD4_Tcell + Mac
305 patients with 101 dying
              coef        HR 95%CI_l      95%CI_u p.value sig
Age          0.010     1.010   0.992        1.027   0.282
gendermale   0.010     1.010   0.620        1.644   0.969
raceBlack    1.067     2.907   1.037        8.146   0.042   *
raceWhite    0.103     1.109   0.659        1.865   0.697
stage2       0.157     1.170   0.689        1.987   0.561
stage3       0.696     2.007   1.228        3.278   0.005  **
stage4       1.529     4.614   1.334       15.966   0.016   *
Purity       0.791     2.205   0.711        6.836   0.171
B_cell      -6.761     0.001   0.000        2.924   0.091   .
CD8_Tcell   -6.579     0.001   0.000        0.232   0.012   *
CD4_Tcell  -10.121     0.000   0.000        0.099   0.011   *
Macrophage  11.312 81771.729 190.969 35014156.817   0.000 ***
Neutrophil   4.170    64.737   0.000 19055733.934   0.516
Dendritic    7.124  1241.229  17.190    89622.916   0.001  **
SRGN        -0.433     0.648   0.476        0.883   0.006  **
Rsquare= 0.153 (max possible= 9.62e-01 )
Likelihood ratio test p= 9.63e-06
Wald test p= 4.75e-06
Score (logrank) test p= 1.45e-06

Model: Surv(LUAD) ~ Age + Gender + Race + Stage + Purity + B_cell + CD8_Tcell + CD4_Tcell + Mac
391 patients with 138 dying
             coef           HR 95%CI_l 95%CI_u p.value sig
Age         0.016        1.016   0.996   1.036   0.111
gendermale -0.140        0.870   0.607   1.246   0.446
raceBlack  16.280 11754419.246   0.000     Inf   0.994
raceWhite  16.383 13031367.747   0.000     Inf   0.994
stage2      0.862        2.367   1.529   3.664   0.000 ***
stage3      1.004        2.729   1.733   4.296   0.000 ***
stage4      1.205        3.336   1.679   6.628   0.001  **
Purity      0.389        1.476   0.595   3.659   0.401
```

```
B_cell     -3.069        0.046   0.003   0.756   0.031   *
CD8_Tcell  -1.173        0.309   0.034   2.843   0.300
CD4_Tcell   2.224        9.242   0.505 169.216   0.134
Macrophage -0.942        0.390   0.017   8.817   0.554
Neutrophil -3.120        0.044   0.001   3.481   0.161
Dendritic  -0.112        0.894   0.212   3.764   0.879
SRGN        0.322        1.379   1.138   1.671   0.001  **
Rsquare= 0.145 (max possible= 9.71e-01 )
Likelihood ratio test p= 1.65e-07
Wald test p= 3.19e-06
Score (logrank) test p= 3.74e-07

Model: Surv(SKCM-Metastasis) ~ Age + Gender + Race + Stage + Purity + B_cell + CD8_Tcell + CD4_
269 patients with 153 dying
             coef     HR 95%CI_l 95%CI_u p.value sig
Age         0.019  1.019   1.008   1.031   0.001  **
gendermale -0.171  0.843   0.589   1.207   0.350
raceWhite  -0.891  0.410   0.118   1.424   0.161
stage2     -0.001  0.999   0.622   1.605   0.998
stage3      0.484  1.622   1.053   2.500   0.028   *
stage4      1.323  3.756   1.678   8.408   0.001  **
Purity     -0.514  0.598   0.211   1.698   0.334
B_cell      1.182  3.262   0.058 183.053   0.565
CD8_Tcell   0.060  1.061   0.064  17.545   0.967
CD4_Tcell   0.421  1.524   0.047  48.948   0.812
Macrophage  2.553 12.846   0.877 188.183   0.062   ·
Neutrophil -2.269  0.103   0.000 451.818   0.596
Dendritic  -2.174  0.114   0.014   0.918   0.041   *
SRGN       -0.230  0.795   0.645   0.980   0.032   *
Rsquare= 0.214 (max possible= 9.95e-01 )
Likelihood ratio test p= 1.59e-08
Wald test p= 3.38e-08
Score (logrank) test p= 9.04e-09

Model: Surv(THYM) ~ Age + Gender + Race + Purity + B_cell + CD8_Tcell + CD4_Tcell + Macrophage
109 patients with 9 dying
              coef           HR 95%CI_l      95%CI_u p.value sig
Age          0.048        1.049   0.971 1.133000e+00   0.222
gendermale   0.122        1.129   0.187 6.821000e+00   0.895
raceBlack  -15.624        0.000   0.000          Inf   0.999
raceWhite   -0.131        0.877   0.077 9.957000e+00   0.916
Purity      -1.095        0.335   0.014 8.082000e+00   0.500
B_cell     -18.097        0.000   0.000 1.298000e+00   0.053   ·
CD8_Tcell   16.174 10573846.751   0.065 1.725711e+15   0.094   ·
CD4_Tcell    0.269        1.309   0.000 5.597848e+06   0.972
Macrophage  -8.319        0.000   0.000 4.037338e+05   0.442
Neutrophil -48.864        0.000   0.000 1.569000e+00   0.052   ·
Dendritic  -10.363        0.000   0.000 6.361620e+04   0.343
SRGN         2.105        8.205   1.432 4.701500e+01   0.018   *
Rsquare= 0.165 (max possible= 4.59e-01 )
Likelihood ratio test p= 7.52e-02
Wald test p= 4.94e-01
Score (logrank) test p= 6.65e-02
```
