## Supplementary material for "Important Role of Hematopoietic Proteoglycan Serglycin in Liver Hepatocellular Carcinoma Associated with Tumor Microenvironment": supplemmental: Supplementary 2,6,8-12 .pptx

### Slide 1
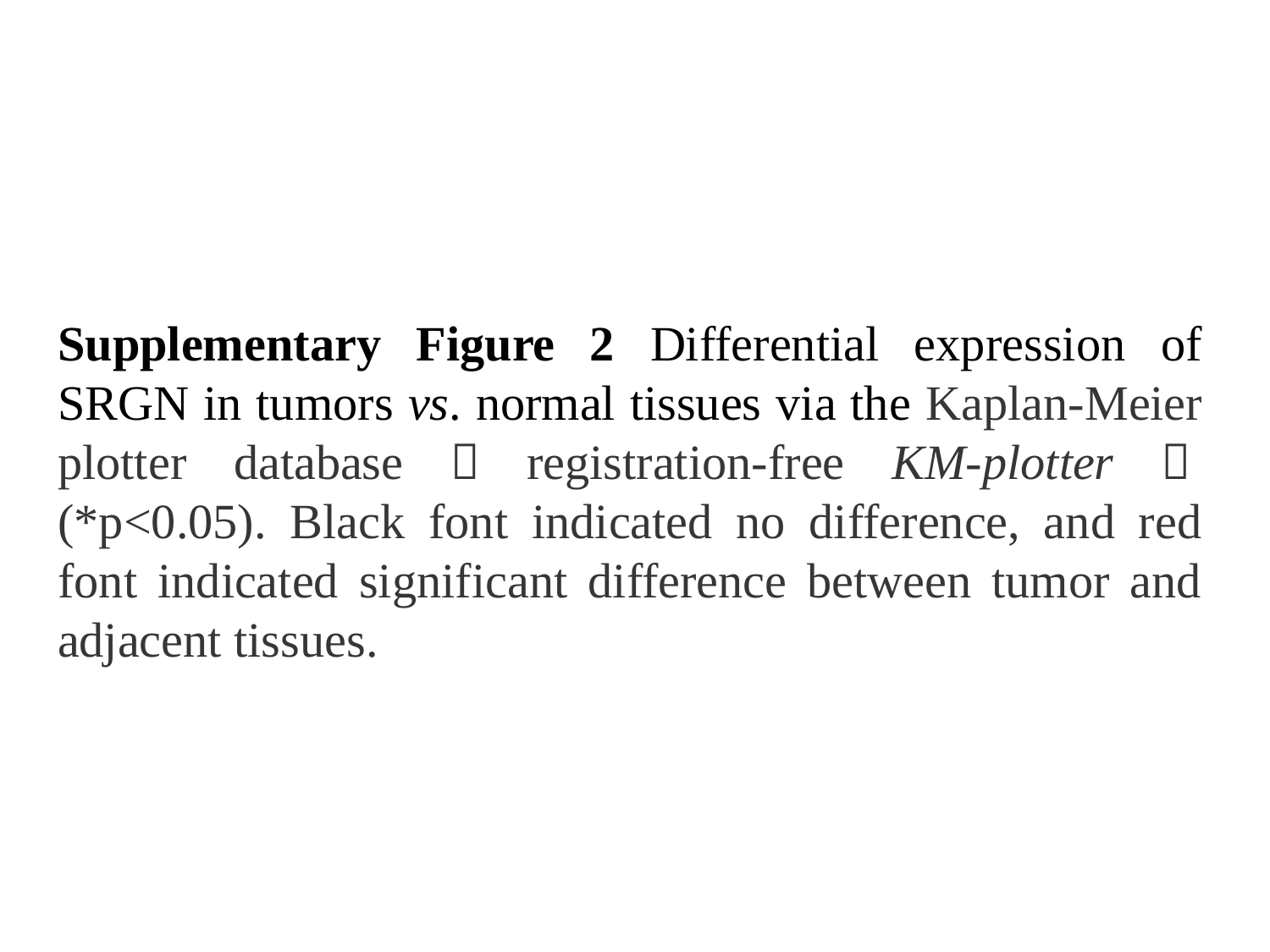

Supplementary Figure 2 Differential expression of SRGN in tumors vs. normal tissues via the Kaplan-Meier plotter database（registration-free KM-plotter）(*p<0.05). Black font indicated no difference, and red font indicated significant difference between tumor and adjacent tissues.

### Slide 2
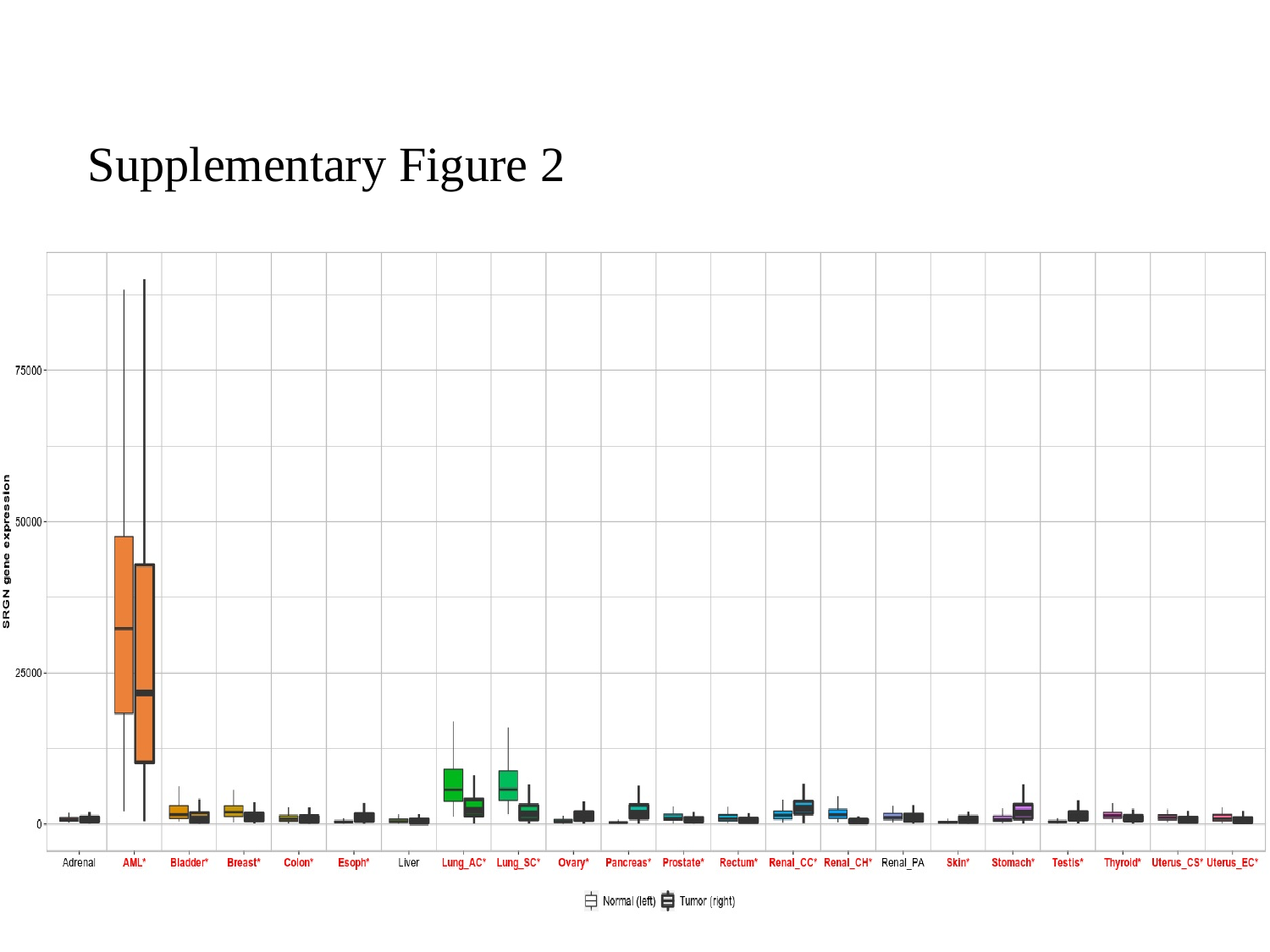

Supplementary Figure 2

### Slide 3
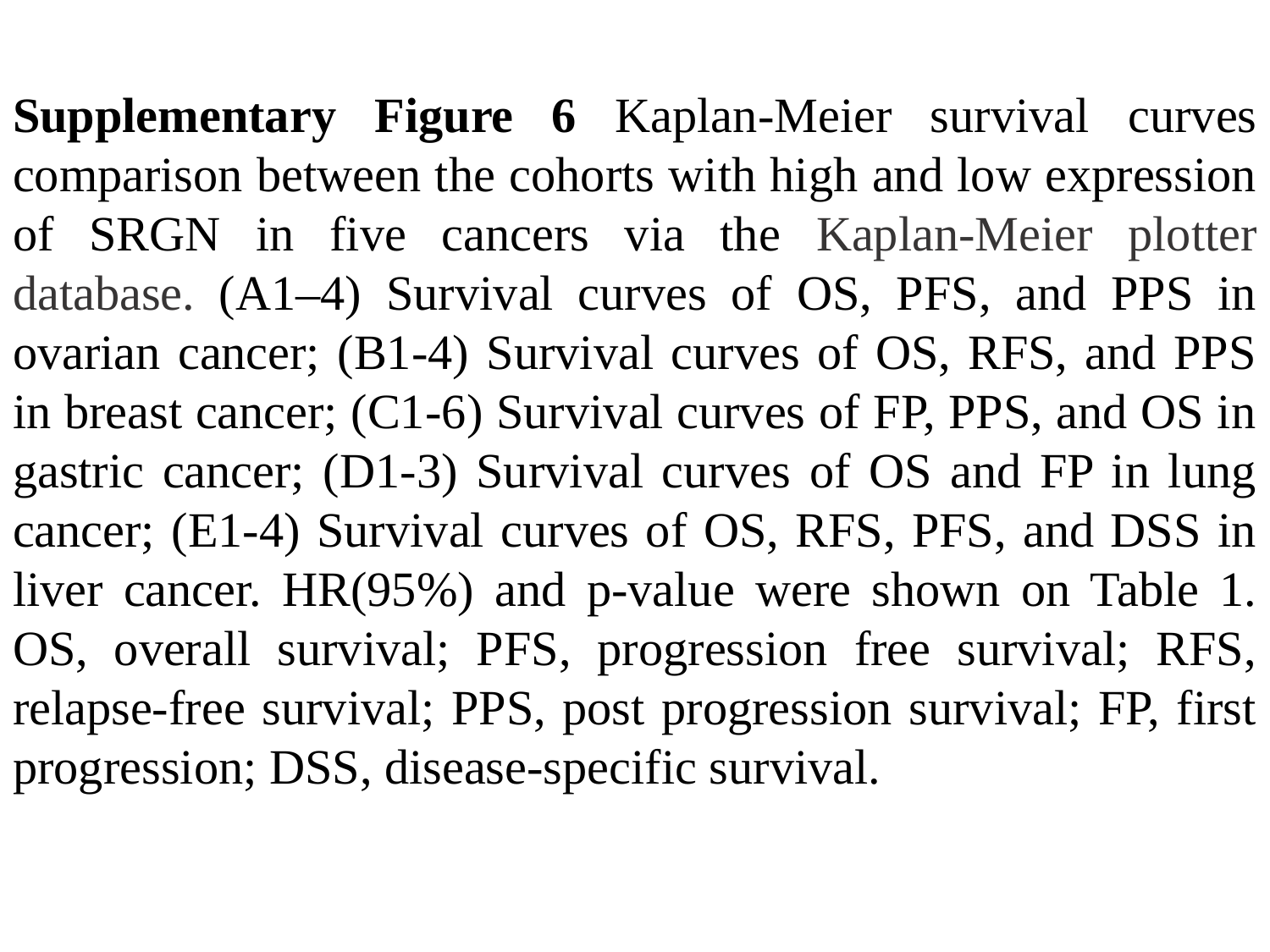

Supplementary Figure 6 Kaplan-Meier survival curves comparison between the cohorts with high and low expression of SRGN in five cancers via the Kaplan-Meier plotter database. (A1–4) Survival curves of OS, PFS, and PPS in ovarian cancer; (B1-4) Survival curves of OS, RFS, and PPS in breast cancer; (C1-6) Survival curves of FP, PPS, and OS in gastric cancer; (D1-3) Survival curves of OS and FP in lung cancer; (E1-4) Survival curves of OS, RFS, PFS, and DSS in liver cancer. HR(95%) and p-value were shown on Table 1. OS, overall survival; PFS, progression free survival; RFS, relapse-free survival; PPS, post progression survival; FP, first progression; DSS, disease-specific survival.

### Slide 4
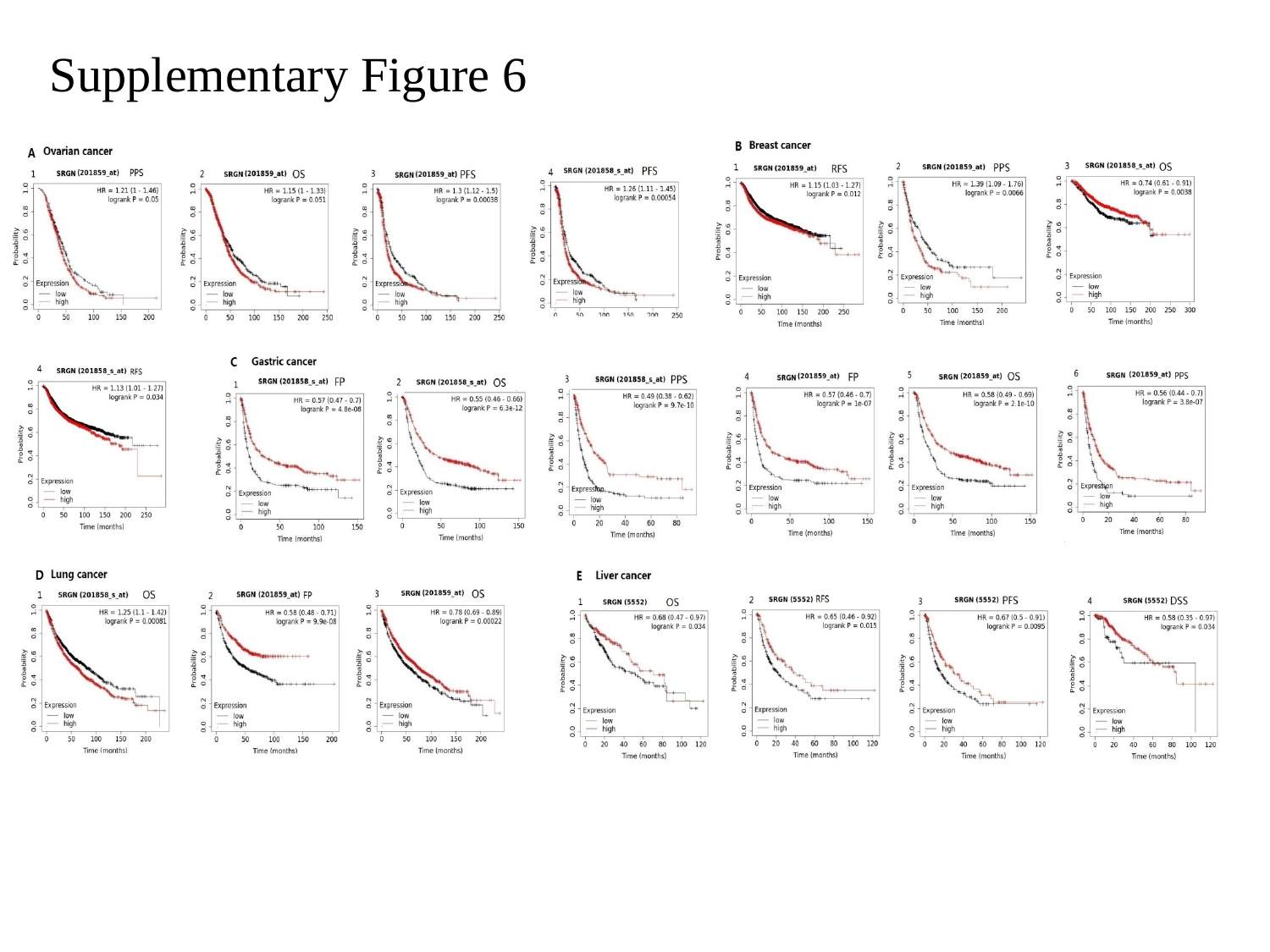

Supplementary Figure 6

### Slide 5
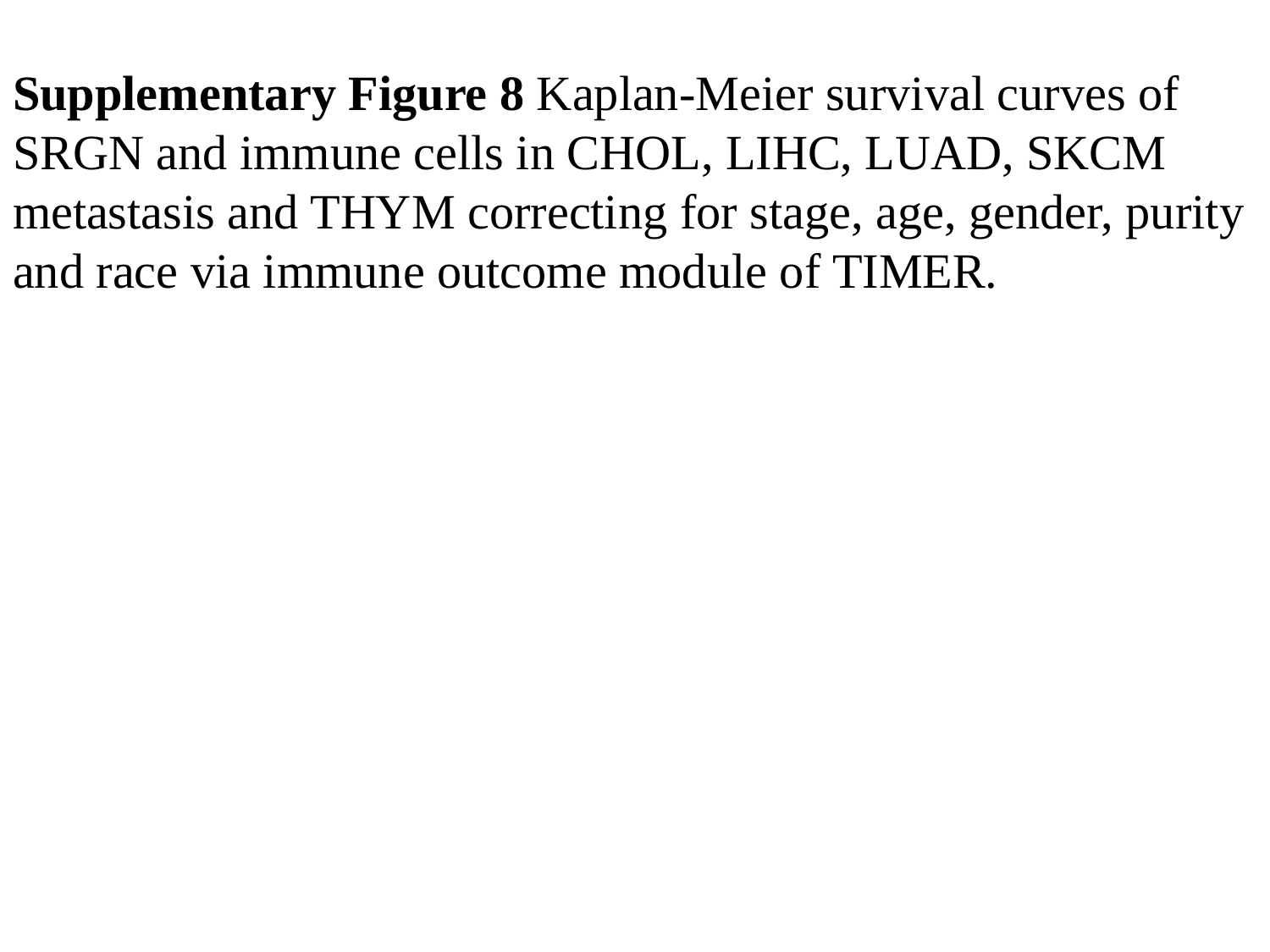

Supplementary Figure 8 Kaplan-Meier survival curves of SRGN and immune cells in CHOL, LIHC, LUAD, SKCM metastasis and THYM correcting for stage, age, gender, purity and race via immune outcome module of TIMER.

### Slide 6
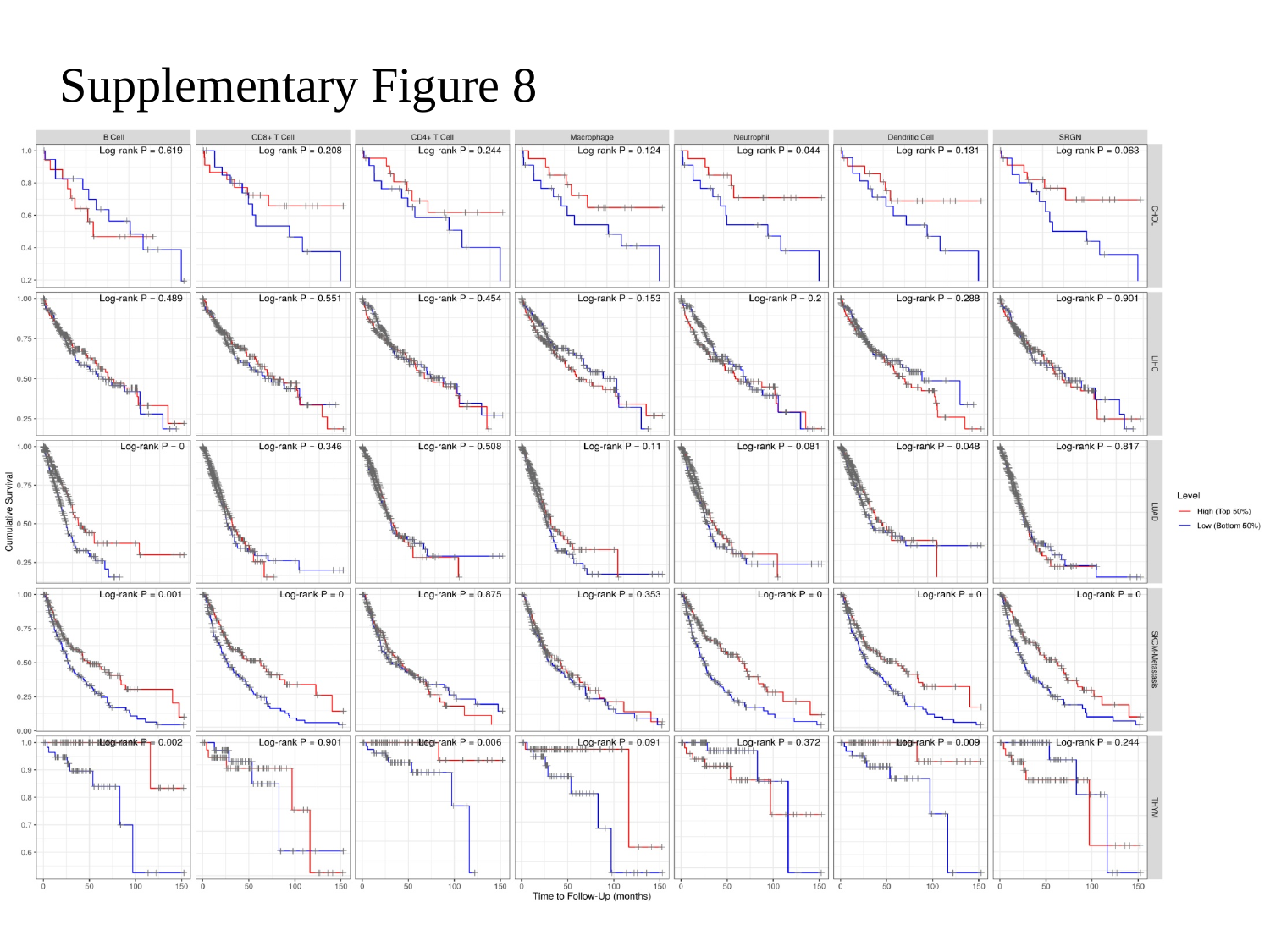

Supplementary Figure 8

### Slide 7
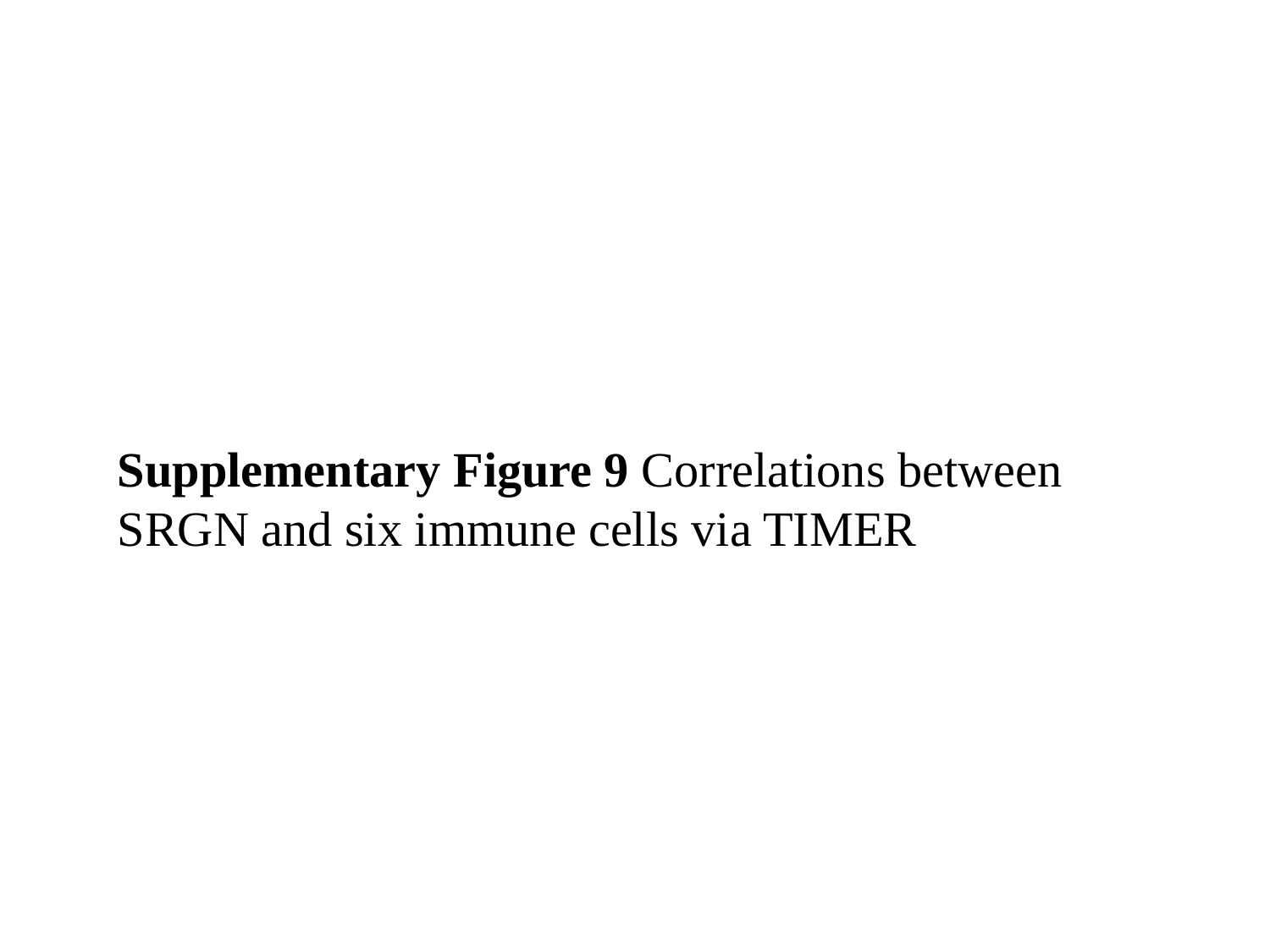

Supplementary Figure 9 Correlations between SRGN and six immune cells via TIMER

### Slide 8
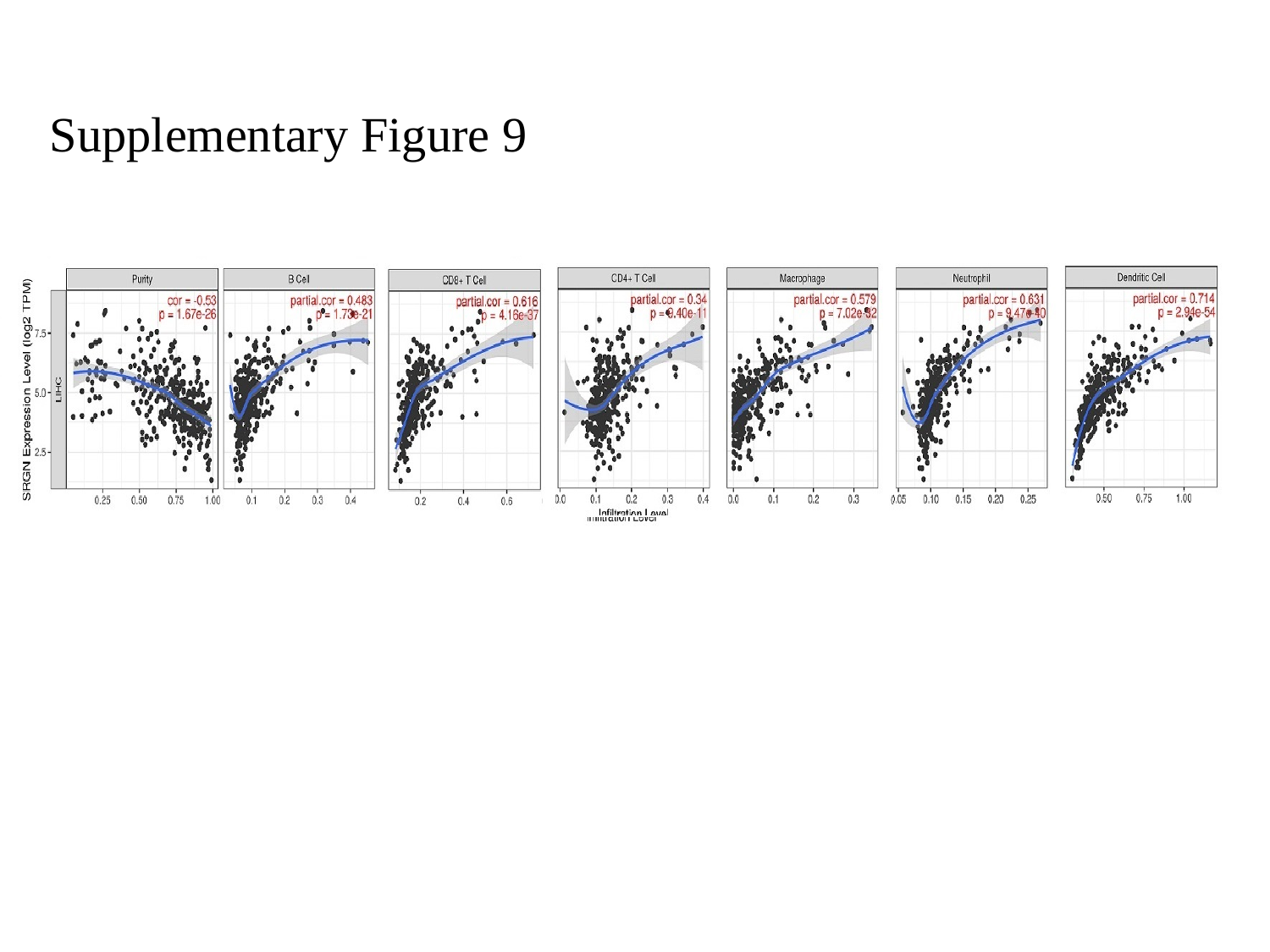

Supplementary Figure 9

### Slide 9
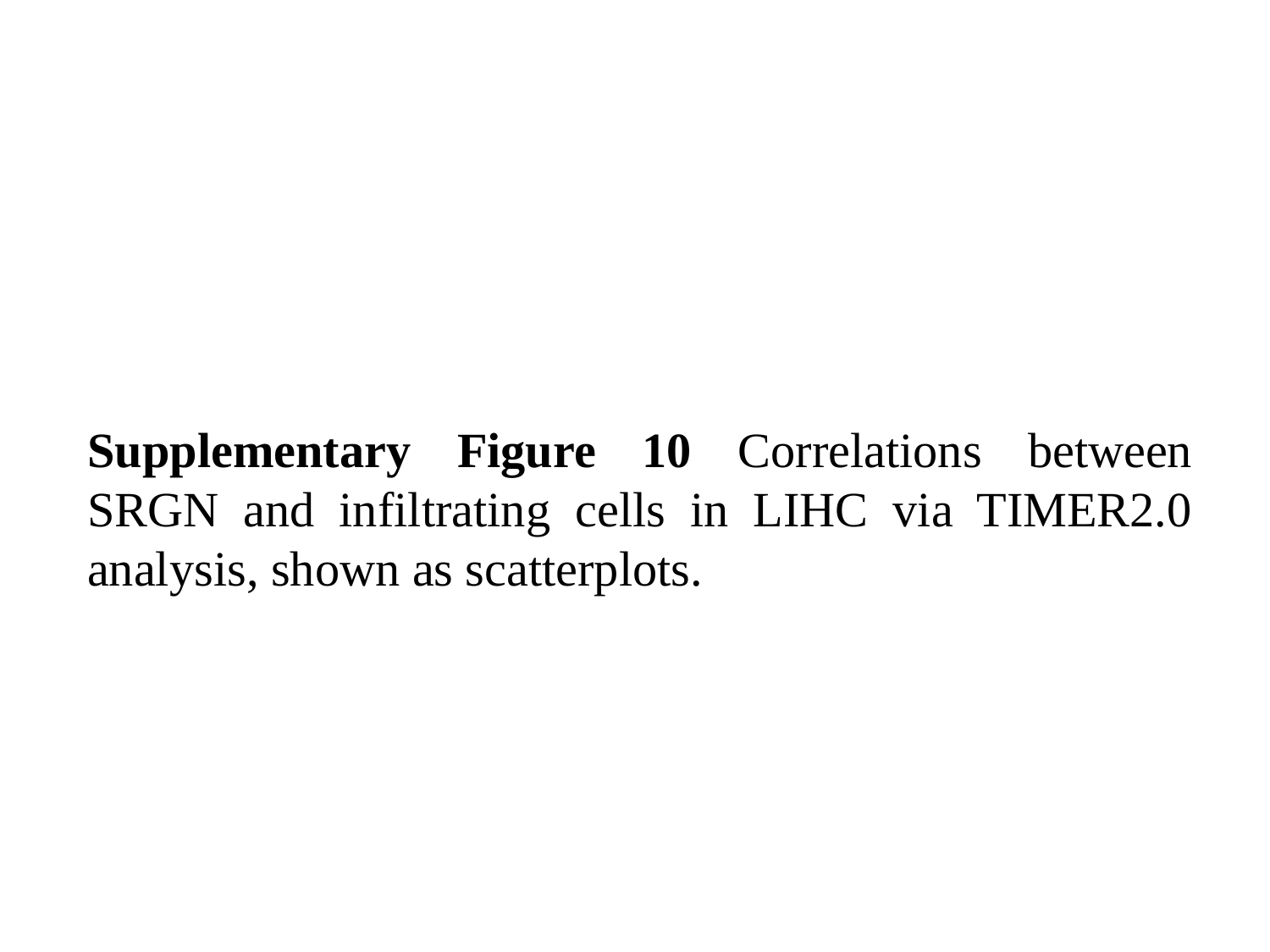

Supplementary Figure 10 Correlations between SRGN and infiltrating cells in LIHC via TIMER2.0 analysis, shown as scatterplots.

### Slide 10
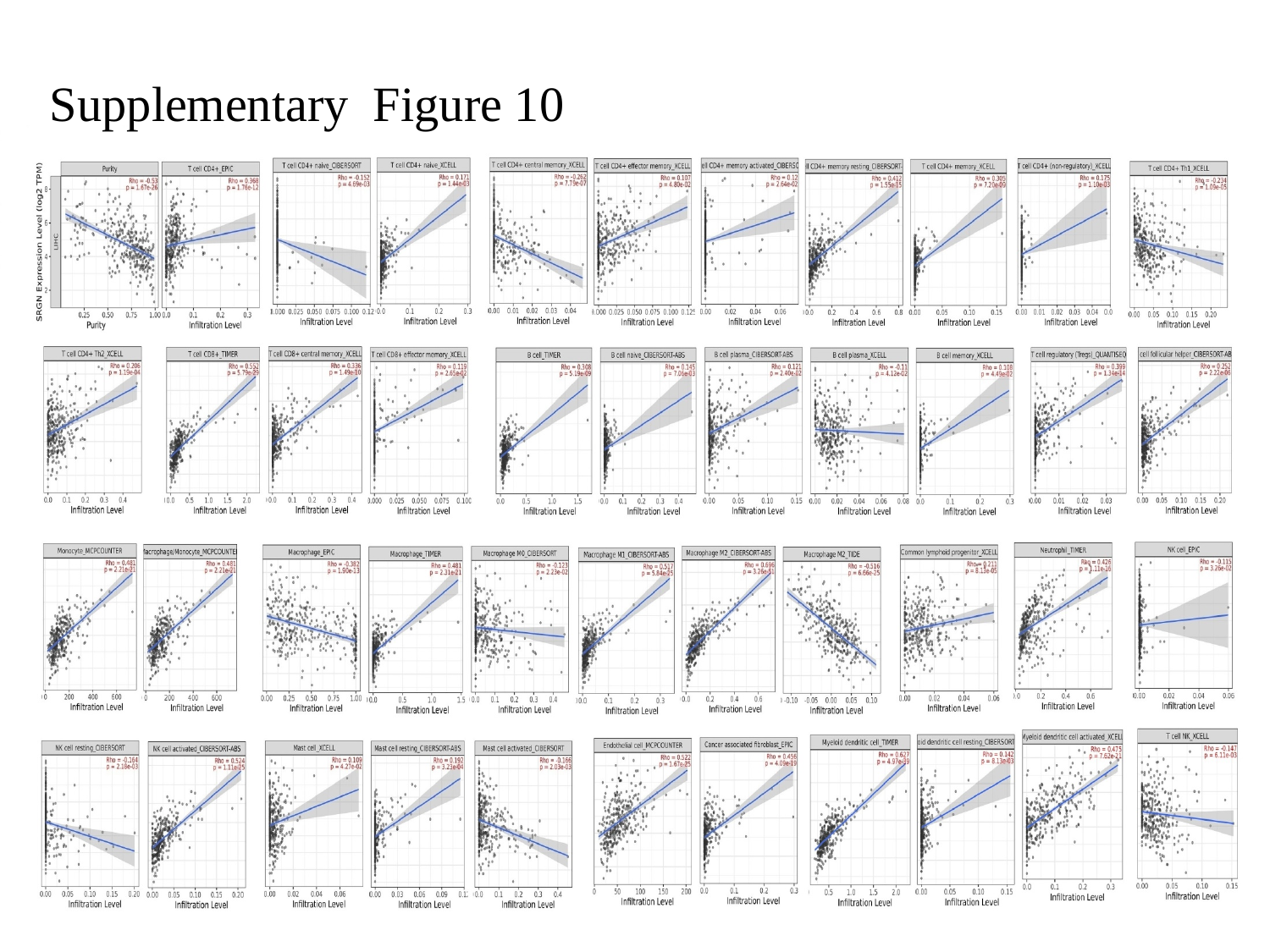

Supplementary Figure 10

### Slide 11
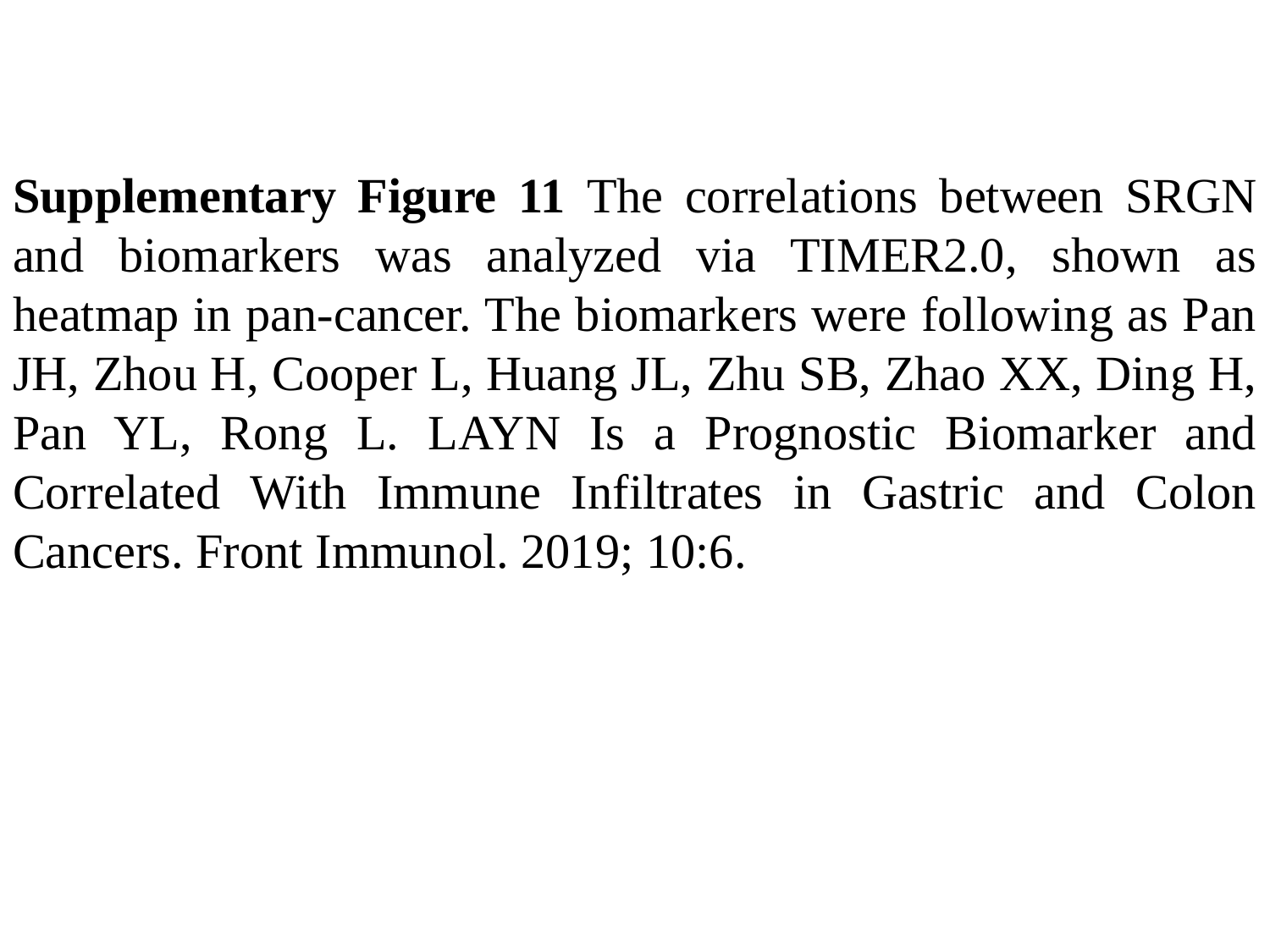

Supplementary Figure 11 The correlations between SRGN and biomarkers was analyzed via TIMER2.0, shown as heatmap in pan-cancer. The biomarkers were following as Pan JH, Zhou H, Cooper L, Huang JL, Zhu SB, Zhao XX, Ding H, Pan YL, Rong L. LAYN Is a Prognostic Biomarker and Correlated With Immune Infiltrates in Gastric and Colon Cancers. Front Immunol. 2019; 10:6.

### Slide 12
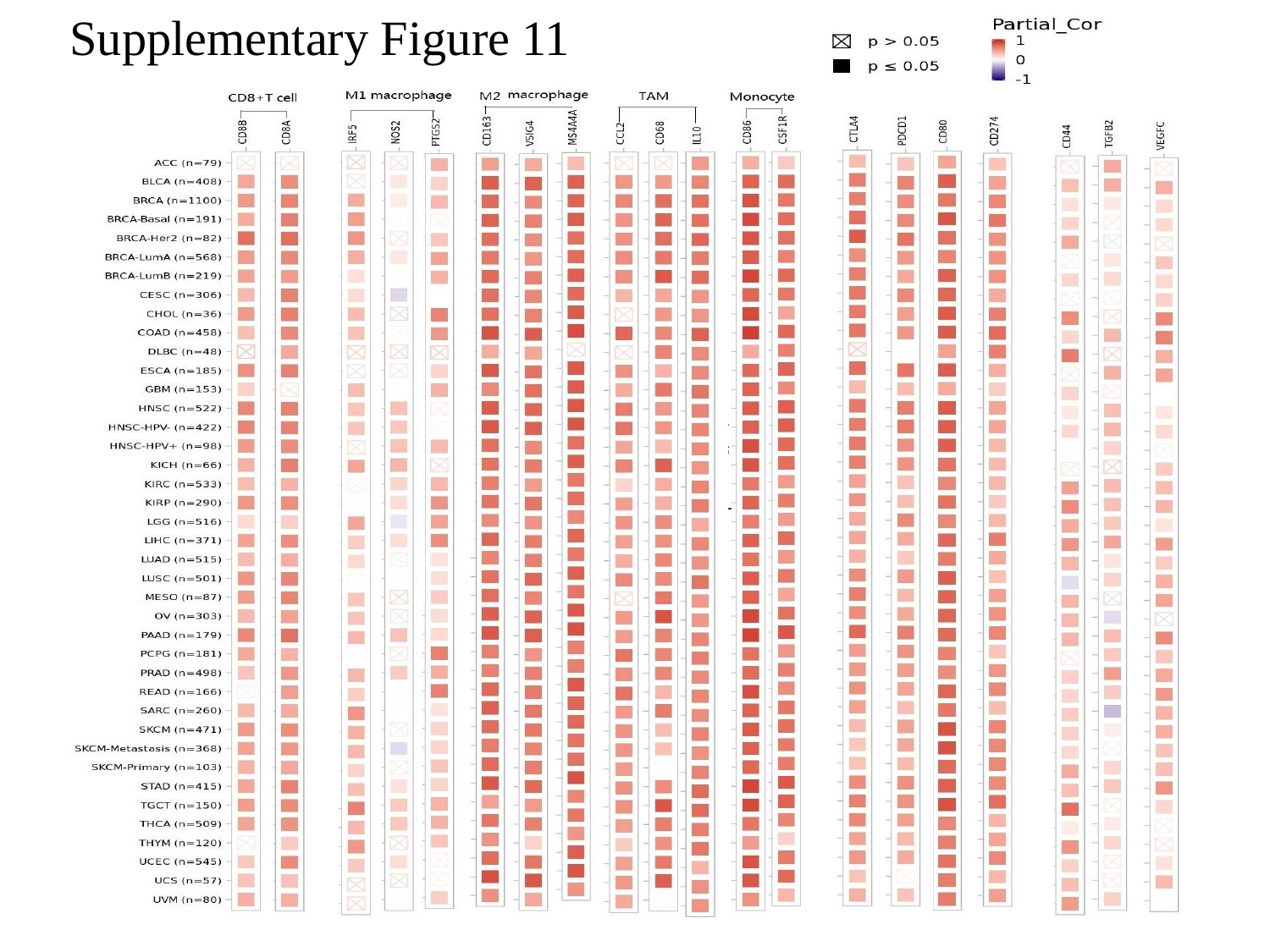

Supplementary Figure 11

### Slide 13
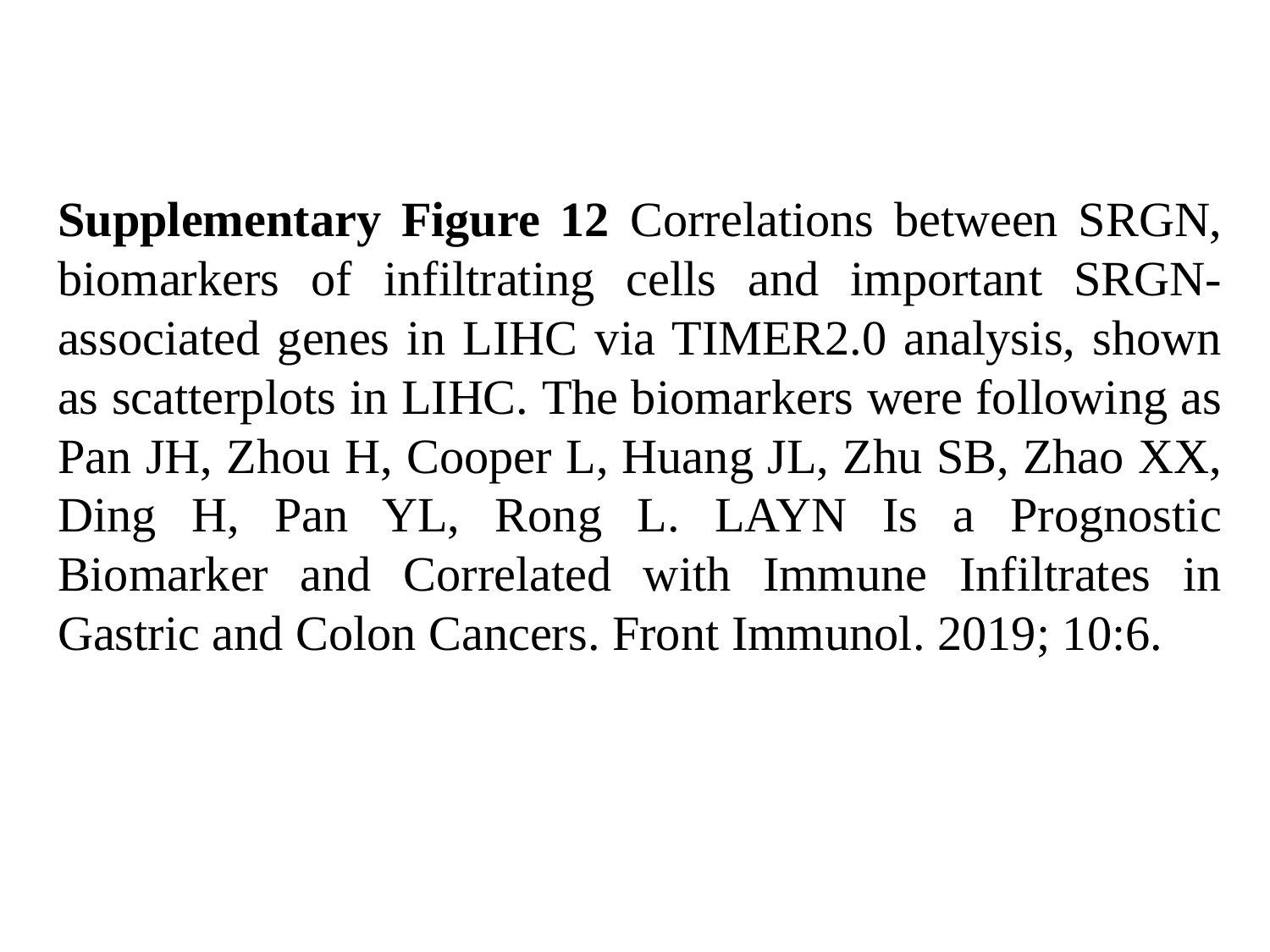

Supplementary Figure 12 Correlations between SRGN, biomarkers of infiltrating cells and important SRGN-associated genes in LIHC via TIMER2.0 analysis, shown as scatterplots in LIHC. The biomarkers were following as Pan JH, Zhou H, Cooper L, Huang JL, Zhu SB, Zhao XX, Ding H, Pan YL, Rong L. LAYN Is a Prognostic Biomarker and Correlated with Immune Infiltrates in Gastric and Colon Cancers. Front Immunol. 2019; 10:6.

### Slide 14
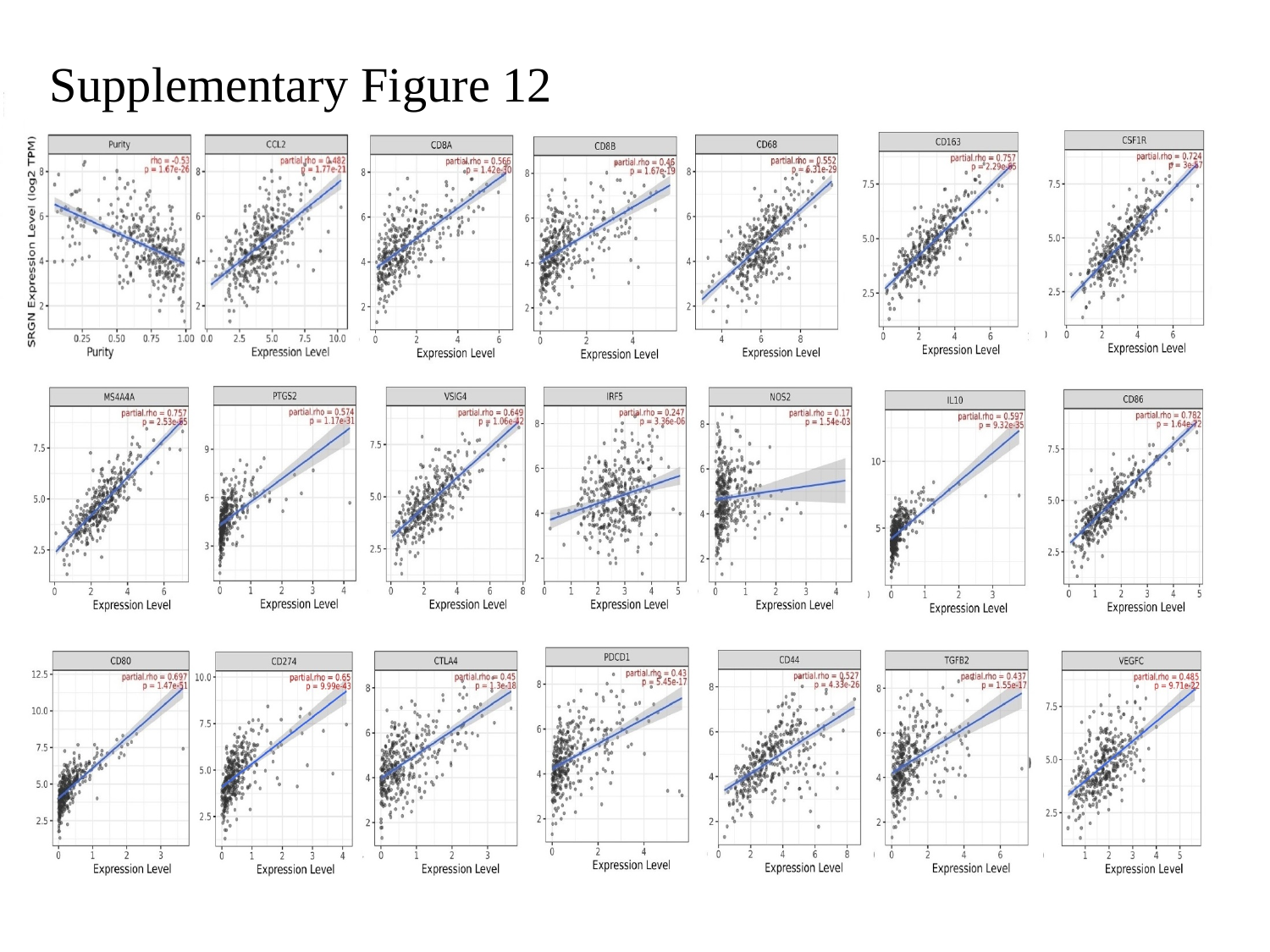

Supplementary Figure 12
