## Supplementary material for "Important Role of Hematopoietic Proteoglycan Serglycin in Liver Hepatocellular Carcinoma Associated with Tumor Microenvironment": supplemmental: Table.docx

.

Table1 The correlation of serglycin expression and prognosis in cancers via four bioinformatics databases

| Database and cancer | | ID | Patients | | Survival | | | Hazard ratio(HR) | | p-value |
| --- | --- | --- | --- | --- | --- | --- | --- | --- | --- | --- |
| PrognoScan | ID | | | Sample size | | Survival | HR(95%CI) | | Cox P | |
| AML | | 1554676_at  201859_at  201858_s_at  201859_at | 35/44  69/10  9/49  36/22 | | OS  OS  OS  OS | | | 1.35(1.03-1.76)  2.59( 1.01-6.63  1.99( 1.31-3.0)  2.67(1.48-4.82) | | 0.03  0.05  0.001  0.001 |
| DLBCL | | 201858_s_at  201858_s_at | 22/31  22/31 | | EFS  OS | | | 2.49(1.03-6.02)  2.59(1.02-6.56) | | 0.04  0.05 |
| Breast cancer | | 201859_at  201858_s_at | 86/29  91/195 | | DMSF  DMSF | | | 0.31(0.12-0.83) 0.77(0.62-0.96) | | 0.02  0.02 |
| Lung cancer | | 1554676_at | 182/22 | | RFS | | | 0.46(0.29-0.72) | | 0.001 |
| Ovarian cancer | | 201859_at | 87/46 | | OS | | | 0.82(0.70-0.97) | | 0.03 |
| GEPIA^#^ | | ID | Sample size | | Survival | | | HR(high) | | p-value |
| TGCT  GBM  SKCM  SKCM | | NA  NA  NA  NA | 68/68  80/81  228/229  228/229 | | OS  RFS  OS  RFS | | | 9.1e8  1.7  0.51  0.74 | | 0.022  0.013  5.2e-07  0.014 |
| TIMER2.0 | | ID | Sample size | | Survival | | | HR | | p-value |
| SARC  SKCM  SKCM metastasis  ACC  LGG | | NA  NA  NA  NA  NA | 260  471  368  79  516 | | OS  OS  OS  OS  OS | | | 0.762  0.746  0.734  1.49  1.19 | | 0.0213  0.000873  0.00184  0.0659  0.0691 |
| Kaplan-Meier plotter^*^ | | ID | Sample size | | Survival | | | HR(95%CI) | | p-value |
| Breast cancer | | 201859 _at  201859_at  201858_s_at  201858_s_at | 458  4929  1879  4929 | | PPS  RFS  OS  RFS | | | 1.39( 1.09-1.76)  1.15(1.03-1.27)  0.74(0.61-0.91)  1.13(1.01-1.27) | | 0.0066  0.012  0.0038  0.034 |
| Ovarian cancer | | 201858_s_at  201859_at  201859_at  201859_at | 1435  1435  1656  782 | | PFS  PFS  OS  PPS | | | 1.26,(1.11-1.45)  1.3(1.12-1.5)  1.15(1-1.33)  1.21(1-1.46) | | 0.00054  0.00038  0.051  0.05 |
| Lung cancer | | 201859_at  201859_at  201858_s_at | 982  1925  1925 | | FP  OS  OS | | | 0.58(0.48- 0.71)  0.78 (0.69 -0.89)  1.25(1.1-1.42) | | 9.9e-08  0.00022  0.00081 |
| Gastric cancer | | 201858_s_at  201858_s_at  201858_s_at  201859_at  201859_at  201859_at | 875  498  640  640  875  498 | | OS  PPS  FP  FP  OS  PPS | | | 0.55(0.46-0.66)  0.49(0.38-0.62)  0.57(0.47-0.7)  0.57(0.46-0.7）  0.58(0.49-0.69)  0.56(0.44-0.7) | | 6.3e-12  9.7e-10  4.8e-08  1e-07  2.1e-10  3.8e-07 |
| Liver cancer | | 5552  5552  5552  5552 | 366  364  313  357 | | PFS  OS  RFS  DSS | | | 0.67( 0.5-0.91)  0.68(0.47-0.97)  0.65((0.46-0.92)  0.58(0.35-0.97) | | 0.0095  0.034  0.015  0.034 |

The survival analysis of *SRGN* expression in pan-cancer. NA: the data was not shown; *Split patients by auto select best cutoff, other factors such as pathology, risk factors, patient, and follow up threshold, etc., were chosen all; ^#^group cutoff: median, cutoff -high/low: 50%. AML, acute myeloid leukemia ; DLBCL, diffuse large B-cell lymphoma ; LGG, brain lower grade glioma; ACC, adrenocortical carcinoma; SKCM, skin cutaneous melanoma; TGCT, testicular germ cell tumor; SARC, sarcoma; GBM, glioblastoma multiforme; OS, overall survival; PFS, progression free survival; RFS, relapse-free survival; PPS, post progression survival; FP, first progression; DSS, disease-specific survival; DMFS, distant metastasis-free survival; EFS, event free survival.

Table 2 Correlation of *SRGN* mRNA expression and prognosis in liver cancer with different clinical factors by Kaplan-Meier plotter.

| Clinical characteristics^*^ | N | OS (n= 364)  Hazard ratio | p-value | N | RFS(n=313)  Hazard ratio | p-value |
| --- | --- | --- | --- | --- | --- | --- |
| Male  Female | 246  118 | 0.56(0.36-0.88)  0.49(0.24-0.99) | **0.01**  **0.044** | 208  105 | 0.56(0.36-0.88)  0.56(0.28-1.1) | **0.0051**  0.085 |
| White  Asian | 181  155 | 0.58(0.36-0.93)  0.51(0.28-0.94) | **0.022**  **0.028** | 147  143 | 0.75(0.48-1.18)  0.55(0.33-0.92) | 0.21  **0.022** |
| Hepatitis virus(yes)  (no) | 150  167 | 0.72(0.38-1.37)  0.39(0.23-0.64) | 0.31  **0.00014** | 138  142 | 0.53(0.31-0.88)  0.65(0.38-1.12) | **0.014**  0.12 |
| Alcohol consumption(yes)  (no) | 115  202 | 0.35(0.18-0.7)  0.65(0.38-1.1) | **0.0017**  0.1 | 98  182 | 0.52(0.28-0.97)  0.63(0.38-1.02) | **0.037**  0.06 |
| Sorafenib treatment | 29 | 0.51(0.17-1.54) | 0.23 | 22 | 0.41(0.16-1.07) | 0.06 |
| Vascular invasion(none)  (micro) | 203  90 | 0.65(0.38-1.12)  2.14(0.74-6.22) | 0.12  0.15 | 175  81 | 0.74(0.45-1.21)  0.59(0.27-1.3) | 0.23  0.18 |
| AJCC_T(1)  (2)  (3) | 180  90  78 | 0.76(0.42-1.39)  0.44(0.21-0.95)  0.46(0.23-0.9) | 0.37  **0.031**  **0.021** | 160  79  65 | 1.27(0.72-2.24)  0.46(0.21-1.01)  0.56(0.27-1.15) | 0.4  **0.046**  0.11 |
| Grade (1)  2  3 | 55  174  118 | 0.44(0.17-1.14)  0.55(0.33-0.94)  1.3(0.69-2.46) | 0.084  **0.026**  0.42 | 45  147  106 | 1.74(0.64-4.71)  0.53(0.31-0.93)  0.54(0.31-0.93) | 0.27  **0.024**  **0.025** |
| Stage 1  1+2  2  2+3  3  3+4 | 170  253  83  166  83  87 | 0.73(0.39-1.38)  0.67(0.42-1.09)  0.45(0.2-1.01)  0.49(0.3-0.79)  0.41(0.21-0.8)  0.45(0.24-0.85) | 0.34  0.11  **0.046**  **0.0028**  **0.0071**  **0.011** | 153  227  74  142  68  68 | 1.55(0.87-2.76)  0.74(0.48-1.13)  0.51(0.25-1.05)  0.52(0.33-0.81)  0.54(0.29-1.01)  0.54(0.29-1.01) | 0.14  0.16  0.063  **0.0033**  0.051  0.051 |
| Clinical characteristics | N | PFS (n=366)  Hazard ratio | p-value | N | DSS(n=357)  Hazard ratio | p-value |
| Male  Female | 246  120 | 0.54（0.37-0.77）  0.56（0.31-1.01） | **0.00073**  0.051 | 241  116 | 0.53（0.29-0.97）  0.41（0.16-1.08） | **0.037**  0.062 |
| White  Asian | 183  155 | 0.73（0.49-1.07）  0.58（0.36-0.94） | 0.11  **0.025** | 177  152 | 0.62（0.35-1.11）  0.31（0.14-0.68） | 0.1  **0.002** |
| Hepatitis virus(yes)  (no) | 152  167 | 0.55（0.34-0.88）  0.64（0.39-1.03） | **0.011**  0.062 | 149  163 | 1.54（0.67-3.55）  0.37（0.2-0.69） | 0.31  **0.0013** |
| Alcohol consumption(yes)  (no) | 115  204 | 0.43（0.25-0.74）  0.68（0.43-1.08） | **0.002**  0.098 | 115  197 | 0.29（0.13-0.66）  0.63（0.31-1.29） | **0.0017**  0.2 |
| Sorafenib treatment | 30 | 0.37（0.15-0.9） | **0.023** | 29 | 0.51（0.17-1.54） | 0.23 |
| Vascular invasion(none)  (micro) | 204  91 | 0.84（0.54-1.3）  0.61（0.34-1.08） | 0.43  0.084 | 200  88 | 1.46（0.7-3.04）  4.62(0.6-35.68) | 0.31  0.11 |
| AJCC_T(1)  (2)  (3) | 180  92  78 | 1.58（0.95-2.63）  0.45（0.25-0.8）  0.46（0.24-0.88） | 0.073  **0.0049**  **0.016** | 177  89  75 | 3.82(0.9-16.26)  0.36(0.13-1.04)  0.4(0.18-0.89) | 0.051  **0.049**  **0.021** |
| Grade (1)  2  3 | 55  175  119 | 0.4（0.18-0.9）  0.6（0.37-0.96）  0.54（0.32-0.93） | **0.023**  **0.031**  **0.024** | 55  169  116 | 0.32(0.09-1.17)  0.48(0.24-0.95)  1.31(0.61-2.83) | 0.072  **0.032**  0.49 |
| Stage 1  1+2  2  2+3  3  3+4 | 170  254  84  167  83  88 | 1.62（0.96-2.74）  0.69（0.45-1.06）  0.44（0.24-0.82）  0.49（0.33-0.74）  0.45（0.24-0.82）  0.45（0.25-0.81） | 0.071  0.088  **0.0083**  **0.00043**  **0.0074**  **0.0066** | 167  249  82  163  81  84 | 3.18(0.74-13.74)  0.67(0.34-1.34)  0.33(0.1-1.08)  0.38(0.21-0.7)  0.36(0.17-0.78)  0.37(0.18-0.79) | 0.1  0.26  0.054  **0.0013**  **0.007**  **0.0075** |

^*^ The sample numbers with some clinical characteristics were too low for meaningful analysis, such as Black or Africa American, Grade 4, macro invasion, etc. N: sample size.

**Table3. The correlation between *SRGN* and infiltrating cells in LIHC by TIMER2.0**

| TME Cells | Rho | p-value |
| --- | --- | --- |
| T cell CD4+  T cell CD4+ naïve_CIBERSORT)  T cell CD4+ naïve_XCELL)  T cell CD4+ central memory  T cell CD4+ effector memory  T cell CD4+ memory activated  T cell CD4+ memory resting  T cell CD4+ memory  T cell CD4+(non-regulatory)  T cell CD4+ Th1  T cell CD4+ Th2 | 0.368  -0.152  0.171  -0.262  0.107  0.12  0.412  0.305  0.175  -0.234  0.206 | 1.76e-12  4.69e-03  1.44e-03  7.79e-07  4.8e-02  2.64e-02  1.55e-15  7.2e-09  1.1e-03  1.09e05  1.19e-04 |
| T cell CD8+  T cell CD8+ central memory  T cell CD8+ effector memory | **0.552**  0.336  0.119 | **5.79e-29**  1.49e-10  2.65e-02 |
| B cell  B cell naïve  B cell plasma_CIBERSORT-ABS  B cell plasma_XCELL  B cell memory | 0.308  0.145  0.121  -0.11  0.108 | 5.19e-09  7.06e-03  2.4e-02  4.12e-02  4.49e-02 |
| T cell regulatory | 0.399 | 1.34e**-**14 |
| T cell follicular helper | 0.252 | 2.22e-06 |
| Monocyte  Macrophage/monocyte  Macrophage_EPIC  Macrophage_TIMER  Macrophage M0  Macrophage M1  Macrophage M2_CIBERSORT-ABS  Macrophage M2_TIDE | 0.481  0.481  -0.382  0.481  -0.123  **0.517**  **0.696**  **-0.516** | 2.21e-21  2.21e-21  1.9e-13  2.31e-21  2.23e-02  **5.84e-25**  **3.26e-51**  **6.66e-25** |
| Myloid dendritic cell  Myloid dendritic cell activated  Myloid dendritic cell resting | **0.627**  0.475  0.142 | **4.97e-39**  7.62e-21  8.13e-03 |
| Common lymphoid progenitor | 0.211 | 8.13e-05 |
| Neutrophil | 0.426 | 1.11e-16 |
| NK cell  NK cell resting  NK cell activated | -0.115  -0.164  **0.524** | 3.26e-02  2.18e-03  **1.11e-25** |
| T cell NK | -0.147 | 6.11e-03 |
| Mast cell  Mast cell resting  Mast cell activated | 0.109  0.192  -0.166 | 4.27e-02  3.23e-04  2.03e03 |
| Endothelial cell | **0.522** | **1.67e-25** |
| Cancer associated fibroblast | 0.456 | 4.09e-19 |

**Table4. The survival analysis of *SRGN* expression associated with infiltrating cells in LIHC by TIMER2.0**

| Cells in TME | Low SRGN  expression  Hazard ratio | | p-value | | High SRGN  expression  Hazard ratio | | p-value | |
| --- | --- | --- | --- | --- | --- | --- | --- | --- |
| T cell CD8+  T cell CD4+ memory resting  Hematopoietic stem cell  MDSC  Endothelial cell  Monocyte(MCPcounter) | | 0.658  0.847  0.834  1.41  0.565  1.65 | | 0.139  0.584  0.551  0.313  0.0566  0.105 | | 0.53  0.501  0.379  2.78  0.364  2.75 | | **0.0448**  **0.0383**  **0.00868**  **0.00668**  **0.00638**  **0.025** |
| Macrophage  Macrophage M1  Macrophage 0  Macrophage 2 | | 1.82  1.19  1.86  1.99 | | **0.0432**  0.558  **0.0356**  **0.0366** | | 1.58  3.09  1.59  1.25 | | 0.198  **0.00633**  0.159  0.568 |
| T cell CD4+ central memory  T cell CD4+ Th2  Tregs  Monocyte(XCELL)  Myloid dendrtic cell  Myloid dendrtic cell resting | | 0.742  1.22  0.873  1.19  0.924  1.34 | | 0.305  0.498  0.659  0.599  0.796  0.283 | | 0.874  1.83  0.671  1.95  3.28  1.05 | | 0.694  0.0945  0.249  0.0811  0.0599  0.871 |

TME: tumor environment
