## Supplementary figures and images for "Important Role of Hematopoietic Proteoglycan Serglycin in Liver Hepatocellular Carcinoma Associated with Tumor Microenvironment"

### Figure 1 DiffExp of SRGN in pan-cancer by TIMER and CPTAC.jpg

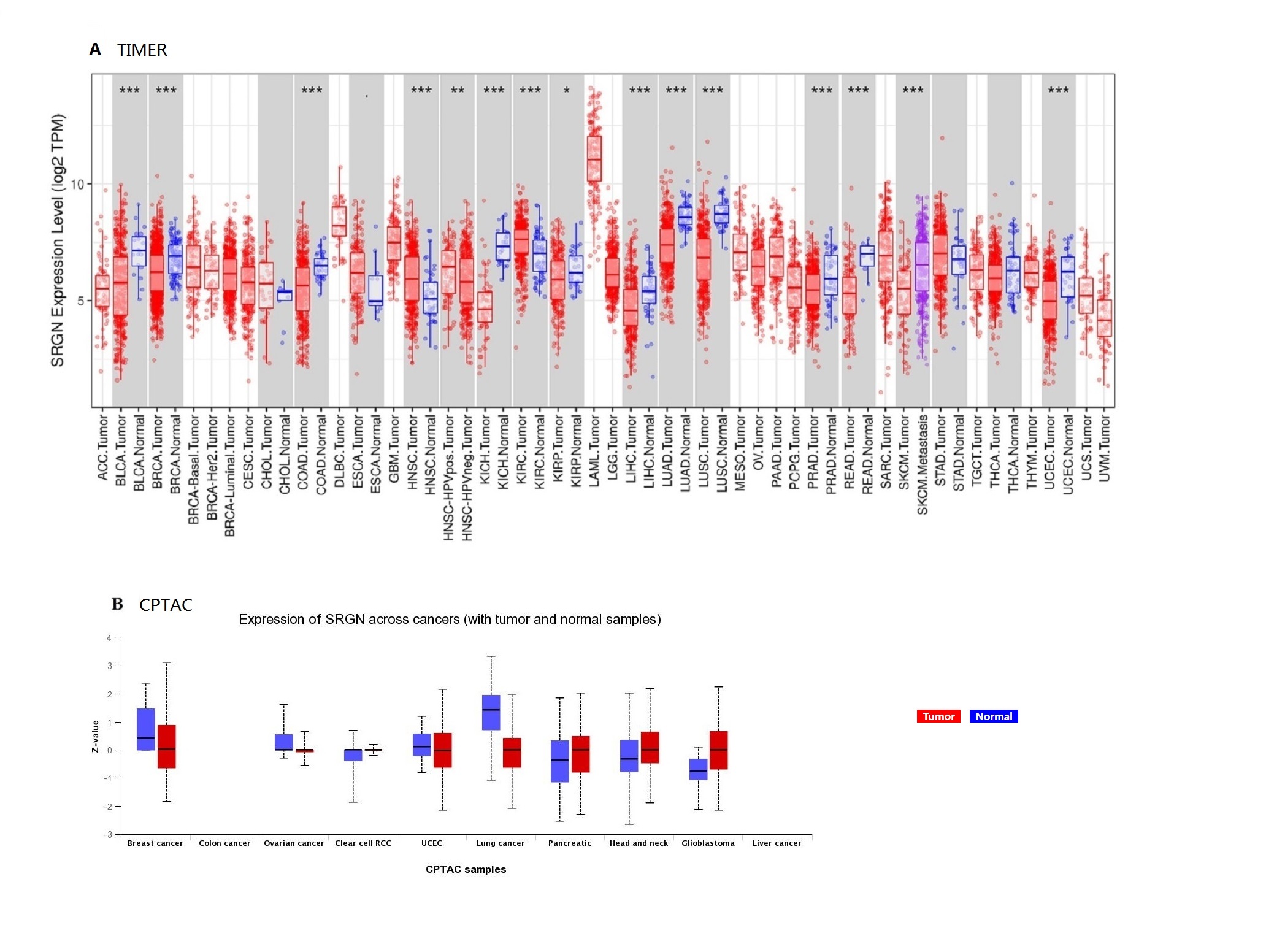

### Figure 2 SRGN expression and cell type.jpg

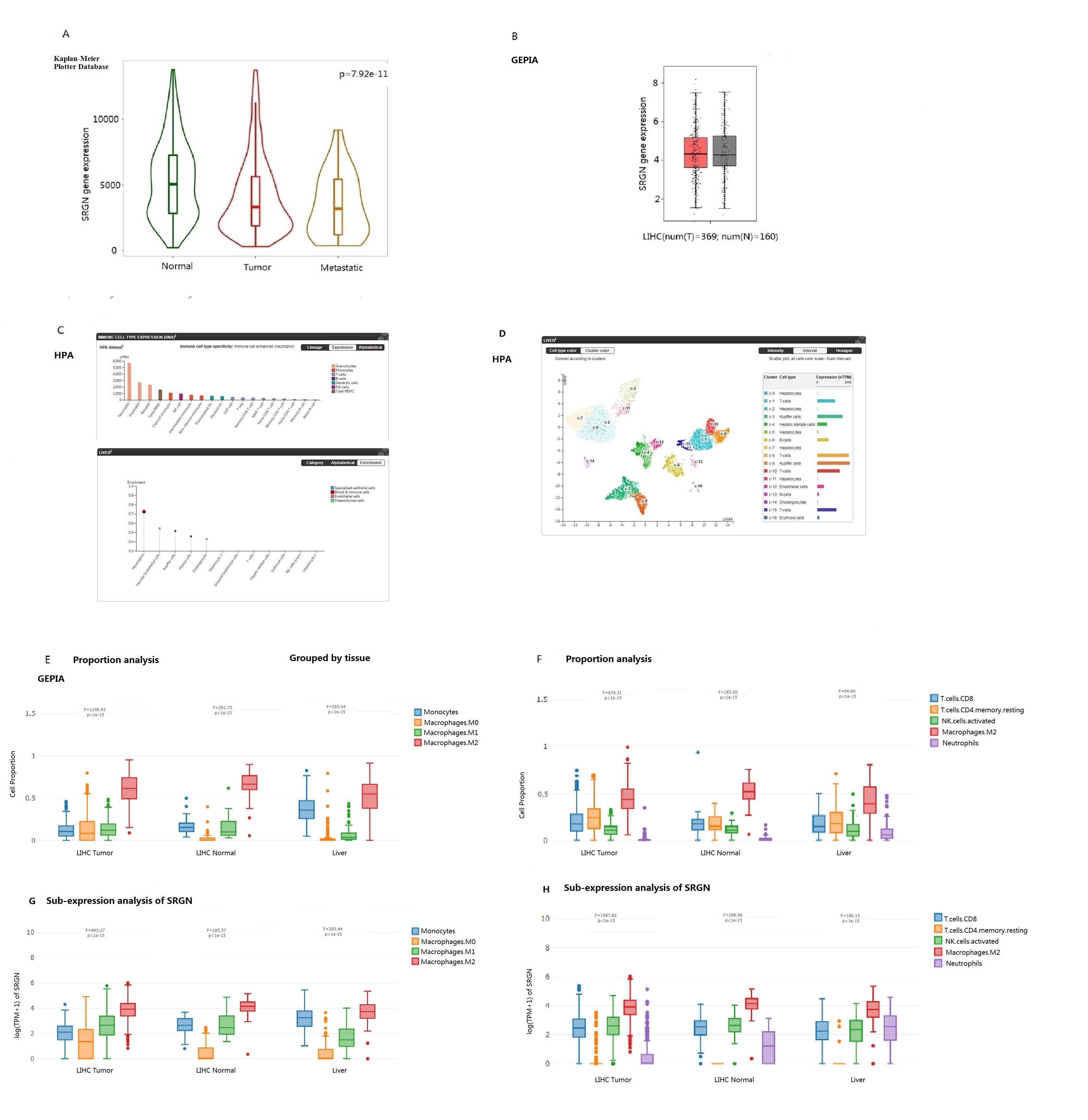

### Figure 3 Correlations between SRGN and markers.jpeg

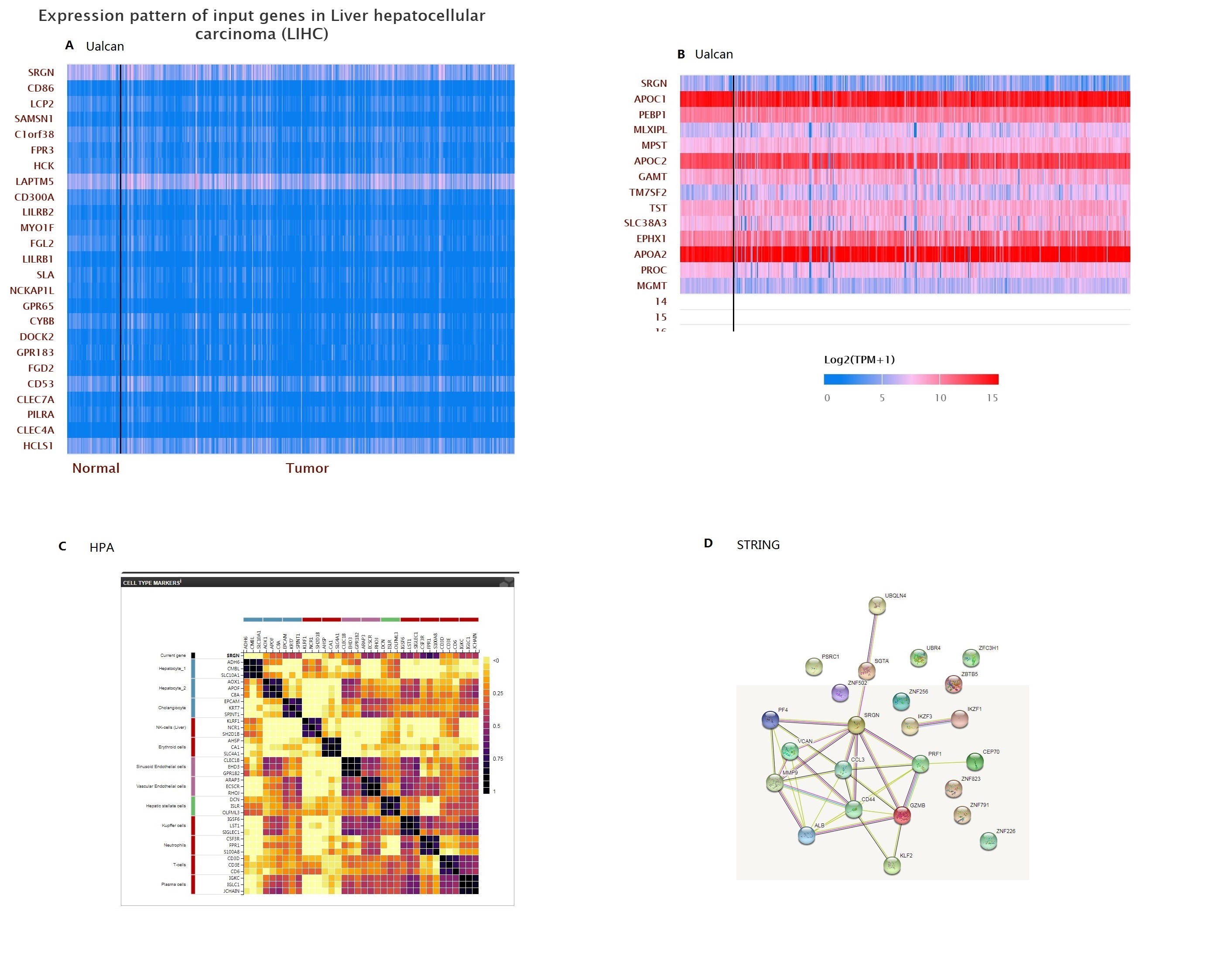

### Figure 4 the KM of SRGN and TME cells by TIMER2.0.jpg

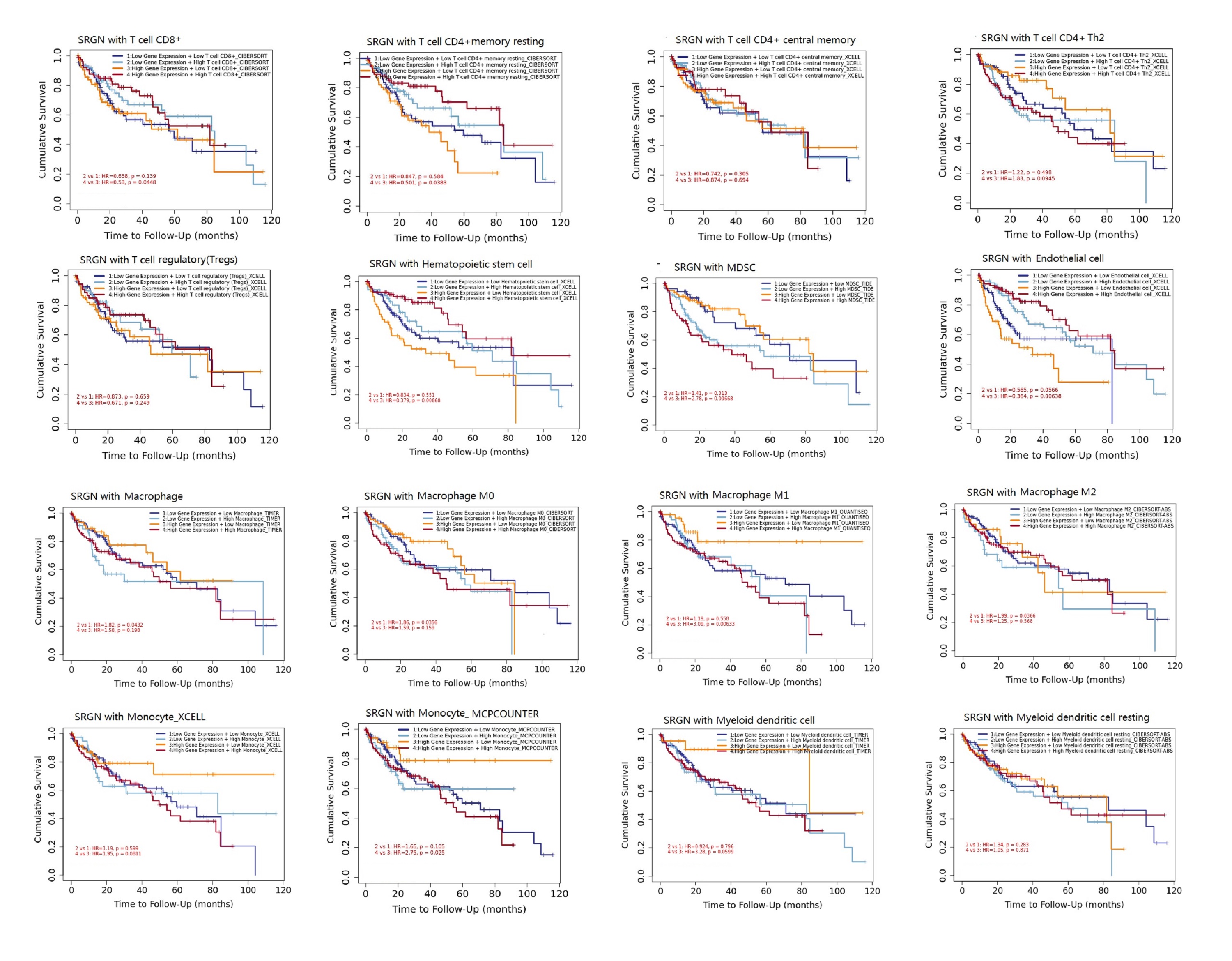

### Figure 5 in vitro and in vivo experiment.jpg

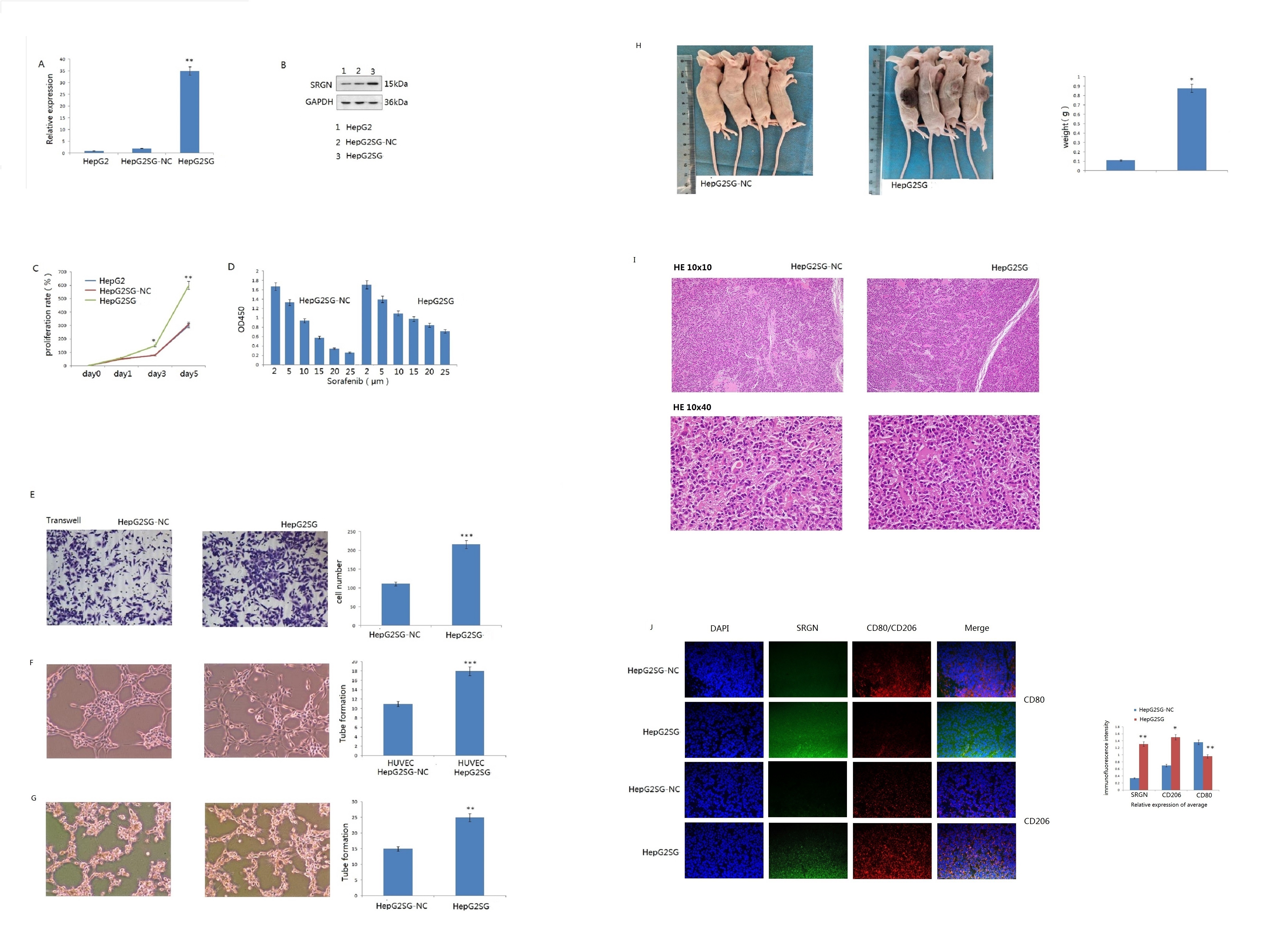
